# Long-term *in vivo* three-photon imaging reveals region-specific differences in healthy and regenerative oligodendrogenesis

**DOI:** 10.1101/2023.10.29.564636

**Authors:** Michael A. Thornton, Gregory L. Futia, Michael E. Stockton, Samuel A. Budoff, Alexandra N Ramirez, Baris Ozbay, Omer Tzang, Karl Kilborn, Alon Poleg-Polsky, Diego Restrepo, Emily A. Gibson, Ethan G. Hughes

## Abstract

The generation of new myelin-forming oligodendrocytes in the adult CNS is critical for cognitive function and regeneration following injury. Oligodendrogenesis varies between gray and white matter regions suggesting that local cues drive regional differences in myelination and the capacity for regeneration. Yet, the determination of regional variability in oligodendrocyte cell behavior is limited by the inability to monitor the dynamics of oligodendrocytes and their transcriptional subpopulations in white matter of the living brain. Here, we harnessed the superior imaging depth of three-photon microscopy to permit long-term, longitudinal *in vivo* three-photon imaging of an entire cortical column and underlying subcortical white matter without cellular damage or reactivity. Using this approach, we found that the white matter generated substantially more new oligodendrocytes per volume compared to the gray matter, yet the rate of population growth was proportionally higher in the gray matter. Following demyelination, the white matter had an enhanced population growth that resulted in higher oligodendrocyte replacement compared to the gray matter. Finally, deep cortical layers had pronounced deficits in regenerative oligodendrogenesis and restoration of the MOL5/6-positive oligodendrocyte subpopulation following demyelinating injury. Together, our findings demonstrate that regional microenvironments regulate oligodendrocyte population dynamics and heterogeneity in the healthy and diseased brain.

---

In the central nervous system (CNS), oligodendrocytes produce myelin, which enwraps neuronal axons to increase axonal conduction, fine-tune circuit timing, and provide metabolic support. In 1901, Paul Flechsig proposed that regional myelination is regulated by the functional maturation of neural circuits (fundamental law of myelogenesis^1^), implying that the cellular microenvironment may shape the generation of oligodendrocytes.

Throughout life, oligodendrocytes are generated via the differentiation of oligodendrocyte precursor cells (OPCs, also called oligodendrocyte progenitor cells). Recent work shows that environmental factors such as brain region and age influence OPCs to become functionally heterogenous via differential ion channel expression^2^. In line with these findings, the rates of OPC proliferation and differentiation differ between gray and white matter areas^3,4^ and white matter-derived precursors generate oligodendrocytes with a higher efficiency than gray matter-derived OPCs after transplantation^5^. Additionally, mature oligodendrocytes make up a heterogeneous cell population with distinct transcriptional subtypes^6^ that exhibit spatial preference^7,8^ and differential responses to injury and disease^9–11^. However, a complete understanding of region-specific regulation of oligodendrogenesis and transcriptional heterogeneity is limited due to the inability to monitor the population dynamics of oligodendrocytes in white matter in the living brain.

In demyelinating diseases, such as multiple sclerosis (MS), death of oligodendrocytes elicits regenerative oligodendrogenesis^12^ and remyelination that can restore neuronal function^13,14^. Historically, MS has been regarded as a disease of the white matter; however, recent work shows that MS patients have extensive gray matter lesions that correlate with late-stage disabilities^15,16^. Interestingly, leukocortical lesions that span white and gray matter regions display distinct pathological hallmarks suggesting that cellular microenvironments of lesions in different brain regions distinctly regulate myelin loss and repair^17^. Yet, the cellular dynamics of oligodendrocyte population loss and regeneration following demyelination remain uncharacterized across brain regions.

Advancements in multiphoton microscopy have revolutionized our understanding of cellular dynamics in intact tissues by extending imaging depths four-fold compared to conventional confocal microscopy^18^. For example, previous studies using longitudinal *in vivo* two-photon imaging in mouse primary motor cortex^14^ and somatosensory cortex^12,19,20^ revealed fundamental aspects of the dynamics of oligodendrocytes in the healthy and demyelinated brain. However, due to light scattering in brain tissue these studies have been limited to the superficial ∼400 microns of the cortical gray matter when extended over multiple months. The recent development and application of three-photon microscopy, which uses a longer excitation wavelength and experiences reduced scattering compared to two-photon, has permitted *in vivo* imaging depths greater than 1 mm at high resolution^21–23^. However, the high peak intensity of excitation pulses required for nonlinear excitation can result in tissue damage particularly when applied repetitively^24^. Whether three-photon microscopy can be reliably utilized for long-term volumetric measurements of cellular behavior in the intact brain remains unclear.

Here, we asked whether the dynamics of oligodendrocyte populations differ across cortical and subcortical regions in the context of health and disease. To address this question, we developed an imaging protocol that permits long-term volumetric *in vivo* three-photon imaging without tissue damage, cellular reactivity, or perturbation of healthy oligodendrogenesis in the adult mouse brain. This protocol enabled us to track the cellular behavior of over 13,000 individual oligodendrocytes across 2-3 months in health and disease. Long-term *in vivo* three-photon imaging showed that white matter generated more oligodendrocytes per volume compared to gray matter, however, the population growth of oligodendrocytes was accelerated in the gray matter. Following cuprizone-mediated demyelination, we found that the oligodendrocytes and their subpopulations were equivalently reduced across gray and white matter, however, the white matter showed enhanced regenerative oligodendrogenesis and population replacement compared to gray matter regions. Layer-specific analyses revealed that the deep cortical layers had a reduced capacity for regenerative oligodendrogenesis and restoration of the MOL5/6 oligodendrocyte subpopulation following demyelinating injury. Together, these results indicate that regional microenvironments differentially modulate oligodendrogenesis and heterogeneous oligodendrocyte subpopulations to influence myelination and regeneration in the healthy and diseased brain.

## Results

### In vivo three-photon imaging of cortical gray and white matter

To determine region-specific dynamics of oligodendrocyte generation in an entire cortical column encompassing the six layers of the cerebral cortex and the subcortical white matter, we built a custom three-photon microscope that enabled imaging to depths of ∼1000 µm across multiple months (**Methods, Extended Data Fig. 1**). We implanted chronic cranial windows over the posterior parietal cortex (PPC) of *MOBP-EGFP* transgenic mice^25^ where enhanced green fluorescent protein (EGFP) expression is driven by the MOBP promoter, thereby labeling all mature oligodendrocytes and their associated myelin sheaths in both the gray and white matter (**Extended Data Fig. 2**). To determine the efficacy of three-photon microscopy to visualize oligodendrocytes across cortical gray and white matter regions, we compared two-photon and three-photon imaging modalities. Two-photon imaging of the highest clarity cranial windows in *MOBP-EGFP* transgenic mice three weeks after implantation allowed the visualization of oligodendrocyte cell bodies to ∼600 µm from the brain surface (**Fig. 1a**). Three-photon imaging allowed for the collection of ∼400 x 400 x 1050 µm 3D volumes of EGFP-labeled oligodendrocytes encompassing the entire cortical column as well as subcortical white matter in chronic cranial windows of a wide range of clarity (**Fig. 1b, Supplementary Video 1**). Because multiphoton imaging quality is dependent on a combination of signal generation and out-of-focus background, we used the signal-to-background ratio (SBR) to compare two-photon and three-photon image quality in *MOBP-EGFP* mice. We found that two-photon imaging had higher SBR of oligodendrocyte cell bodies compared to three-photon imaging in the superficial 100 µm of cortex. This enhanced SBR is likely due to the increased probability of two-photon excitation in regions of minimal scattering^24^. In contrast, we found that three-photon imaging had increased SBR at depths greater than 400 µm beneath the brain surface compared to two-photon imaging (21.8±3.9 vs. 7.3±3.1 for 401 – 500 µm depth, **Fig. 1c**). This increase in three-photon image quality resulted in the detection of a higher number of oligodendrocytes in layer 5 when compared with two-photon data (78.8±10.2 vs. 41.4±10.7, **Fig. 1d**), highlighting that two-photon imaging only partially samples the labeled oligodendrocyte population at depths below superficial cortical layers 1-4.

**Figure 1.**
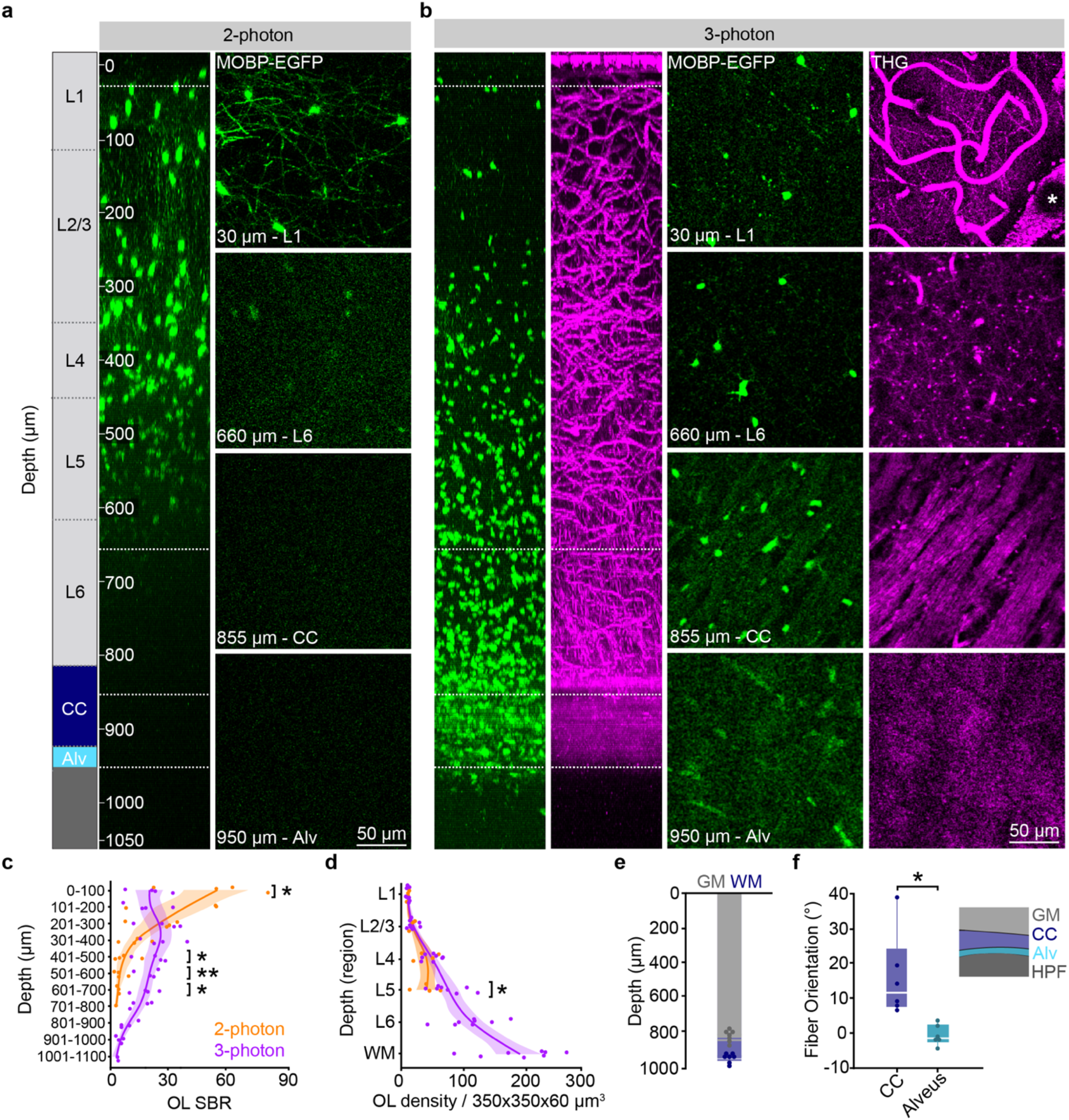
Three-photon microscopy enhances the detection of oligodendrocytes and myelin deep in the adult mouse brain. **a**) xz projection of a 2-photon z stack of oligodendrocytes and myelin in a high-quality chronic cranial window acquired with the Olympus 25X objective. Power was modulated from 8-160 mW. **b**) xz projection of a 3-photon z stack of *MOBP-EGFP* and third harmonic generation (THG) signal shows high SBR through the PPC, subcortical corpus callosum (CC), and the alveus of the hippocampus. The average power at the sample was modulated from 0 – 45 mW. **c**) Two-photon (2P) SBR of oligodendrocyte cell bodies is significantly increased in the superficial 100 µm of cortex compared to three-photon (3P, unpaired, two-tailed Student’s t-test, t(6) = -2.45, p = 0.043), while 3P imaging at depths greater than 400 µm has enhanced SBR (unpaired, two-tailed Student’s t-test, t(6) = 2.92, p = 0.027 (401 - 500 µm), t(6) = 4.72, p = 0.003 (501 - 600 µm), t(3.07) = 4.41, p = 0.021 (601 - 700 µm). **d**) 2P and 3P allow for the detection of similar numbers of oligodendrocytes in layers 1-4, yet significantly fewer cells are detected with 2P in layer 5 (unpaired, two-tailed t-test, t(10) = 2.29, p = 0.045. **e**) Stacked bar chart showing the cortical (gray) and total (blue) imaging depth across the PPC for n = 8 mice at 10 weeks of age. **f**) The corpus callosum (CC) can be differentiated from the alveus by the orientation of THG-positive fibers (t(5.57) = -3.24, p = 0.026) for n = 6 mice. *p < 0.05, **p < 0.01, n.s., not significant; cumulative growth curves represent cubic splines with 95% confidence intervals, bar plots represent mean +/- SEM, box plots represent the median, interquartile ranges and minimum/maximum values. For detailed statistics, see **Supplementary Table 3**.

Three-photon microscopy allowed for the collection of third harmonic generation (THG) signal, which is generated at optical found that repeated, long-term three-photon volumetric imaging using these parameters caused cumulative tissue damage that was visualized using THG imaging (**Fig. 2a**). This laser-induced damage resulted in oligodendrocyte cell death and a subsequent burst of oligodendrogenesis that could be predicted by a two-fold increase in the THG intensity at the previous time point (**Fig. 2b-c**). The onset of diffuse 1300 nm excitation-induced oligodendrocyte cell death was more rapid than reported for targeted single-oligodendrocyte ablation with 775 nm excitation^32^, and was sustained for up to fifty days. The injury-induced oligodendrogenesis had variable dynamics with a peak rate at ∼20-40 days post-lesion (**Fig. 2b,c**). To develop an imaging protocol appropriate for multi-month longitudinal *in vivo* imaging of cell behavior, we implemented several optical and scanning modifications (**Extended Data Fig. 4**). First, to reduce optical aberrations and increase image quality, we used dual-axis goniometers to align the chronic cranial window to be perpendicular to the light path^33^. Second, the excitation laser was blanked on the galvonometer over-scan to reduce the average power applied to the brain to < 52 mW (**Extended Data Fig. 4**). Third, frame averaging was utilized rather than pixel averaging or extended dwell time techniques, and the acquisition was paused for one minute per every three minutes of continuous scanning Fourth, we used adaptive optics optimized on the THG signal just above the corpus callosum to enhance the SBR of deep interfaces of different refractive index and third-order nonlinear susceptibility^26,27^ (**Supplementary Video 1**). Similar to previous studies^21,28^, we found that THG strongly labeled blood vessels from venules to capillaries, individual myelin sheaths, and myelinated fiber bundles in the subcortical white matter (**Fig. 1b**). We confirmed that THG-positive sheaths in the superficial PPC colocalized with EGFP+ sheaths in *MOBP-EGFP* transgenic mice and revealed a similar *in vivo* myelin labeling pattern to Spectral Confocal Reflectance (SCoRe) microscopy^29^ (**Extended Data Fig. 2**). Adaptive optics (AO) via image-based Zernike-mode aberration correction with a deformable mirror revealed individual THG-positive myelin sheath cross sections at depths of up to 1000 µm and increased the SBR and axial resolution of oligodendrocytes in the corpus callosum (**Extended Data Fig. 3**). At depths of 750 – 900 µm, AO correction resulted in a 68% increase in the normalized 3P signal of oligodendrocyte cell bodies and a 202% increase in oligodendrocyte processes. Furthermore, AO optimization allowed for a ∼56% reduction in the average power applied to the sample at depth (**Extended Data Fig. 3**). Additionally, we used THG microscopy to determine the depth from cortical surface to white matter and to further define anatomical landmarks within the white matter. Across eight, ten-week-old mice, we found that the average distance from brain surface to white matter in the PPC was 823±16 µm, while the thickness of the white matter was 118±8 µm, for a total imaging depth of 941±11 µm (**Fig. 1e**). The subcortical white matter below the posterior parietal cortex consists of myelinated fiber tracts of output axons from the cortex (corpus callosum) and the hippocampus (alveus)^30^ (**Fig. 1f**). We found that the orientation of the THG+ fibers allowed for the delineation of the corpus callosum and the alveus of the hippocampal formation (15.3±4.9 vs. -0.4±1.04 degrees, respectively, **Fig. 1b,f**). Together, these data show that three-photon imaging provides greater imaging depth, higher imaging quality, and an additional imaging modality to study oligodendrocytes and myelination in cortical gray and white matter regions.

**Figure 2.**
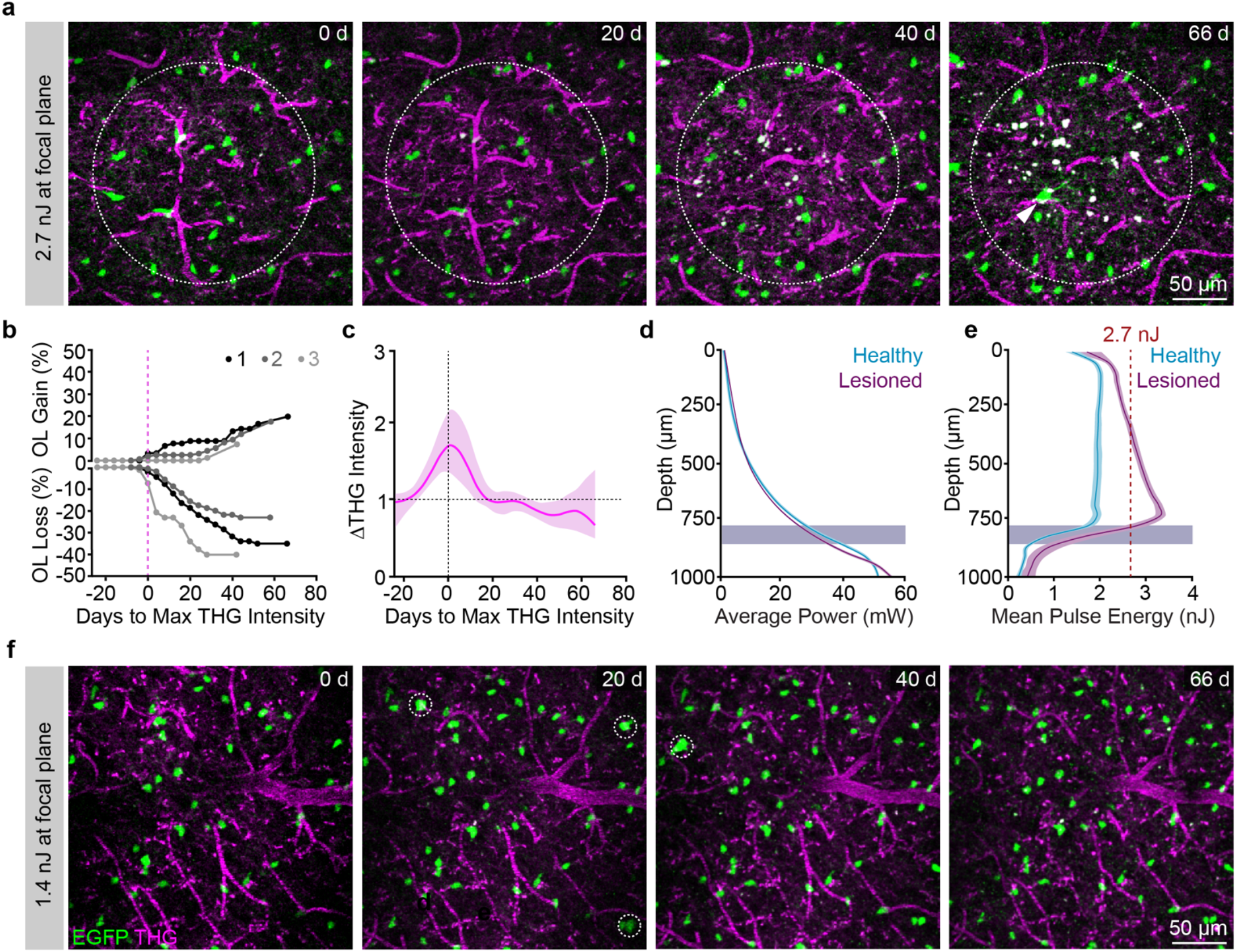
Optimization of excitation parameters permits long-term three-photon microscopy without tissue damage. **a**) Example imaging time points from a mouse exposed to 2.7 nJ pulse energy at the focus. Dotted circle represents the area of the lesion as defined by an increase in THG intensity; white arrowhead shows a newly-generated OL in response to cell ablation. Depth of lesion = 594 – 786 µm. **b**) Cumulative OL population cell loss (%) and cell gain (%) in lesioned regions shows rapid OL cell death that proceeded for up to 40 days following the time point of max THG intensity increase. OL cell regeneration was rapid and biphasic for lesion 1 (black) but delayed by ∼20 days and linear for lesions 2 and 3. Mean OL population loss was 32.9 +/- 5.2 %; Mean replacement of the lost OLs was 50.0 +/- 16.9 % (mean +/- SEM). **c**) Mean intensity of the THG signal over time in the lesion area (dotted circle), aligned to the time point of max intensity change for n=3 mice lesioned mice. **d**) Similar exponential power vs. depth curves were applied at all time points in lesioned vs. healthy mice that underwent long-term 3P imaging. The excitation laser was blanked on the galvanometer overscan. Data are from n=3 mice with laser-induced injury and n= 11 mice with successful longitudinal imaging without THG damage or cell death. **e**) The pulse energy at the focus vs. z-depth was determined for each mouse by calculating mouse-specific effective attenuation lengths (EALs) at the first time point. Mouse-specific differences in cortical EAL, and not average power (**d**), drove the increase in pulse energy that caused cellular damage. The minimum damage threshold was 2.7 nJ at the focus (red dotted line). Healthy imaged mice had a mean pulse energy at the focus of less than 2 nJ across the cortical depth (cyan). **f**) Example imaging time points from a healthy mouse with optimized settings and ∼1.4 nJ excitation at the focus across the cortical depth. Note the detection of newly generated oligodendrocytes (white dotted circles) without cell death or changes in THG intensity. THG, power, and pulse energy curves are cubic splines with 95% confidence intervals. Mean z-width of the lesions was 149 +/- 26.1 µm.

### Optimization of excitation parameters for long-term three-photon microscopy without tissue damage

Although previously published three-photon average power guidelines^31^ allowed for successful acquisition of single time-point three-dimensional volumetric and continuous single-plane time lapse imaging (**Fig. 1**), we oligodendrocytes (**Extended Data Fig. 4**), and linearly modulated the spherical correction with depth to maintain SBR in more superficial regions. Due to mouse-to-mouse variability in the scattering properties of the intact brain, nearly identical exponential power curves resulted in either healthy long-term imaging or lesions with oligodendrocyte cell death (**Fig. 2d**). Therefore, we measured mouse-specific effective attenuation lengths (EALs) in the adult posterior parietal cortex to estimate the pulse energy delivered at each focal plane as described previously^21,28,34^ (**Extended Data Figure 5**; **Methods**). The pulse energy at the focal plane was calculated using the equation below:

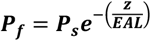

Where ***P***_*f*_ is pulse energy at the focal plane (nJ), ***P***_*s*_ is pulse energy at the surface (nJ), Z is depth from the brain surface (µm), EAL is the measured effective attenuation lengths (gray and white matter) for each mouse. Maintaining a relatively constant pulse energy of ∼2 nJ across the cortical depth (1.98±0.20 nJ; n = 11 longitudinally-imaged mice) with an absolute maximum threshold of 2.7 nJ (**Fig. 2e**) was sufficient to maintain high oligodendrocyte SBR and track healthy oligodendrogenesis without oligodendrocyte death (**Fig. 2f**). The average power modulation curves varied slightly for each mouse due to differences in the thickness and opacity of the meninges at the surface of the brain under chronic cranial windows **(Extended Data Fig. 5**). The average power modulation by depth curves that were applied over multiple months of 3P imaging without damage can be described by the exponential equation:

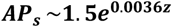

Where is ***AP***_*s*_ is Average Power at the Surface (mW) and **z** is Z-depth from the brain surface (µm) **(Extended Data Fig. 5**). The entire depth of the cortex, excluding the white matter, could be imaged with less than 30 mW of average power at the surface.

### Long-term *in vivo* three-photon imaging does not cause tissue damage or cellular reactivity

To determine whether these modifications permitted long-term *in vivo* three-photon imaging without tissue damage, we assessed cellular reactivity, phototoxicity, and molecular stress via immunohistochemistry in tissue from mice that were exposed to three-photon longitudinal imagining for over 2.5 months. Mice were perfused immediately following the final day of *in vivo* imaging and the 385 x 385 x 1000 µm imaged regions were localized, processed for immunofluorescence, and compared to the contralateral cerebral hemisphere as described previously^31,35^ (**Fig. 3, Extended Data Fig. 6**). To assess the sensitivity and specificity of our histological damage readouts, in a separate group of mice, we induced laser lesions by saturating the THG channel at each z-plane during 3D acquisition and perfused the mice 18-24 hours later **(Fig. 3a,b, Extended Data Fig.6**). These laser-induced lesions displayed an increase in the diffuse extracellular THG signal and tissue opacity, suggesting that we induced mild nonlinear- and heating-related damage (**Fig. 2**). First, we found no differences in the cellular reactivity of oligodendrocytes, microglia, or astrocytes between the imaged and contralateral hemispheres in healthy mice that received 2.5 months of optimized long-term 3P imaging **(Extended Data Fig. 6**). In contrast, positive control tissue, showed a marked increase in the density of microglia and astrocytes in the lesion sites (440.7±45.6 vs. 262.2±18.3 microglia / mm^2^; 290.8±76.4 vs. 142.4±49.7 astrocytes / mm^2^, **Extended Data Fig. 6**). Analyses of the RNA oxidation marker 8-hydroxyguanosine (8-OHG)^36^ did not reveal significant oxidative stress responses in neurons to either type of laser exposure (**Extended Data Fig. 6**). Next, we analyzed blood vessel coverage and pericyte density (Lectin-649, vasculature; CD13, pericytes) using automated segmentation^37,38^ and found no effects of long-term three-photon imaging (**Extended Data Fig. 6**). Interestingly, blood vessel coverage increased in acutely damaged positive control mice (22.8±3.4 vs. 16.8±2.3%) suggesting a dilation of the vasculature in response to this type of photodamage. Pericyte density was unchanged following laser injury mice, confirming that this type of photodamage does not cause cellular ablation. Together, these results suggest that while long-term, repeated three-photon excitation has the potential to induce phototoxicity, our engineering and operational controls allow for multi-month longitudinal imaging without the induction of cellular reactivity at the tissue level.

**Figure 3.**
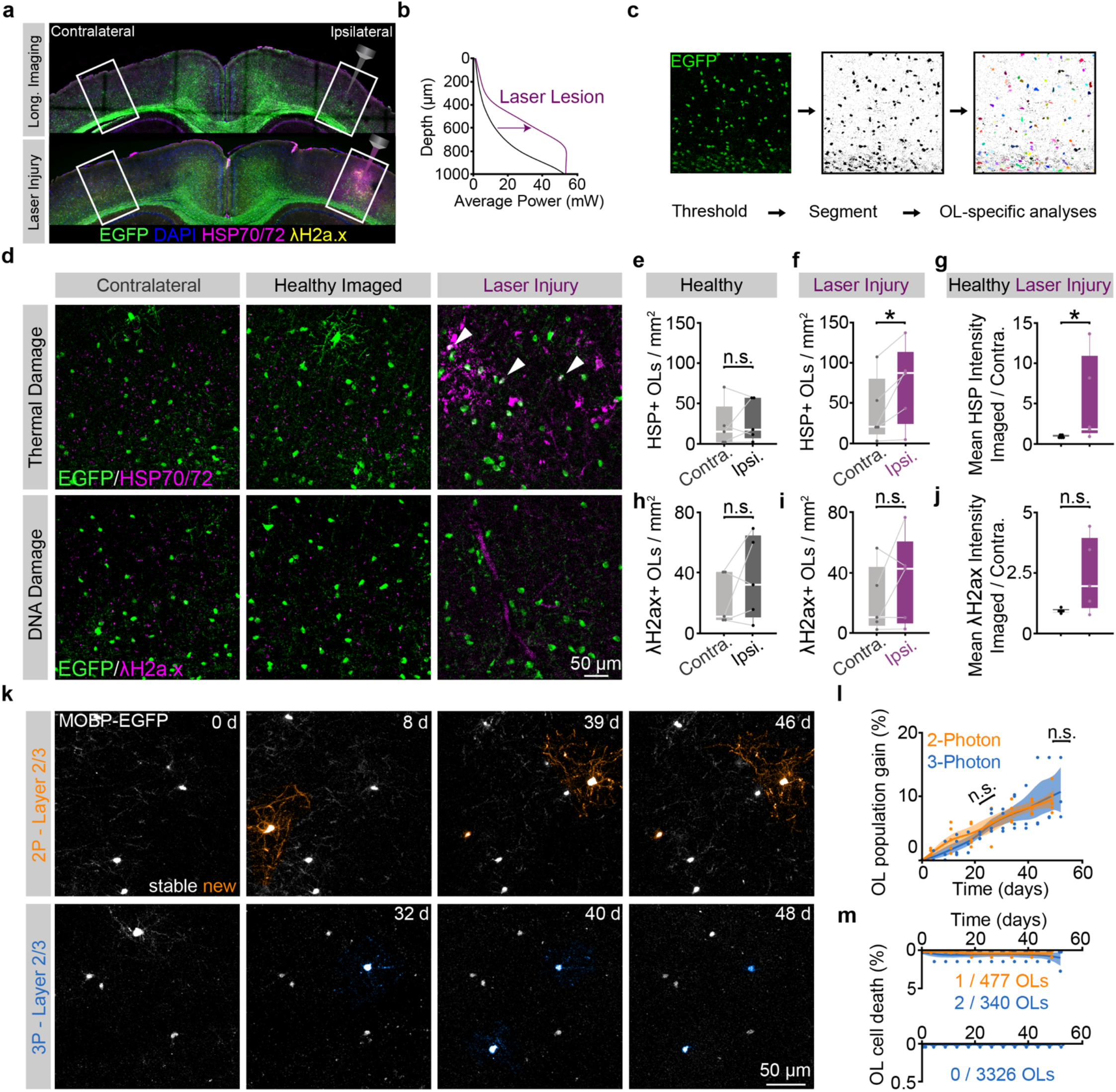
Long-term three-photon imaging does not perturb oligodendrocyte damage markers or healthy oligodendrogenesis. **a**) Coronal brain section from the imaged (right) and contralateral (left) posterior parietal cortex (PPC) of a mouse that was perfused 24 hrs. following 10 weeks of chronic 3P imaging (top) stained for MOBP-EGFP, heat shock proteins 70/72 and the gamma-H2ax marker of DNA double stranded breaks. Coronal brain section from the imaged (right) and contralateral (left) posterior parietal cortex (PPC) of a mouse that received supra-threshold excitation across the cortex and white matter to generate laser-induced positive control tissue (laser injury, right). **b**) To generate laser-damage tissue, the power at the sample was increased to saturate the THG PMT signal without causing tissue ablation or bubbling and the mouse was perfused 24 hrs. following laser exposure. **c**) Example images of oligodendrocyte (OL) cell body segmentation and cell-specific intensity analyses. **d**) High-resolution confocal images of the contralateral (left), longitudinal imaging field (middle) and laser-induced injury (right) regions in the deep cortical layers of the PPC of MOBP-EGFP mice, stained for heat shock proteins 70/72 (HSP70/72, thermal damage), and the phosphorylated histone protein H2A.X (gamma-H2A.X, DNA double-strand breaks). **e**) No difference in the density of HSP70/72 – positive segmented OLs (HSP70/72 intensity > 2 * background intensity) between the contralateral and ipsilateral (imaged) hemispheres of healthy mice that underwent multi-month longitudinal 3P imaging with optimized scan and excitation settings. **f**) Increased density of HSP70/72 – positive OLs on the ipsilateral hemisphere of laser-induced injury positive control mice (Paired, two-tailed Student’s t-test, p = 0.041). **g**) Increase in the ratio of the full-field ipsilateral : contralateral hemisphere mean intensity of HSP70/72 in laser-induced injury vs. long-term 3P mice (Wilcoxon rank sum test, z = 2.089, p = 0.037). **h**) No difference in the density of term 3P mice (Wilcoxon rank sum test, z = 2.089, p = 0.037). **h**) No difference in the density –H2a.X – positive OLs (term 3P mice (Wilcoxon rank sum test, z = 2.089, p = 0.037). **h**) No difference in the density – H2a.X intensity > 2 * background intensity) between the contralateral and ipsilateral (imaged) hemispheres of healthy multi-month longitudinal 3P imaging mice. **i**) No difference in the density of term 3P mice (Wilcoxon rank sum test, z = 2.089, p = 0.037). **h**) No difference in the density –H2a.X – positive OLs between the contralateral and ipsilateral hemispheres of laser-induced injury positive control mice. **j**) No difference in the ratio of the full-field ipsilateral : contralateral hemisphere mean intensity of term 3P mice (Wilcoxon rank sum test, z = 2.089, p = 0.037). **h**) No difference in the density –H2a.X in laser-induced injury vs. long-term 3P mice. **k**) Two-photon imaging of stable (white) and newly generated (orange) oligodendrocytes in layer 2/3 of the PPC (top). Three-photon imaging of stable (white) and newly generated (blue) oligodendrocytes in layer 2/3 of the PPC (bottom). **l**) Comparison of cumulative oligodendrocyte gain (%) in the healthy PPC over ∼7 weeks with standard 2P settings (orange) vs. 3P (blue). No significant differences in the mean rate of healthy oligodendrocyte gain per week (Unpaired, two-tailed Student’s t-test for equal variance, t(6) = 0.82, p = 0.440), or the total cumulative OL gain (%) (Unpaired, two-tailed Student’s t-test for equal variance, t(6) = 0.491, p = 0.641) between the two imaging modalities. **m**) Mature oligodendrocytes are stable over time in the adult mouse brain, and 3P imaging does not increase mature cell death. Layers 1-3 (L1-3) represents ∼0-340 µm depth; Layers 4–corpus callosum (L4-CC) represents ∼341-1000 µm depth, as defined by the Allen Brain Atlas. Images in (**c**) are confocal maximum z-projections of 16 µm (2 µm step), pixel size = 0.57 µm. Images in (**m**) are maximum z-projections of 30 µm (3 µm z step), pixel size (2P) = 0.39 µm; pixel size (3P) = 0.75 µm. *p < 0.05, n.s., not significant; cumulative growth curves represent cubic splines with 95% confidence intervals, box plots represent the median, interquartile ranges and minimum/maximum values. For detailed statistics, see **Supplementary Table 3**

### Long-term three-photon imaging does not affect healthy oligodendrogenesis or cell death

Next, we examined if long-term three-photon imaging induced molecular stress or cellular behavioral changes specifically in mature oligodendrocytes. To examine oligodendrocyte molecular stress, we segmented EGFP-positive cell bodies in *MOBP-EGFP* transgenic tissue from long-term three-photon and acute laser injury mice and counted the density of oligodendrocytes that were positive for HSP70/72, a known marker of thermal stress, and γ-H2A.X, a marker induced by DNA damage (**Fig. 3c,d**; **Methods**). We found no differences in HSP70/72- or γ-H2A.X-positive oligodendrocytes between the imaged and contralateral cerebral hemispheres following long-term three-photon imaging (**Fig. 3e,h**). Acute laser injury tissue showed an increase in the density of HSP70/72-positive oligodendrocytes and the fluorescence intensity on the exposed hemisphere (71.7±22.6 vs. 39.2±18.6 HSP+ OLs/mm^2^; 0.91±0.08 (healthy) vs. 5.26±2.46 (laser-injury) **Fig. 3f,g**), confirming that oligodendrocytes respond rapidly to thermal stress, yet the density and intensity of γ-H2A.X-positive oligodendrocytes did not reach statistical significance (**Fig. 3i-j**). To determine whether three-photon longitudinal *in vivo* imaging affects the dynamics of oligodendrogenesis in the healthy brain, we implanted chronic cranial windows over the posterior parietal cortex in two groups of age-matched mice and compared the rate of oligodendrogenesis following longitudinal two-photon or three-photon imaging in the superficial cortex (0 – 350 µm depth) (**Fig. 3k**). We found no difference in the rate or total oligodendrocyte population gain (%) between groups imaged with different modalities (two-photon = 1.7±0.2 vs. three-photon = 1.4±0.3% gained per week, **Fig. 3l**). Finally, we explored whether longitudinal three-photon imaging affected oligodendrocyte survival. Mature oligodendrocytes in the healthy brain are remarkably stable and seldomly undergo cell death even when exposed to longitudinal two-photon imaging conditions^19,25,39^. In the superficial cortex, we found 1/477 mature oligodendrocytes were lost in the two-photon imaging group compared to 2/340 oligodendrocytes during three-photon imaging (**Fig. 3m**). In regions below those accessible with longitudinal two-photon imaging (350 – 1000 µm depth), 0/3326 tracked oligodendrocytes were lost during 9.5 weeks of longitudinal three-photon imaging in healthy mice (**Fig. 3m**; n = 4-5 mice per imaging modality). These results show that quantitative measures of health and behavior of oligodendrocytes obtained from long-term *in vivo* three-photon imaging in age- and region-matched mice are indistinguishable from those acquired via longitudinal two-photon imaging.

### Increased population expansion of gray matter oligodendrocytes

We used long-term three-photon *in vivo* imaging to track oligodendrogenesis across the entire depth of the posterior parietal cortex (gray matter, GM) and the subcortical corpus callosum (white matter, WM) over multiple months in healthy mice (**Fig. 4a-d**). Similar to past histological studies^3,4,40^, we found that the white matter generated approximately 3X the number of new oligodendrocytes per volume compared to the cortical gray matter (33.8±5.5 vs. 10.9±1.3 OLs / imaging volume; 3.9±0.5 vs. 1.2±0.2 OLs / volume / week, **Fig. 4e**). This difference in the density of newly-generated oligodendrocytes may be related to regional differences in oligodendrocyte precursor cell (OPC) density and proliferation^41,42^. To examine OPC proliferation in these regions we delivered EdU, a thymidine analog carrying an ethynyl group, (5 mg/kg, 2x per day i.p.) for one week (5 mg/kg, 2x per day i.p.) to P70 mice (identical to onset of three-photon *in vivo* imaging). Tissue collection and post-hoc immunohistochemistry for Platelet-Derived Growth Factor Receptor Alpha (PDGFR α), a ubiquitous marker of oligodendrocyte precursors, and EdU were performed to assess OPC density and proliferation (**Extended Data Fig. 7**). We found that the density of OPCs is ∼1.4X higher in the white compared to the gray matter of the PPC (248.6±23.8 vs. 181.2±7.7 OPCs/mm^2^). Furthermore, we found that a ∼3.5X higher percentage of OPCs were EDU-positive in the white matter compared to the gray matter (51.6±6.6 vs. 14.5±2.5%). Together, these results suggest that region-specific regulation of OPC proliferation and density may support the increased number of adult-generated oligodendrocytes in the white matter.

**Figure 4.**
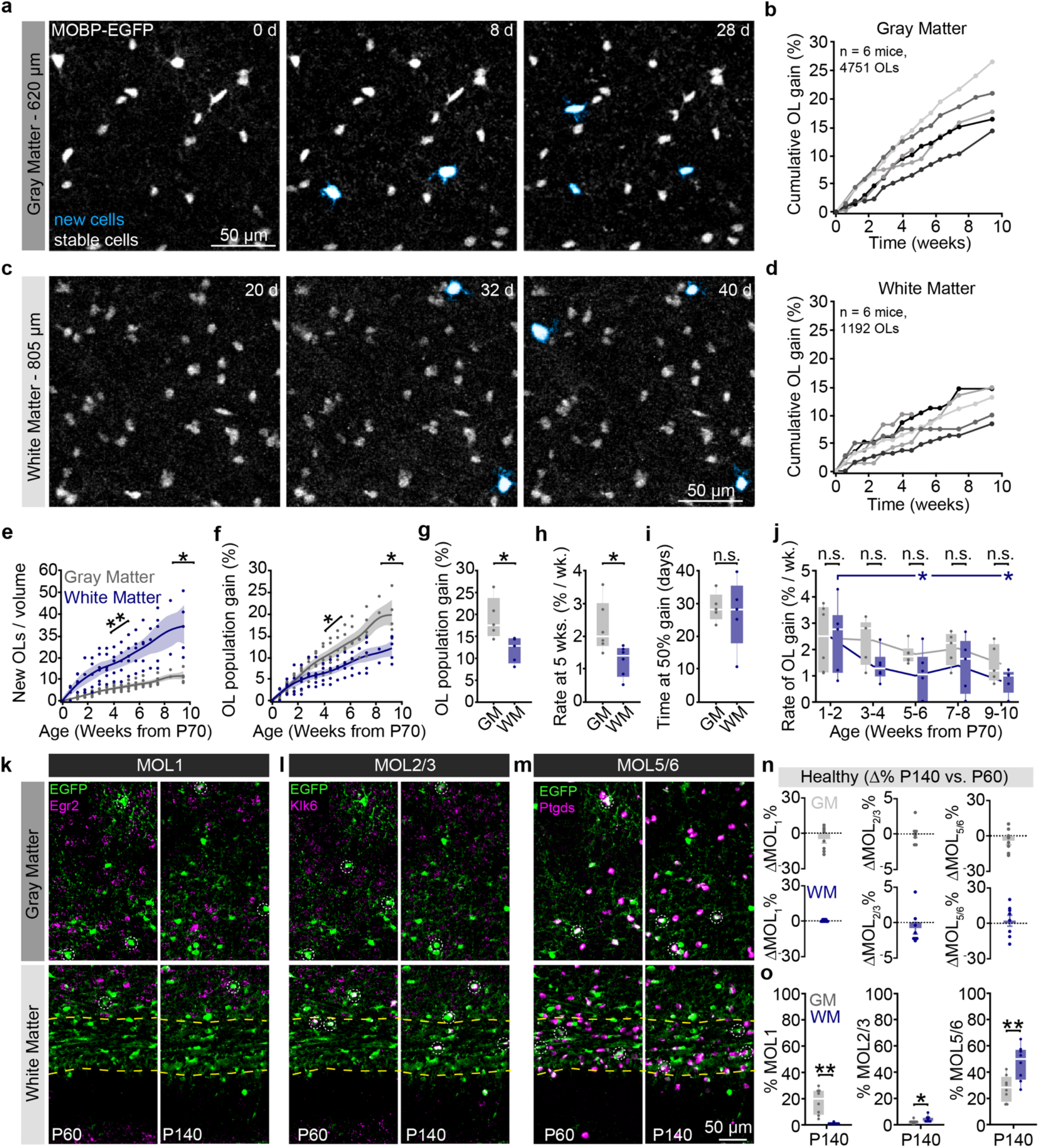
Gray versus white matter differences in population growth correlate with transcriptional heterogeneity. **a**) Oligodendrogenesis (cyan) over 4 weeks of imaging in layer 5 of the posterior parietal cortex (PPC). **b**) Cumulative OL gain curves for individual mice over 10 weeks in the entire depth of the gray matter (GM). **c**) Oligodendrogenesis in the dorsal corpus callosum over ∼6 weeks of imaging in the white matter (WM). **d**) Cumulative OL gain curves for individual mice in the WM. **e**) The WM gained substantially more OLs per imaging volume at a faster rate than the GM (Unpaired two-tailed Student’s t-test for unequal variance, t(4.45) = 4.03, p = 0.013 for total gain at 66 d; Unpaired two-tailed Student’s t-test for unequal variance t(6.86) = 5.14, p = 0.002 for mean rate). **f**) The cumulative percentage gain relative to the initial population of OLs and the rate of OL population gain (%) were higher in the GM vs. WM. **g**) Total population gain was increased in the GM vs. the WM (Unpaired two-tailed Student’s t-test for equal variance, t(8) = -2.83, p = 0.022). **h**) The rate of OL population growth during the fifth week of imaging (∼P100) was increased in the GM vs. WM (Unpaired two-tailed Student’s t-test for equal variance, t(10) = -2.756, p = 0.020). **i**) The time at 50% of the maximum cell gain of the mechanistic growth curves did not differ between GM and WM. **j**) The rate of white matter population gain (% per week) decreased more rapidly with age than in the gray matter (Dunnett’s comparison with control, p = 0.037 for Weeks 5-6 vs. Weeks 1-2; p = 0.022 for Weeks 9-10 vs. Weeks 1-2). **k-m**) Spinning disk confocal images of the distribution of MOL1+ (EGFP/Egr1+), MOL2/3+ (EGFP/Klk6+), and MOL5/6+ (EGFP/Ptgds+) oligodendrocyte populations in the PPC and subcortical WM at P60 (left) and P140 (right). **n**) No differences in the percent change of population proportions for MOL1, MOL2/3, and MOL5/6 between P60 and P140. **o**) Differences in transcriptional heterogeneity between the GM and WM. The proportion of MOL1+ OLs was higher in the GM vs. WM (Unpaired two-tailed Student’s t-test for unequal variance, t(7.033) = -4.804, p = 0.002). The proportion of MOL2/3+ OLs was lower in the GM vs. WM (Wilcoxon rank sum test, z = 2.363, p = 0.018). The proportion of MOL5/6+ OLS was lower in the GM vs. WM (Unpaired two-tailed Student’s t-test for equal variance, t(14) = 3.283, p = 0.005). Data in **h,i** were calculated from the slope and inflection point of the mechanistic growth curves, respectively. Data in **j** were calculated as the percentage of the initial cell population generated per week (not modeled). Data in **a-j** represent n = 6 mice / group. Data in **n-o** represent n = 8 mice, two sections per mouse. *p < 0.05, **p < 0.01, n.s., not significant; cumulative growth curves represent cubic splines with 95% confidence intervals, box plots represent the median, interquartile ranges and the minimum/maximum values. Line plots represent the mean at each time point. For detailed statistics, see **Supplementary Table 3**.

*In vivo* imaging permits a unique perspective to evaluate changes in cellular behavior at the population level. We utilized this long-term approach to determine how the population of mature oligodendrocytes in gray and white matter regions changed over 2.5 months in the adult brain **(Fig. 4f, Methods**). In contrast to regional differences in the densities of newly-generated oligodendrocytes (**Fig. 4e)**, the oligodendrocyte population in the adult gray matter expands by a greater percentage than the white matter **(Fig. 4f**). The total extent of oligodendrocyte population gain was significantly elevated in the healthy gray vs. white matter ((New OLs / Initial OLs) * 100; 19.1±2.1% vs. 12.1±1.3%, **Fig. 4g**). To examine the temporal dynamics of oligodendrocyte population growth in the adult gray versus white matter, we used exponential mechanistic growth modeling to describe the cell population-based decline in adult growth response protein 1 (Egr1), Kallikrein-6 (Klk6 and Prostaglandin D1 Synthase (Ptgds). These markers represent the distinct subpopulations of mature oligodendrocytes, MOL1, MOL2/3, and MOL5/6, respectively, that differ with development, brain region, and in responses to injury^6–8^. MOL1-positive oligodendrocytes were predominantly detected in the gray matter, MOL2/3-oligodendrocytes in the deep cortex and white matter, and MOL5/6 oligodendrocytes in both the gray and white matter (**Fig. 4k-m**). We found no differences in the relative expression of these oligodendrocyte subtypes between P140 and P60 indicating that heterogeneity in these regions is largely established by P60 (**Fig. 4n**). The representation of oligodendrocyte subpopulations exhibited regional heterogeneity at P140, as the MOL1-positive oligodendrocyte population was significantly larger in the gray matter (17.2 ± 3.4 vs. 0.6 ± 0.2%), the MOL2/3-positive population larger in the white matter (1.8 ± 0.5 vs. 3.8 ± 0.8%), and the MOL5/6-positive population proportionally higher in the white matter (27.4 ± 3.4 vs. 46.4 ± 4.7%) (**Fig. 4o**). Analysis of mid-thoracic spinal cord tissue processed in parallel with the brain tissue revealed a similar pattern of spatial heterogeneity to past studies^7^ (**Extended Data Fig. 7**). These data show that there is marked regional oligodendrocyte heterogeneity in the posterior parietal cortex and subcortical white matter well into adulthood (P140).

### Cuprizone-mediated oligodendrocyte population loss does not differ across gray and white matter

Histological studies of regional differences in the cuprizone-mediated demyelination model have produced varied results that differ due to dose, age, time of administration, route of delivery, sex, and/or mouse strain^44–47^. To examine changes to gray and white matter populations of oligodendrocytes before, during, and after oligodendrogenesis with age^43^ (**Extended Data Fig. 8**). Oligodendrogenesis rates were similar between regions at early time points, but significantly decreased in the white matter by 15 weeks of age (2.3±0.3 vs. 1.3±0.2% per week, **Fig. 4h**). The time of 50% population growth did not differ between the gray and white matter (26.8±2.2 vs. 24.3±4.4 days post-P70, respectively, **Fig. 4i**). Binning the raw growth rate data by two-week intervals revealed that the white matter experiences a larger age-dependent decrease in population growth than the gray matter (1.0±0.4 vs. 2.5±0.5, WM, Weeks 5-6 vs. Weeks 1-2; 0.8±0.2 vs. 2.5±0.5, WM, Weeks 9-10 vs. Weeks 1-2, **Fig. 4j**). Together, these data reveal that the gray matter experiences greater oligodendrocyte population expansion that remains elevated later into life than the white matter.

To examine whether regional differences in oligodendrocyte population growth were mirrored by transcriptional heterogeneity in the healthy adult mouse brain, we collected tissue from *MOBP-EGFP* mice at P60 and P140 for RNA *in situ* hybridization (ISH) of Early demyelination, we implanted 10-week-old *MOBP-EGFP* mice with chronic cranial windows over posterior parietal cortex and fed them a diet of 0.2% cuprizone chow for 3 weeks while performing long-term *in vivo* three-photon imaging over approximately ten weeks (**Fig. 5a**). Previous work from our lab and others, showed that longitudinal *in vivo* imaging allows for the visualization of the overlapping time courses of oligodendrocyte loss and regeneration and this cuprizone treatment protocol results in ∼80-90% oligodendrocyte loss in the superficial primary motor or somatosensory cortex^14,19^. To determine the extent of gray and white matter captured with longitudinal *in vivo* three-photon imaging, we used the THG signal to quantify gray and white matter regions of posterior parietal cortex. Our three-photon imaging parameters permitted longitudinal tracking of individual oligodendrocytes throughout the entire gray matter and the dorsal ∼90% of the white matter over the entire imaging time course (**Fig. 5b**). The analyzed proportion of the white matter did not differ between healthy and cuprizone groups (84±8.0% vs. 86.8±3.5%, **Fig. 5c**). Long-term three-photon imaging allowed for the simultaneous visualization of the overlapping time courses of cuprizone demyelination and regeneration in the GM and WM **(Fig. 5d-g**). Volumetric cell loss mirrored differences in the regional density of mature oligodendrocytes (**Fig. 1**), as the white matter lost significantly more oligodendrocytes per volume than the gray matter (157.4±37.8 vs. 36.8±6.0 OLs / imaging volume, **Fig. 5h**). Next, we examined the effects of cuprizone-mediated demyelination on the oligodendrocyte population dynamics in gray and white matter regions. We found that the population loss of oligodendrocytes was equivalent in the gray versus white matter (**Fig. 5i**). These data indicate that the probability of survival of individual oligodendrocytes is similar across the gray and white matter regions, as would be expected from delivery of a systemic oligodendrocyte toxin at a saturating dosage. Accordingly, we found no differences in the total oligodendrocyte population loss (75.3±6.3% vs. 75.6±7.4%), rate of oligodendrocyte population loss (9.0±0.7% vs. 9.2±1.0% loss per week), or the timing of the inflection point of loss between the gray and white matter (0.4±1.4 vs. 2.5±2.0 days post cuprizone, **Fig. 5j-l**). Within-groups analyses of binned rates of demyelination revealed that gray matter oligodendrocyte loss occurs slightly earlier than in the white matter, yet there were no between-groups differences in the rate of oligodendrocyte loss (9.6±1.8 vs. 3.7±0.8 % per week, GM, Weeks ^-^2 to 0 vs. Week ^-^3; 18.9±1.9 vs. 3.7±0.8 % per week, GM, Weeks 0 to 2 vs. Week ^-^3; 22.6±3.1 vs. 4.1±2.5 % per week, WM, Weeks 1 to 2 vs. Week ^-^3, **Fig. 5m**). RNA ISH experiments showed that the prevalence of MOL1, MOL2/3, and MOL5/6-positive oligodendrocytes decreased across both the gray and white matter at four days post-cuprizone cessation (**Fig. 5n-q**, see **Supplementary Table 3** for detailed statistics). While both surviving and newly-generated oligodendrocytes are present four days following cuprizone cessation (**Fig. 5d,f**), we found that the majority of oligodendrocytes at this time point are not labeled with MOL1, MOL2/3, or MOL5/6 subpopulations markers. Despite the reduction in oligodendrocyte subpopulation markers, we found that gray and white matter heterogeneity was preserved for MOL1 and MOL2/3 but abolished for MOL5/6 (1.8±0.7 vs. 0.1%, MOL1; 0.1±0.05 vs. 1.2±0.2%, MOL2/3; 3.1±1.1 vs. 4.9±1.6%, MOL5/6). Together, these data show that cuprizone-mediated demyelination results in similar loss of oligodendrocyte populations and transcriptional heterogeneity across the posterior parietal cortex and underlying corpus callosum.

**Figure 5.**
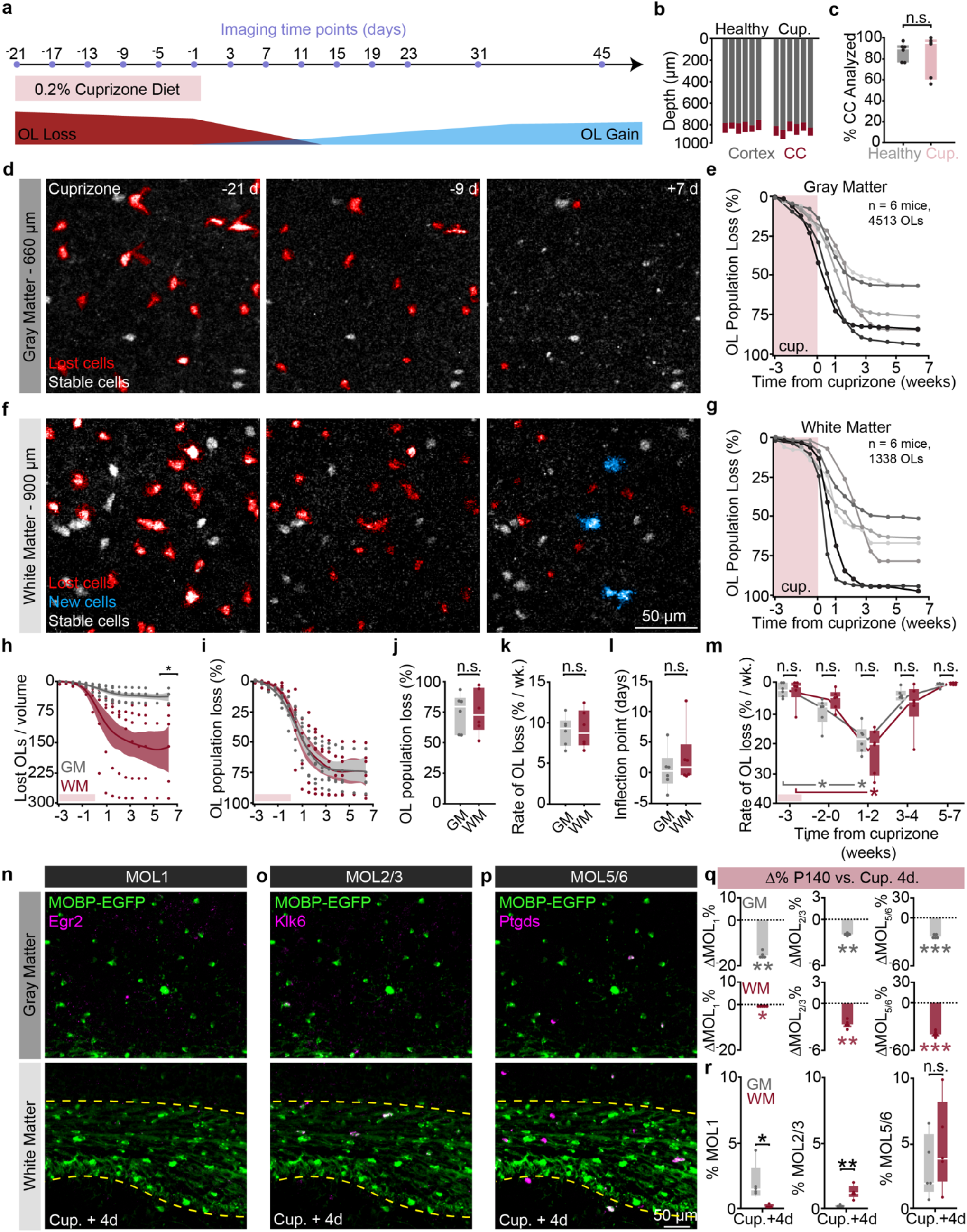
Cuprizone affects oligodendrocyte population decline and molecular heterogeneity similarly across the gray and white matter. **a**) 3P imaging timeline to track oligodendrocyte (OL) loss and gain induced by cuprizone administration. **b**) Depth of PPC and subcortical WM that was imaged over 66 days for n = 6 mice (healthy) and n= 6 mice (cuprizone). **c**) Percentage of the corpus callosum (CC) that was imaged and analyzed longitudinally did not differ by treatment group. **d**) Example time series of cuprizone-mediated OL loss in the deep cortex. **e**) Cumulative OL loss (%) over time in the GM for individual mice. **f**) Example time series of cuprizone-mediated OL loss in the corpus callosum (CC) ventral to the PPC. **g**) Cumulative OL loss (%) over time in the CC for individual mice. **h**) Cumulative number of OLs lost per 350 x 350 x 60 µm imaging volume was lower in the GM vs. WM (Unpaired, two-tailed Student’s t-test for unequal variance, t(5.25) = 3.15, p = 0.024). **i**) Similar dynamics of cumulative % loss of the initial population between regions. **j**) No difference in the total % OL loss for the GM vs. WM (Unpaired, two-tailed Student’s t-test for equal variance t(10) = 0.04, p = 0.970). **k**) No difference in the weekly rate of % OL loss for the GM vs. WM during the loss phase (−21 to +23 days, Unpaired, two-tailed Student’s t-test for equal variance, t(10) = 0.16, p = 0.876). **l**) No difference in the inflection point of the Gompertz 3-parameter cumulative % OLloss curve (Wilcoxon rank sum test, Z = 1.04, p = 0.298). **m**) The rate of GM and WM population loss (% per week), binned by 2-3 week intervals with respect to the 3-week cuprizone administration and late plateau phases. No significant differences were found between the GM and WM at each time point. GM cell loss is increased compared to the first week of cuprizone from -2 to +2 weeks post-cuprizone (Steel method for comparison with control, p = 0.046 for Weeks -2-0 vs. Weeks -3; p = 0.046 for Weeks 1 to 2 vs. Week -3). WM cell loss was increased at 1-2 weeks post-cuprizone compared to the first week of cuprizone (Steel method for comparison with control, p = 0.045 for Weeks 1-2 vs. Weeks 5-7; p = 0.019 for Weeks -2 to 0 vs. Weeks 5-7; p = 0.019 for Weeks 1-2 vs. Week -3). **n-p**) Spinning disk confocal images of the distribution of MOL1+ (EGFP/Egr1+), MOL2/3+ (EGFP/Klk6+), and MOL5/6+ (EGFP/Ptgds+) OL populations in the PPC and subcortical WM at 4 days post-cuprizone removal. **q**) The percent change of population proportions for MOL1, MOL2/3, and MOL5/6 (Healthy P140 vs. 4 days post-cuprizone) were reduced for all markers in the GM and WM (Wilcoxon rank sum test, Z = -2.71, p = 0.007, MOL1 GM; Wilcoxon rank sum test, Z = -2.07, p = 0.038, MOL1 WM; Wilcoxon rank sum test, Z = -2.90, p = 0.004, MOL2/3 GM; Wilcoxon rank sum test, Z = - 2.86, p = 0.004, MOL2/3 WM; Unpaired, two-tailed Student’s t-test for unequal variance, t(8.39) = -6.75, p =0.0001, MOL5/6 GM; Unpaired, two-tailed Student’s t-test for unequal variance, t(8.50) = -8.40, p <0.0001, MOL5/6 WM).. **r**) Differences in transcriptional heterogeneity between the GM and WM 4 days following cessation of cuprizone. The proportion of MOL1+ OLs was higher in the GM vs. WM (Wilcoxon rank sum test, z = -2.538, p = 0.011). The proportion of MOL2/3+ OLs was lower in the GM vs. WM (Wilcoxon rank sum test, z = 2.586, p = 0.0097). No difference in the proportion of MOL5/6+ OLS in the GM vs. WM. Data in **k,l** were calculated from the slope and inflection point of the mechanistic growth curves, respectively. Data in **m** were calculated as the percentage of the initial cell population generated per week (not modeled). Data in **a-m** represent n = 6 mice / group. Data in **q-r** represent n = 5 mice, two sections per mouse. *p < 0.05, **p < 0.01, ***p<0.001, n.s., not significant; cumulative growth curves represent cubic splines with 95% confidence intervals, box plots represent the median, interquartile ranges and the minimum/maximum values. Line plots represent the mean at each time point. For detailed statistics, see **Supplementary Table 3**.

### Enhanced regeneration of oligodendrocyte populations in white compared to gray matter

Since cuprizone treatment affects oligodendrocyte populations in gray and white matter equivalently (see **Fig. 5i,q**), we compared the regenerative capacity of gray and white matter regions following cuprizone-mediated demyelination. We analyzed the region-specific extent, rate, and timing of oligodendrogenesis following cuprizone treatment (**Fig. 6a-d**). Similar to previous fate-mapping studies^47^, we found that there were more newly-generated oligodendrocytes in the white matter compared to the gray matter following cuprizone administration for all mice analyzed (**Fig. 6e**, 123±47.3 vs. 13.3±2.0 OLs/imaging volume). Next, we examined the restoration of the oligodendrocyte populations in these brain regions.

**Figure 6.**
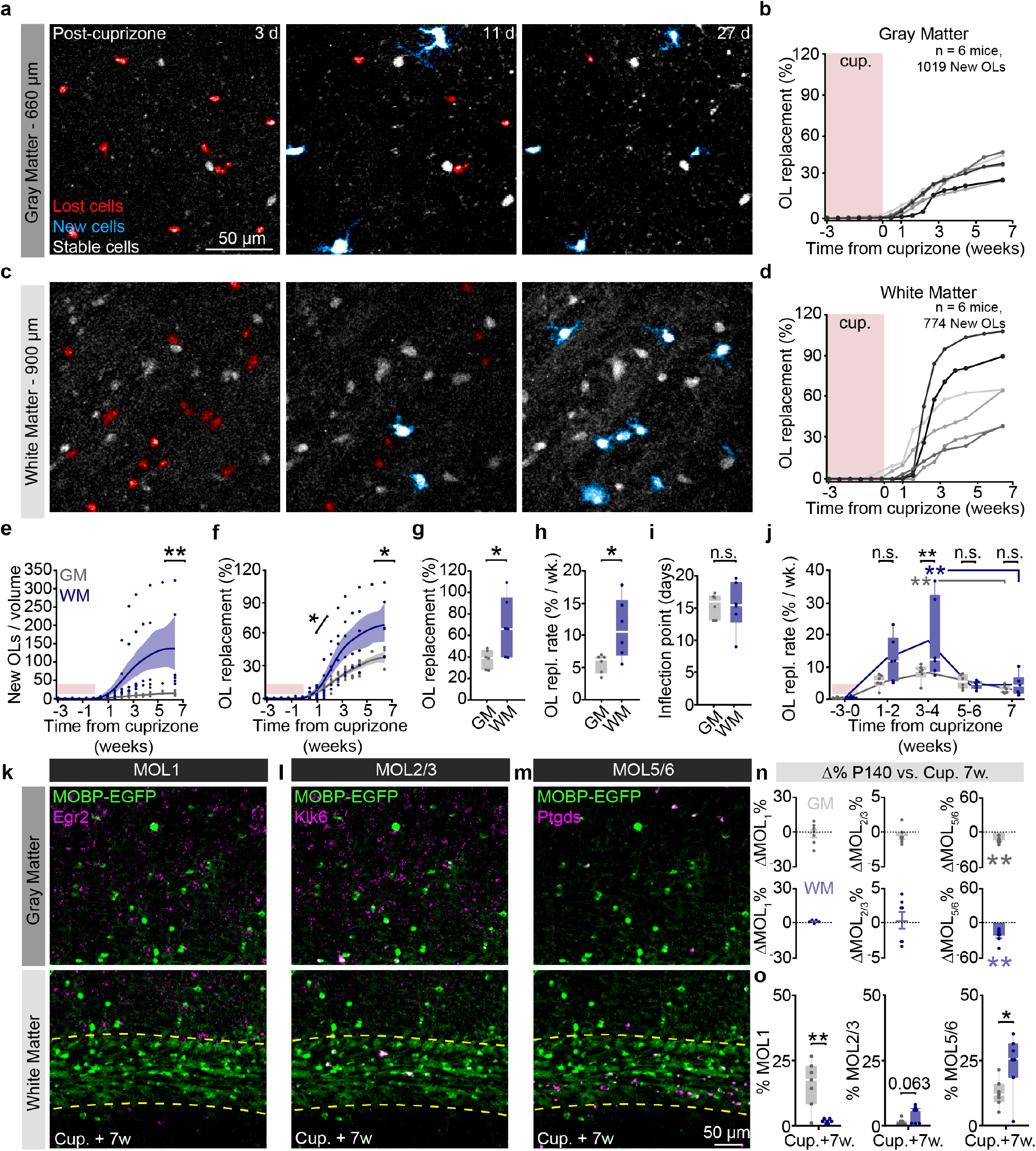
Replacement of lost oligodendrocytes following cuprizone is enhanced in the corpus callosum and regional heterogeneity is partially restored through regeneration. **a**) Timelapse of 24-day period after cuprizone cessation showing oligodendrocyte (OL) cell body compaction and loss (red), followed by the generation of mature myelinating OLs (cyan) in the deep posterior parietal cortex. **b**) Cumulative OL replacement (% gain normalized to % loss) across the cortex for individual mice. **c**) Same time points and mouse as in (**a**) in the corpus callosum. **d**) Cumulative OL replacement (% gain normalized to % loss) for individual mice in the corpus callosum. **e**) Density of newly-generated OLs was significantly increased in the WM compared to the GM (Wilcoxon rank sum test, Z = 2.80, p = 0.005). Note the high variability in the dynamics of OL cell gain calculated by volumetric density. **f**) Cumulative OL replacement over time in WM vs. GM. Significance asterisks represent data in (**g**) and (**h**). **g**) OL replacement (%) is significantly increased at 45 days post-cuprizone cessation in the WM (Unpaired, two-tailed Student’s t-test for unequal variance t(5.97) = 2.57, p = 0.043). **h**) Rate of replacement is enhanced in the WM compared to the GM (Unpaired, two-tailed Student’s t-test for unequal variance t(5.72) = 2.59 p = 0.043). **i**) No difference in the timing of OL replacement between GM and WM. **j**) The rate of GM and WM population replacement (% per week), binned by 1-3 week intervals with respect to the 3-week cuprizone administration and late plateau phases. OL replacement rate (% per week) is significantly increased between the WM vs. GM at the 3-4 week post-cuprizone phase (Two-way ANOVA followed by piecewise Student’s t comparison with Bonferroni correction for multiple comparisons, p = 0.0009). Both the WM and the GM are significantly increased at the 3–4-week phase compared to the 7-week plateau phase (Dunnett’s method for comparison with control, p = 0.002 for Weeks 3-4 vs. Week 7 in the GM; p = 0.0052 for Weeks 3-4 vs. Week 7 in the WM). **k-m**) Spinning disk confocal images of the distribution of MOL1+ (EGFP/Egr1+), MOL2/3+ (EGFP/Klk6+), and MOL5/6+ (EGFP/Ptgds+) OL populations in the PPC and subcortical WM at 7 weeks post-cuprizone removal. **n**) The percent change of population proportions did not differ for MOL1 and MOL2/3 (Healthy P140 vs. 7 weeks post-cuprizone), yet MOL5/6 remained significantly decreased from healthy levels (Unpaired, two-tailed Student’s t-test for equal variance t(13) = -3.712, p = 0.003, GM; Wilcoxon rank sum test, z = -2.720, p = 0.007, WM). **o**) Differences in transcriptional heterogeneity between the GM and WM 7 weeks following cuprizone cessation. The proportion of MOL1+ OLs was higher in the GM vs. WM (Unpaired, two-tailed Student’s t-test for unequal variance t(6.118) = -4.234, p = 0.005). The proportion of MOL2/3+ OLs was unchanged in the GM vs. WM (Unpaired, two-tailed Student’s t-test for unequal variance t(7.560) = 2.187, p = 0.063). The proportion of MOL5/6+ OLs was lower in the GM vs. WM (Unpaired, two-tailed Student’s t-test for equal variance t(12) = 2.432, p = 0.032). Data in **h,i** were calculated from the slope and inflection point of the Gompertz 3-parameter growth curves respectively. Data in **j** were calculated as the percentage of lost cell population that was replaced per week (not modeled). *p < 0.05, **p < 0.01, n.s., not significant; cumulative growth curves represent cubic splines with 95% confidence intervals, box plots represent the median, interquartile ranges and minimum/maximum values. Line plots represent the mean at each time point. For detailed statistics, see **Supplementary Table 3**.

We found that, even when normalizing to the initial population density, the subcortical white matter showed enhanced replacement of lost oligodendrocytes over the 2.5-month imaging timeline compared to gray matter (**Fig. 6f**). Forty-five days following removal of cuprizone, the total oligodendrocyte replacement was 37.6±3.6% vs. 68.0±11.3% for gray versus white matter, respectively (**Fig. 6g**). To analyze the temporal dynamics of post-cuprizone regeneration, we used three-parameter logistic modeling as described previously^14,48^ (**Extended Data Fig. 8)**. Analyzing the Gompertz growth curves of regeneration revealed that the mean rate of oligodendrocyte population replacement per week during remyelination was 5.6±0.6% versus 11.1±1.9% for gray versus white matter **(Fig. 6h**). Yet, the inflection point of oligodendrocyte population growth did not differ suggesting that the time course of the replacement response was similar in both regions (**Fig. 6f,i**). This effect was further supported by analyzing the binned oligodendrocyte growth data in which both the gray and white matter were elevated at the 3-4 week post-cuprizone time bin (8.2±1.1 vs. 3.0±0.5% per week, GM, Weeks 3-4 vs. Week 7; 18.1±5.1 vs. 4.5±1.3% per week, WM, Weeks 3-4 vs. Week 7) yet with a significantly higher peak in the white matter (18.1±5.1 vs. 8.2±1.1% per week, **Fig. 6j**). Finally, since the gray matter replaces 37.6% of lost oligodendrocytes compared to 68% replacement in the white matter, we estimated the time required to reach 100% replacement for each region. Assuming linear growth after 7 weeks post-cuprizone, we estimate that ∼14 weeks would be required to replace the lost oligodendrocytes in the gray matter whereas the white matter would require ∼7 weeks highlighting the reduced recovery rate of gray matter regenerative oligodendrogenesis compared to the white matter. When we examined recovery of oligodendrocyte transcriptional subpopulations via RNA ISH experiments at 7 weeks post-cuprizone cessation (age-matched to final imaging time point) we found that the MOL1 and MOL2 populations returned to healthy proportions in GM, while MOL5/6 remained decreased in both regions (−14.9±2.0%, GM; -22.6±4.2%, WM, **Fig. 6n**). Interestingly, the pattern of healthy gray versus white matter heterogeneity for these markers was nearly restored to healthy conditions (**Fig. 6o**), despite the decreased prevalence of MOL5/6 at this time point (GM vs. WM; 15.8±3.3 vs. 1.5±0.3%, MOL1; 1.2±0.4 vs. 4.1±1.2%, MOL2/3; 12.6±2.0 vs. 24.0±4.2%, MOL5/6). Together, these data show that the white matter restores the lost oligodendrocyte population more effectively than the gray matter, and bulk gray and white matter regions re-establish spatial distributions of oligodendrocytes subtypes to healthy levels following a demyelinating injury.

### Layer-dependent regulation of oligodendrocyte growth, loss, and regeneration

Recent studies using longitudinal *in vivo* two-photon imaging revealed layer-specific differences in cortical oligodendrogenesis following motor learning^14^ and cuprizone-mediated demyelination^19^, yet these studies were limited to the superficial ∼300 – 400 µm of cortex. We utilized the enhanced imaging depth of three-photon microscopy to assess layer-specific differences in oligodendrocyte population dynamics across all cortical layers and the underlying corpus callosum in both the healthy brain and following cuprizone-mediated demyelination. Based on the myelo-, neuronal, axonal, and synaptic anatomy of the adult posterior parietal cortex (**Fig. 7a**), we divided gray and white matter regions into layers 1-3 (superficial cortex), layer 4 (thalamic input layer to the PPC), layer 5-6 (deep cortex), and the subcortical corpus callosum (CC). We assessed layer-dependent differences in oligodendrocyte generation and transcriptional subpopulations across four experimental time points spanning P60 to P140 in healthy and cuprizone-treated mice (**Fig. 7b, Extended Data Fig. 9**). In healthy mice, we found that there was increased oligodendrogenesis in L4 compared to the CC, suggesting that enhanced oligodendrogenesis in this specific cortical microenvironment drives increased population growth in the gray versus the white matter (2.6±0.2 vs. 1.4±0.2% per week, **Fig. 7c-d, Fig. 3**). Analyses of layer-specific oligodendrocyte subpopulations showed the following patterns in the healthy posterior parietal cortex: The proportion of MOL1-positive oligodendrocytes declined with cortical depth and were largely absent from the white matter (**Fig. 7e**). The proportion of MOL2/3-positive oligodendrocytes increased with depth and, within the cortex, were only detected in L5-6 (**Fig. 7f**). The white matter had increased proportion of MOL5/6 oligodendrocytes compared to the gray matter (**Fig. 7g**, see **Supplementary Table 3** for detailed statistics). We found that cuprizone-induced oligodendrocyte loss did not differ across cortical or subcortical layers (**Fig. 7h,i**). The proportion of MOL1-postive oligodendrocytes was increased in layers 1-3 compared to the corpus callosum (**Fig. 7j**), MOL2/3-positive oligodendrocytes were increased in the corpus callosum compared to all cortical layers (**Fig. 7k**), while the proportion of MOL5/6 oligodendrocytes did not differ across cortical and subcortical layers (**Fig. 7l**, see **Supplementary Table 3** for detailed statistics). When examining oligodendrocyte regeneration seven weeks following cuprizone removal, we found that the deep cortex (L5-6) had a decreased rate of oligodendrocyte replacement compared to the corpus callosum (5.6±0.56 vs. 11.2±1.9% per week, **Fig. 7m,n**). At 7 weeks post-cuprizone, the proportion of MOL1-postive oligodendrocytes was increased in layer 4 compared to the corpus callosum (**Fig. 7o**), MOL2/3-positive oligodendrocytes were increased in layers 5/6 and the corpus callosum compared to layers 1-3 and 4 (**Fig. 7p**), while the proportion of MOL5/6 oligodendrocytes was decreased compared to the corpus callosum (10.6±2.2 vs. 24.0±4.2%; **Fig. 7q**, see **Supplementary Table 3** for detailed statistics).

**Figure 7.**
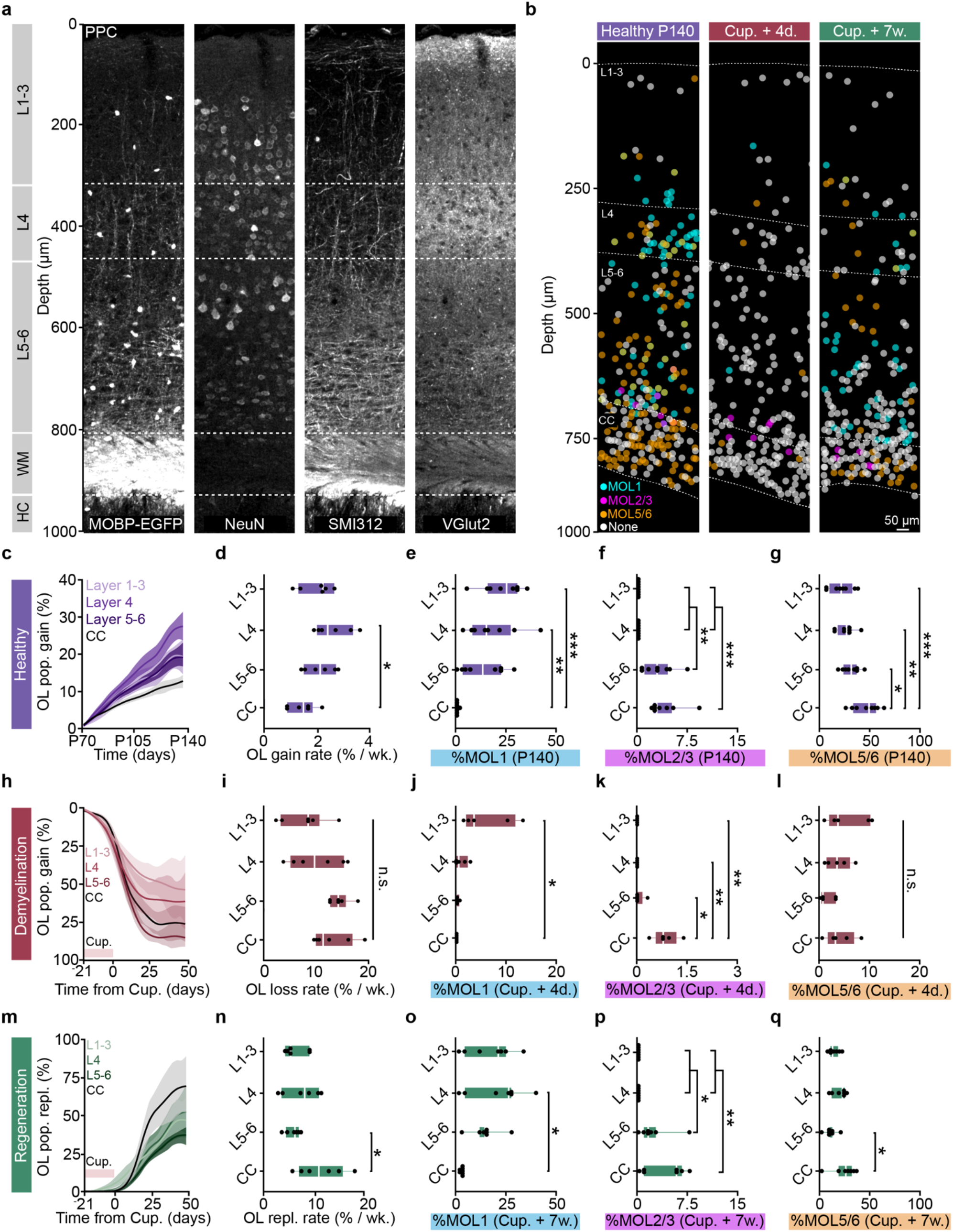
Layer-dependent differences in healthy and regenerative oligodendrogenesis correlate with changes in molecular heterogeneity. a)Confocal image of the cortex and subcortical white matter showing myelo-, neuronal, axonal, and thalamic input architecture of the posterior parietal cortex (PPC). There is a Vglut2-positive layer 4 in the PPC (+2 mm lateral, - 2 mm posterior, from Bregma). **b**) Spatial map of MOL1+ (cyan), MOL2/3+ (magenta), and MOL5/6+ (orange) OLs across the PPC and subcortical white matter at the Healthy P140, Cuprizone + 4 days, and Cuprizone + 7 weeks time points. Approximately ∼50% of the mature OLs were not labeled for any of the three markers (None, gray) and single OLs could have multiple identities in these analyses. **c**) Cumulative healthy OL growth curves plotted by sub-region across the cortical and subcortical depth. **d**) Healthy oligodendrogenesis is increased in layer 4 vs. the corpus callosum (CC, One-way ANOVA followed by Tukey’s HSD, p = 0.012). **e**) The percentage of MOBP-EGFP OLs that were positive for Egr1 (%MOL1) decreased with depth in the healthy brain and was significantly increased in L1-3 and L4 compared to the WM (One-way ANOVA followed by Tukey’s HSD, p = 0.0004, L1-3 vs. CC; p = 0.006, L4 vs. CC). **f**) The percentage of Klk6-positive OLs (%MOL2/3) increased with depth in the healthy brain and was significantly increased in L5-6 and CC vs. L1-3 and L4 (Kruskal-Wallis test followed by Dunn’s test for multiple comparisons, p = 0.0034 (L4 vs. L5-6); p = 0.0034 (L1-3 vs. L5-6); p = 0.0009 (L4 vs. CC); p = 0.0009 (L5-6 vs. CC)). **g**) The percentage of Ptgds-positive OLs (%MOL5/6) was greater than 22% in all analyzed regions, but also increased with depth in the healthy brain (One-way ANOVA followed by Tukey’s HSD, p = 0.042 (L5-6 vs. CC); p = 0.005 (L4 vs. CC); p = 0.0005 (L1-3 vs. CC)). **h**) Cumulative OL loss curves plotted by subregion across the cortical and subcortical depth. **i**) No layer-dependent differences in cuprizone OL loss rate per week between L1-3, L4, L5-6, and the CC. Note the increased variability of demyelination in the superficial layers L1-4. **j**) Layer-specific differences in %MOL1-positive OLs across cortical and subcortical layers at 4 days post-cuprizone removal (Kruskal-Wallis test followed by Dunn’s test for multiple comparisons, p = 0.016, L1-3 vs. CC). **k**) Layer-specific differences in %MOL2/3-positive OLs across cortical and subcortical layers at 4 days post-cuprizone (Kruskal-Wallis test followed by Dunn’s test for multiple comparisons, p = 0.022 L5-6 vs. CC; p = 0.004 L4 vs. CC; p = 0.004 L1-3 vs. CC). **l**) No differences across layers for the %MOL5/6-positive OLs at 4 days post-cuprizone (One-way ANOVA). **m**) Cumulative OL replacement curves plotted by sub-region across the cortical and subcortical depth. **n**) OL replacement (% of lost cell population) is decreased specifically in L5-6 vs. the CC (One-way ANOVA followed by Tukey’s HSD, p = 0.035). **o**) The percentage of MOL1-positive OLs was specifically increased in L4 compared to the WM at 7 weeks post-cuprizone cessation (One-way ANOVA followed by Tukey’s HSD, p = 0.012). **p**) The percentage of MOL2/3-positive OLs was increased in L5-6 and the CC compared to the superficial layers 1-4 (Kruskal-Wallis test followed by Dunn’s test for multiple comparisons, p = 0.025 (L1-3 vs. L5-6); p = 0.025 (L4 vs. L5-6); p = 0.003 (L1-3 vs. CC); p = 0.003 (L4 vs. CC)). a)The percentage of MOL5/6-positive OLs was only significantly decreased in L5-6 vs. CC (One-way ANOVA followed by Tukey’s HSD, p = 0.015)). Data in **c-d, h-I, m-n** were calculated with the raw % growth or % replacement, n=6 mice per treatment group. **E-g, j-l, o-q**, n= 5-8 mice per group. *p < 0.05, **p < 0.01, ***p < 0.001, n.s., not significant. Cumulative growth curves represent cubic splines with 95% confidence intervals, box plots represent the median, interquartile ranges and minimum/maximum values. For detailed statistics, see **Supplementary Table 3**.

### Insufficient oligodendrogenesis and regeneration of MOL5/6 subpopulation in deep cortical layers

To determine the layer-specific capacity to generate heterogeneous oligodendrocyte subpopulations across health and disease, we examined the dynamics of oligodendrogenesis and transcriptional oligodendrocyte subpopulations in healthy and cuprizone treated mice. Compared to the healthy brain, we found that regenerative oligodendrogenesis following cuprizone treatment was increased by an order of magnitude (1 – 7% vs. 0.1 – 0.65%, **Fig. 7d,n**). Therefore, we normalized the modeled growth curves to the maximum value of oligodendrocyte gain or replacement to enable between-groups comparisons of healthy vs. regenerative oligodendrogenesis (**Fig. 8a**; **Methods**). We found that the scaled total oligodendrocyte population growth (0.3±0.03 vs. 0.5±0.1, **Fig. 8a,b**) and rate of growth (0.3±0.03 vs. 0.6±0.1, **Fig. 7a,c**) in L5–6 was decreased compared to healthy conditions, yet the growth curves had equivalent relative inflection points (**Fig. 8d**). Further analyses of the dynamics of regeneration revealed layer-dependent differences in response duration (full-width at half-maximum) and a decrease in the integrated response (area under the curve, above healthy) in L5-6 compared to the corpus callosum (19.89±2.14 vs. 11.70±2.37 days; 22.8±2.9 vs. 56.2±12.1%, **Extended Data Figure 10**).

**Figure 8.**
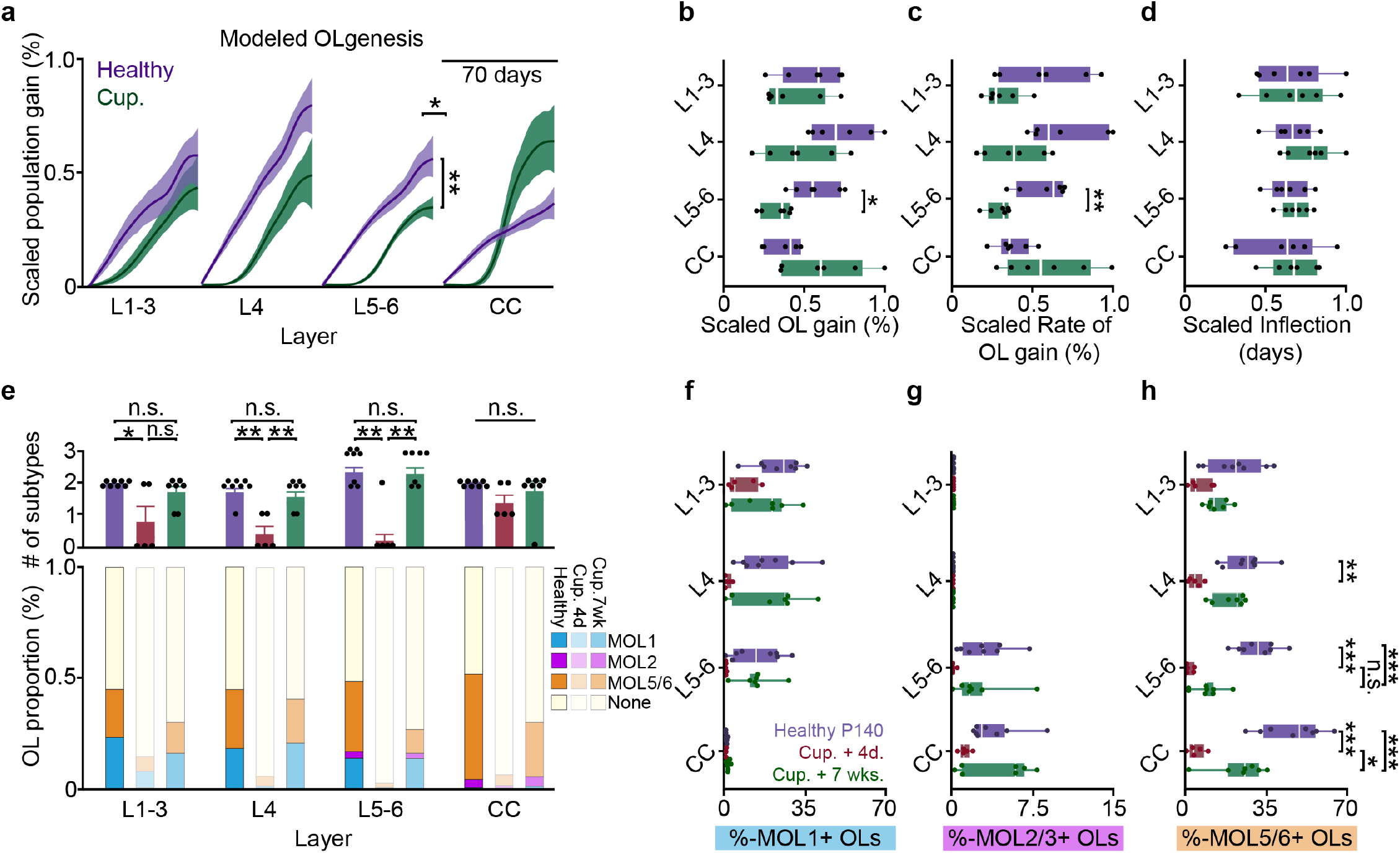
Decreased oligodendrogenesis and insufficient recovery of molecular identity in deep layers 5-6 drive diminished cortical regeneration. **a**) Growth curves plotted by cortical sub-region and scaled to the maximum % gain or replacement to enable between groups comparisons of healthy vs. regenerative oligodendrogenesis. Significance asterisks are related to data plotted in (**b-c**). **b**) Significant decrease in scaled total OL replacement (%, modeled maximum at 66d) in layers 5-6 after demyelination relative to healthy OL gain (%, modeled maximum at 66d,Two-way ANOVA followed by pairwise comparisons with Bonferroni correction within cortical layers, p = 0.006). **c**) Significant decrease in scaled rate of OL replacement in L5-6 compared to healthy OL gain. (Two-way ANOVA followed by pairwise comparisons with Bonferroni correction, p = 0.0025) **d**) No differences in the scaled inflection points across regions. **e**) (Top) Differences in the number of molecular subtypes (MOL1, MOL2/3, MOL5/6) represented at the Healthy P140, Cup. + 4d. and Cup. + 7-weeks time points (classes were counted if the %-positive cells were greater than the minimum detected proportion in healthy mice; Kruskal-Wallis test followed by Dunn’s test for joint ranks). (Bottom) Stacked bar chart showing changes in the relative proportion of MOL1-, MOL2/3- and MOL5/6-positive OLs at 7 weeks post-cuprizone treatment. **f**) No differences in the percentage of MOL1-positive OLs within individual cortical layers, compared between the three represented time points (Two-way ANOVA). **g**) No differences in the percentage of MOL2/3-positive OLs within individual cortical layers, compared between the three represented time points (Two-way ANOVA). **h**) The percentage of MOL5/6-positive OLs decreased significantly at four days post-cuprizone cessation in L4, L5-6, and the CC (Healthy P140, purple vs. Cuprizone + 4 days, red; Two-Way ANOVA followed by Tukey’s HSD; p = 0.001, L4; p = 0.0001, L5-6; p = 0.0001, CC). The percentage of MOL5/6-positive OLs returned significantly at 7 weeks post-cuprizone only in the CC (Cup. 4d (red) vs. Cup. 7w (green), Two-way ANOVA with Tukey’s HSD, p = 0.012). The percentage of MOL5/6-positive OLs returned to Healthy P140 levels in the superficial cortical layers 1-4, but remained decreased in L:5-6 and the CC (Healthy P140 (purple) vs. Cup. + 7wks. (green) Two-way ANOVA followed by Tukey’s HSD, p = 0.0007, L5-6; p = 0.001, CC). Data in **a-d** were derived from the modeled and scaled growth curves. *p < 0.05, **p < 0.01,***p < 0.001, n.s., not significant; Cumulative growth curves represent cubic splines with 95% confidence intervals, box plots represent the median, interquartile ranges and minimum/maximum values. For detailed statistics, see **Supplementary Table 3**.

Next, we analyzed the loss and restoration of layer-specific oligodendrocyte heterogeneity four days and seven weeks following cuprizone removal compared to age-matched healthy controls (**Fig. 8e**). To calculate the overall oligodendrocyte heterogeneity per layer, we determined the number of oligodendrocyte subpopulations that are present within each layer. If the proportion of oligodendrocyte subpopulations was higher than the minimum proportion in healthy control animals, this oligodendrocyte subpopulation was considered to be present within that layer. Four days following cuprizone removal, we found that oligodendrocyte heterogeneity was reduced in all cortical layers whereas oligodendrocyte heterogeneity in the corpus callosum remain unchanged (**Fig. 8e**). Seven weeks following cuprizone removal, we found that oligodendrocyte heterogeneity across all layers was restored to healthy levels (**Fig. 8e**). Following seven weeks of oligodendrocyte regeneration, we found that layer-specific MOL1- and MOL2/3-positive subpopulations were indistinguishable from healthy controls (**Fig. 8f,g**). However, while MOL5/6 oligodendrocytes were regenerated in layers 1-3 and 4, we found this subpopulation remained decreased in layers 5-6 and the white matter compared to healthy conditions (10.6±2.2% vs. 31.3±3.1%, L5-6; 24.0±4.2% vs. 46.4±4.7%, CC; **Fig. 8h**). In the corpus callosum MOL5/6 oligodendrocytes were increased at seven weeks compared to four days following cuprizone removal (24.0±4.2% vs. 4.9±1.6%) indicating ongoing, albeit incomplete, regeneration of this subpopulation. In contrast, the proportion of MOL5/6 oligodendrocytes in layers 5/6 did not differ at seven weeks compared to four days post-cuprizone (10.6±2.2% vs. 1.9±0.8%) indicating a deficit in MOL5/6 regeneration in layers 5/6 following cuprizone-mediate demyelination. Together these results show that deep cortical layers 5-6 have pronounced deficits in regenerative oligodendrogenesis and restoration of the MOL5/6-positive oligodendrocyte subpopulation compared to other cortical and subcortical layers.

## Discussion

The generation of myelin continues throughout life via adult oligodendrogenesis and plays essential roles in cognition and tissue regeneration following injury or disease. Exploring the regulation oligodendrogenesis in specific brain regions is essential to understand how these processes may be harnessed to promote learning or recovery in the adult CNS. In this study, we empirically determined excitation and scanning parameters to allow for longitudinal *in vivo* three-photon imaging over multiple months without cellular reactivity or tissue damage. This approach enabled the analysis of oligodendrocyte generation and population dynamics in the healthy and demyelinated posterior parietal cortex and the subcortical corpus callosum at the level of individual oligodendrocytes. Long-term *in vivo* three-photon imaging confirmed previous findings via fate-mapping and EdU-labeling that the white matter generates substantially more new oligodendrocytes per brain volume compared to gray matter. Conversely, measuring the proportional change of adult gray and white matter oligodendrocyte populations over time revealed an elevated rate of population expansion in the gray matter in five-month-old adult mice. Following demyelinating injury in the absence of autoimmunity, we found that the white matter had enhanced replacement of lost oligodendrocytes irrespective of baseline regional differences in oligodendrocyte density in white and gray matter regions. Additional layer- and subregion-specific analyses revealed a deficiency in oligodendrocyte regeneration in the deep gray matter regions that correlated with a reduced recovery of the MOL5/6 oligodendrocyte subpopulation. As deep cortical layers are essential for transmitting cortical output signals, our results suggest that regional differences in oligodendrocyte regeneration may contribute to deficits in cognitive function observed in human MS patients. Overall, this work provides a roadmap for long-term *in vivo* three-photon imaging of cellular behaviors across the gray and white matter regions of the adult mouse brain.

Despite progress in deep tissue imaging, state-of-the-art two-photon (2P) microscopy systems have limited imaging depths in intact brain tissue and imaging depths decrease over longitudinal imaging timelines. These approaches have not been able to explore deeper areas of the adult rodent brain without tissue removal or insertion of optical lenses which cause widespread tissue damage and glial reactivity. Recent three-photon imaging studies using 1300 nm excitation to examine fluorescent transgenic mice established a practical maximum *in vivo* imaging depth of ∼1.2 mm below the brain surface for acute imaging time points immediately following a craniotomy^22,49^. However, the high peak intensity of excitation pulses required for nonlinear three-photon excitation induce cumulative tissue damage when applied longitudinally over multiple months **(Extended Data Fig. 5**). In this study, we achieved longitudinal imaging depths of ∼1 mm via optimization of optical and scanning parameters in chronic imaging windows from 3 weeks to over 3.2 months following a craniotomy (**Figs. 1,3; Extended Data Fig. 5**). As the effective attenuation length (EAL) of gray matter is nearly 2X that of the white matter^49^, slightly larger imaging depths may be achieved in regions of thicker neocortex, which do not require imaging through the highly scattering subcortical white matter. Our data presents guidelines to develop effective long-term three-photon imaging parameters for a variety of tissues without disrupting sensitive cellular behaviors or inducing cellular reactivity (**Fig. 2**; **Extended Data Fig. 5**). Future work exploring potential laser-induced phototoxicity on transcriptomic or proteomic changes following long-term three-photon imaging may also provide additional insights into the effects of these imaging modalities. Overall, the ability to make longitudinal measurements over weeks-months timescales via *in vivo* three-photon imaging delivers a powerful approach to study cellular behaviors across the gray and white matter regions of the adult mouse brain.

One advantage of longitudinal measurements of cellular behaviors is that the initial population(s) of labeled cells are defined at before, during, and after interventions. This enables the analysis of population growth or decline within a defined subregion with single-cell resolution. We used these methods to examine the link between regional changes in oligodendrogenesis and oligodendrocyte population dynamics. Across development and into adulthood, the rate of oligodendrogenesis in the mouse CNS slows asynchronously across CNS regions and undergoes population decline at late stages of life^4,50^. Although the population density of oligodendrocytes is significantly lower in the gray matter versus the white matter, we found that the population grew at a higher rate in the cortex than in the corpus callosum. These data imply that regional microenvironments regulate the carrying capacity of the oligodendrocyte population to maintain heterogenous myelination dynamics throughout life. Mature oligodendrocytes cluster into at least six specific transcriptional subpopulations in the healthy brain^6^ that have defined spatial preferences in the CNS^7,8^, yet it is unknown how specific regional microenvironments regulate this spatial and transcriptional heterogeneity. Complimentary cross-sectional analyses using RNA *in situ* hybridization revealed gray and white matter regional differences in the proportions of MOL1, MOL2/3, and MOL5/6 oligodendrocytes in the posterior parietal cortex that were stable across young adulthood (P60 to P140). Whether these differences in the transcriptional makeup of oligodendrocytes in these regions reflect specific cellular functions, or simply differences in the maturation state of these populations remains unknown. However, it is thought that the MOL5/6 subpopulation emerges with age in postnatal development and may represent the most-mature oligodendrocyte state^6^, supporting our findings that MOL5/6 is enriched in the white matter and that that the overall population growth declines more rapidly in the white compared to the gray matter. How these region-specific oligodendrocyte population growth dynamics are governed remains an open question. Microenvironmental cues may act at specific stages of oligodendrocyte maturation, such as OPC proliferation, differentiation, or premyelinating cell survival and integration into axonal circuits^25,51^. Similar to previous studies, we found that OPC proliferation is significantly higher in the subcortical white matter compared to the posterior parietal cortex^3,52^(**Extended Data Fig. 7**). As proliferation is linked to OPC differentiation or death^53^, these results suggest a reduced time to cell cycle exit in the white matter that maintains homeostatic population density in response to the increased frequency of oligodendrogenesis in this region. Alternatively, regional variation in integration rates of premyelinating oligodendrocytes^25^ in different microenvironments may regulate regional oligodendrocyte population dynamics. The recent discovery that specific red fluorescent proteins can be excited in the 1300 nm spectral window with high efficiencies^25,54,55^, opens the door to future work with dual-color, oligodendrocyte stage-specific genetic reporter approaches^56^ to dissect how regulation of specific oligodendrocyte maturational stages leads to variable population growth across cortical and subcortical regions.

Similar to lesions in MS patients, the cuprizone-mediated demyelination model results in overlapping oligodendrocyte death and regeneration^57^. The demyelinating injury caused by cuprizone treatment occurs in the absence of notable peripheral immune response^58^ making it an ideal model to evaluate the intrinsic capacity of oligodendrocyte regeneration without the confound of autoimmunity. We found that the survival rate of individual oligodendrocytes is similar across the gray and white matter regions as would be expected from a saturating concentration of an oligodendrocyte toxin. However, a number of factors can affect feeding behavior^59^ and varied oral consumption of cuprizone diet could directly influence the dosage and resulting extent of demyelination in a region-dependent manner at sub-saturating doses. Long-term *in vivo* imaging approaches are essential for analyzing the dynamics of the overlapping periods of oligodendrocyte loss and gain observed with cuprizone-mediated demyelination. Here, we showed that systemic cuprizone administration affects gray and white matter regions similarly, both in terms of oligodendrocyte cell death and changes in transcriptional heterogeneity, yet the white matter has an enhanced intrinsic capacity for regeneration of oligodendrocytes and their subpopulations compared to the gray matter. These gray and white matter differences in intrinsic capacity for regeneration highlight the need for additional insights into the regional effects of “remyelination therapies” (approaches that enhance oligodendrocyte regeneration^60^) to harness their full potential to increase remyelination and accelerate functional recovery following demyelinating injuries. Might remyelination therapies be more successful in augmenting regeneration in the gray matter, where endogenous regeneration is stunted? Further work examining multicellular interactions in regional microenvironments will aid in our mechanistic understanding of differences in regional oligodendrocyte regeneration. For example, the type 1 transmembrane neuropilin-1 is specifically expressed on white matter microglia and transactivates platelet-derived growth factor-α to induce OPC proliferation following demyelination^61^. Conversely, this increased white matter regeneration could be mediated by the recruitment of Gli1-positive adult neural stem cells from the subventricular zone^62^. Protein–protein interaction databases and recent advances in RNA sequencing technologies will likely provide key access to ligand–receptor pairs^63^ that can help explore the role of multicellular interactions in oligodendrocyte population growth and regeneration.

In MS patients, in contrast to our findings with cuprizone-mediated demyelination, lesions in the white matter regions are thought to have more severe demyelination and decreased remyelination compared to gray matter lesions^64,65^. MS white matter lesions differ from gray matter lesions by often displaying blood-brain barrier disruption, infiltrating peripheral immune cells, and complement deposition (reviewed by Geurts and Barkhof^66^) suggesting that the inflammatory microenvironment of white matter limits the regenerative capacity of oligodendrocyte populations. Indeed, mouse white matter oligodendrocyte precursor cells have delayed differentiation when exposed to a combination of interferon-γ and tumor necrosis factor-α compared to gray matter-derived OPCs *in vitro*^67^. Recent findings show that OPCs can cross-present antigen and upregulate the immunoproteasome subunit PSMB8 specifically in the white matter^10,68^ suggesting that oligodendrocyte regeneration is disproportionately affected in the inflammatory context of white compared to gray matter regions in MS patients. Furthermore, regional heterogeneity of oligodendrocytes and other cell types may regulate the disease progression and regenerative capacity in a region-specific manner^11^. Future work using long-term *in vivo* three-photon imaging of oligodendrocytes in gray and white matter regions of immune-mediated demyelination models will provide important insights into the role of inflammation in regional regenerative capacity of oligodendrocyte populations.

Recent longitudinal *in vivo* two-photon imaging following cuprizone-mediated demyelination showed that oligodendrocyte regeneration declined with depth across the superficial 300 µm of somatosensory cortex^19^. Our results confirm and extend these findings as we show that deep cortical layers 5-6 have reduced oligodendrocyte regeneration compared to the baseline level of healthy oligodendrogenesis. Additionally, we identified a specific oligodendrocyte subpopulation in deep layers 5-6 that fails to be reestablished following cuprizone-mediated demyelination. Since mature oligodendrocyte density increases with depth (**Fig. 1d**), one possibility is that increased myelin debris suppresses intrinsic mechanisms of oligodendrocyte regeneration and reestablishment of subpopulation heterogeneity in the deep cortex, but not in the white matter^69^. The consequences of regional deficits in oligodendrocyte regeneration and remyelination are not well understood. Cortical layers 5-6 contain the primary output neurons that project to downstream brain regions to control behavior^70^ and reduced oligodendrocyte regeneration in these regions could underlie the debilitating behavioral deficits experienced by MS patients such as impaired hand function^71^. Oligodendrocytes that closely appose layer 5 pyramidal neurons (satellite oligodendrocytes) locally shape their intrinsic excitability via buffering potassium through the glial syncytium^72^. Loss of these oligodendrocytes could underlie increases in neural firing rates resulting in hyperexcitability following demyelination as well as lead to a loss of local neural circuit synchronization^14,73^. Furthermore, regional variation in oligodendrocyte and myelin loss may lead to cell-type-specific neuron vulnerability and glial activation patterns relevant to neurodegeneration^74^. Remyelination is protective for damaged axons and recent work shows that cell-type specific remyelination is driven by a combination of cell type and axonal diameter^20^. Additional work to understand the specificity of myelin placement on neural circuits across gray and white matter in the healthy and diseased brain will greatly aid in uncovering the potential for remyelination to restore circuit and behavioral function following demyelination.

Emerging optical techniques and approaches will continue to reduce barriers for future work examining the long-term dynamics of subcellular structures such as individual myelin sheaths, axonal boutons, and dendritic spines in deep brain regions. For example, the three-photon effective attenuation length (EAL) is highly dependent on the inhomogeneity factor of the fluorescent signal, therefore adopting a sparse labeling approach will reduce out-of-focus background and increase SBR at depth^75^. In the current study, the *MOBP-EGFP* mice are advantageous for the study of oligodendrogenesis as all of mature oligodendrocytes and myelin sheaths are labeled^25^, however, this transgenic mouse line has less EGFP expression compared to previously lines used for three-photon microscopy (e.g. *Thy1-EGFP*^76^). Moreover, the broad distribution of EGFP signal throughout the cortex generated substantial out-of-focus background, which likely limited the SBR when imaging deep in the brain. Therefore, longitudinal quantifications of cell behavior in the current study were limited to the detection of cell bodies rather than subcellular morphological dynamics or single myelin sheaths. Sparse labeling approaches^25,77^, combined with enhanced detection techniques^22,78^ will allow for the application of longitudinal three-photon microscopy to track subcellular structures at even greater depths in biological tissues^79^. Innovations in adaptive optics like direct wavefront sensing^80^ and zonal aberration correction^33^ will further advance the application of three-photon microscopy to studies of biological dynamics within single cells. Overall, strategies that enhance SBR without increasing average excitation power will be paramount to push the limits of deep-brain three-photon imaging to uncover fundamental discoveries in neurobiology.

## Acknowledgements

We thank Mike Hall for machining expertise, Andrew Scallon and the CU Anschutz Optogenetics and Neural Engineering Core (P30NS048154) for 3D printing and stage design, past and current members of the Hughes and the CU Anschutz Myelin Group for discussions, Drs. Julie Siegenthaler and Stephanie Bonney (Andy Shih Lab) for helpful discussions on the vascular and pericyte analyses. Michael Cammer for assistance with the histogram measurement ImageJ macros (NYU Langone Health Imaging Core). Dominik Stitch and the personnel from the University of Colorado Anschutz Medical Campus Advance Light Microscopy Corefor imaging assistance for the RNA in situ hybridization samples. This imaging was performed in the Advanced Light Microscopy Core Facility of the NeuroTechnology Center at the University of Colorado Anschutz Medical Campus, which is supported in part by Rocky Mountain Neurological Disorders Core Grant (P30 NS048154) and by Diabetes Research Center Grant (P30 DK116073). Funding was provided by the National Institutes of Health NINDS (F31NS120540) to MAT. Funding was provided by the University of Colorado Department of Cell and Developmental Biology Pilot Grant, the Whitehall Foundation, and the National Multiple Sclerosis Society (RG-1701– 26733) and NINDS (NS115975, NS125230, NS132859) to EGH. Funding was provided by the National Institutes of Health NINDS (R01 NS118188 and UF1 NS116241) and the National Science Foundation (BCS-1926676) to EAG and DR. Funding was provided by the National Institutes of Health NEI (R01 EY030841) to APP.

## Author Contributions

EGH and MAT conceived the project. MAT designed and performed experiments, analyzed data, and generated all figures. GLF and EAG built the three-photon microscope. MES, SAB, and MAT performed the RNA in situ hybridization labeling and imaging experiments. MES and SAB analyzed the RNA in situ sequencing data. MES contributed images and data to **Figs. 4-8, Extended Data Figs. 7,9**. SAB contributed images and data to **Figs. 4-7, Extended Data 7,9** and wrote the R code to analyze the QuPath output datasets (see *Code Availability* in Methods section). ANR performed experiments and analyzed data for **Figs. 2,3** and **Extended Data Figs. 4,5**. BNO, OT, and KK developed the software and helped with three-photon microscope setup and expertise. EAG, DR, EGH supervised the three-photon microscope development and application. EAG, DR, APP, and EGH secured funding and EGH supervised the project. MAT and EGH wrote the manuscript with input from other authors.

## Data and materials availability

All data that support the findings, tools and reagents will be shared on an unrestricted basis; requests should be directed to the corresponding author.

## List of Supplementary Materials

Supplementary Tables 1 to 3

Supplementary Figs. 1 to 10

Supplementary Video 1

## Methods

### Animals

All animal experiments were conducted in accordance with protocols approved by the Animal Care and Use Committee at the University of Colorado Anschutz Medical Campus. Approximately equivalent numbers of male and female mice were used in these experiments and were kept on a 14-h light–10-h dark schedule with ad libitum access to food and water. Mice were housed at a temperature of 72 ± 2°C with 50 ± 20% humidity levels. All mice were randomly assigned to conditions and were precisely age-matched (±5 days) across experimental groups. C57BL/6N MOBP–EGFP (MGI:4847238) mice, which have been previously described^81^, were used for *in vivo* multiphoton imaging.

### Custom three-photon microscope

The custom microscope consisted of a VIVO Multiphoton Open (3i) microscope, based on a Sutter Moveable Objective Microscope (MOM) stand, that was modified for multichannel three-photon imaging. The laser output from a regenerative amplifier with 1030 nm center wavelength, 70 W average power, <300 fs pulse duration, and adjustable repetition rate up to 2 MHz (Spirit-1030-70, Spectra Physics, 1 µJ pulse energy at 2 MHz) was wavelength converted to 1300 nm by a noncollinear optical parametric amplifier (NOPA-VIS-IR, Spectra Physics). The laser was operated at a repetition rate of 1 MHz (pulse picking = 2) and the final output power from the idler at this repetition rate was 0.8 to 1.1 W at 1300 nm. The power was modulated using a motorized rotation mount (KPRM1E/M - Ø1, Thorlabs) with a half-wave plate and Glan-Thompson prism. Beam conditioning of the NOPA output consisted of an expansion/collimating lens relay (f1 ¼ 75 mm, f2 ¼ 100 mm, Newport). The beam was expanded using a 4× reflective beam expansion telescope (BE04R, Thorlabs) to uniformly fill the deformable mirror (DM, ALPAO) and then demagnified with a telescope relay back into the microscope (f1 ¼ 500 mm, f2 ¼ 200 mm, Edmund Optics). A dual prism compressor system consisting of two 25 mm SF10 prisms cut at Brewster’s angle (10NSF10, Newport) and a gold roof mirror (HRS1015-M01, Thorlabs) was used to compensate for group delay dispersion. The beam was directed to the galvanometers (Cambridge Technologies) and through a scan lens (SL50-3P, Thorlabs), tube lens, and a 760 nm long-pass primary dichroic (Semrock). Three-photon excitation was focused through a high-NA multiphoton objective (XLPLN25XWMP2, 25x/1.05 NA, Olympus) with the correction collar set for the 0.15 mm coverslip at the surface of the sample. The fluorescent emission was separated from the excitation path by a long pass dichroic mirror, spectrally filtered (green channel 525 / 50 nm, THG channel 430 / 25 nm), and detected by photomultiplier tubes (H10770PA-40, Hamamatsu). The electronic signal was amplified, low-pass filtered and digitized. Laser-clocked acquisition was not implemented on the current system and 2 µs pixel dwell times were used for all experiments to allow for the collection of fluorescence from ∼2 laser pulses per pixel. Data were acquired with SlideBook 2022 (Intelligent Imaging Innovations).

### Cranial window surgery

Chronic cranial windows were prepared as previously described^53^ with minor modifications for three-photon imaging (described below). Six-to eight-week-old mice were anesthetized with isoflurane inhalation (induction, 5%; maintenance, 1.5–2.0%, mixed with 0.5 liter per min O2) and kept at 37 °C body temperature with a thermostat-controlled heating plate. After removal of the skin over the right cerebral hemisphere, the skull was cleaned and a 2 × 2 mm region of skull centered over posterior parietal cortex (1–3 mm posterior to bregma and 1–3 mm lateral) was removed using a high-speed dental drill. A piece of cover glass (VWR, No. 1) was then placed in the craniotomy and sealed with Vetbond (3M) and then dental cement (C&B Metabond). For head stabilization, a custom metal plate with a central hole of diameter = 6 mm was attached to the skull parallel to the implanted glass window, and the dental cement was drilled out around the perimeter of the window to allow objective advancement deep into the brain without headbar contact. A 5 mg / kg dose of carprofen (Zoetis) was subcutaneously administered before awakening and for two additional days for analgesia. *In vivo* imaging sessions began 2–3 weeks post-surgery and took place once every 4 days or weekly (see imaging timeline in Fig. 5). During imaging sessions, mice were anesthetized with isoflurane and immobilized by attaching the head plate to a custom stage (See *Longitudinal three-photon imaging with adaptive optics* section). The internal temperature of the mice was maintained at 37 ± 0.5 C using a feedback temperature controller (TC-1000, CWE) and the Isoflurane was scavenged using a Minivac Gas Evacuation Unit and Charcoal Fluosorber (Harvard Apparatus).

### Longitudinal two-photon imaging

Chronic cranial windows were prepared over the PPC and *in vivo* imaging was performed as above. 4D two-photon datasets were acquired using a Bruker Ultima Investigator upright microscope equipped with Hamamatsu GaAsP detectors and a mode-locked Ti:sapphire laser (Coherent Ultra) tuned to 920 nm. The average power at the sample was between 5 - 30 mW, corresponding to an exponential power curve of 5 - 15 mW to 80 - 120 mW with depth. At least three *MOBP*-EGFP-labeled mature oligodendrocytes distributed across the field of view near the surface of the brain were used to identify the same cortical area over longitudinal imaging sessions. Image stacks were acquired with Zeiss W plan-apochromat ×20/1.0 NA water immersion objective giving a volume of 444 x 444 x 350 µm (X,Y,Z; 1024 x 1024 pixels) from the cortical surface.

### Long-term three-photon imaging with adaptive optics

The custom stage for three-photon imaging consisted of 3D-printed headbar clamps for anesthetized imaging affixed to a small breadboard (MB2020/M, Thorlabs) mounted on 2-axis brass goniometers (GOH-60B50, Optosigma) with the rotation center height approximately level with the cranial window. The first day of imaging (2.5-3 weeks post-surgery) consisted of the following steps: The window was flattened to be orthogonal to the excitation angle using the THG signal generated at the surface of the cranial window^33,82,83^. The tip/tilt of the stage goniometers were manually adjusted while maintaining the focal plane at the surface of the window until the field uniformity of the THG signal was maximized and did not translate significantly when moving in z. Imaging fields were chosen based on image quality at the superficial ∼20 µm of the corpus callosum (∼800 µm depth). Epifluorescent landmark images were taken at the end of the first time point to relocate the imaging field on subsequent time points. To correct for optical aberrations induced by the inhomogeneity of mouse brain tissue, we adopted an indirect, image-based correction technique that has been described previously^22,84^. The deformable mirror (DM, DM97-15, ALPAO) was controlled in SlideBook 2022 and custom MATLAB scripts were used to perform plotting and peak-finding for individual Zernike modes. AO correction was performed on the THG signal of the vasculature and transverse myelin sheaths in the deep cortex just above the corpus callosum (∼750 µm depth) at 256 x 256 resolution, corresponding to 5.1 frames / second. To calibrate for spherical-defocus coupling, we used two-photon imaging of a small ROI surrounding a single 0.2 µm fluorescent bead in a 2-dimensional preparation. The peak intensity of the bead sample was plotted against the spherical and defocus amplitudes at each z-position and the linear relationship between the modal amplitudes was applied to the DM during correction. The raw optimization data was reviewed to confirm that the focal plane was constant at different AO configurations. Parameters for AO correction consisted of the number and order of Zernike modes for correction, DM stroke range, and the stroke increment. We corrected first for the spherical mode followed by astigmatism, coma, trefoil, and then higher-order modes with a DM range of -2 to 2 um and an increment of 0.3, which resulted in the acquisition of 196 images in ∼50 seconds (including analysis time). Zernike mode settings were optimized based on the mean intensity of the THG signal, which resulted in the greatest improvements in EGFP-positive oligodendrocyte signal to background (SBR) ratio in the subcortical WM (Extended Data Fig. 3). A single AO iteration was performed for chronic structural imaging in healthy mice. The single-plane AO correction deep in the cortex resulted in a suboptimal DM pattern for signal quality in the superficial cortex of the z-column. Therefore we implemented a dynamic linear spherical aberration correction that was updated with imaging depth without the need to reoptimize at each z plane. After optimizing the DM settings the power map was set stepwise starting from the meninges. 11 power map points were set (0, 50, 100-1000 µm at 100 µm intervals) using the fluorescent counts on the THG detection channel. The power was slowly increased at 0.2 (surface) or 1% intervals while monitoring the fluorescence intensity histogram to obtain a dynamic range of signal intensity between 0-7000 counts without saturation. In cuprizone mice, the power modulation maps were reduced by ∼10% during the two weeks surrounding peak demyelination (16 – 32 days post-cuprizone) to account for decreased excitation scattering after myelin loss. Large-volume z-stacks were acquired with bidirectional scanning at 512 x 512 pixels (385 x 385 µm), dwell time = 2 µs, frame averaging = 2, and z-step = 3 µm, and scan time ∼.8 frames / s. The PMT gain was set to 91% for THG and 93% for GFP and a custom blackout box was built to reduce ambient light noise. Raster scanned images are only recorded during ∼80% of the x-galvonometer sweep (i.e. during constant velocity), therefore the Spirit pulsed output was blanked during the overscan to reduce the average power applied to the tissue. Finally we implemented a one minute frame pause after every three minutes of z-scanning to allow for periodic heat dissipation. For subsequent imaging time points, mature oligodendrocytes in the superficial cortex were aligned to the first time point using ROI overlays around the cell body positions on day one, the spherical depth correction was applied to the DM as in day one, and the AO correction was performed daily at the same depth before acquisition.

### Cuprizone-mediated demyelination

Global demyelination was induced in our C57\B6N MOBP–EGFP mice using 0.2% cuprizone (Sigma Chemical, C9012), stored in a glass desiccator at 4 °C. Cuprizone was added to powdered chow (Harlan), mixed for ∼10 minutes to ensure homogeneity, and provided to mice in custom feeders (designed to minimize exposure to moisture) for 3 weeks on an ad libitum basis. Feeders were refilled every 2–3 days, and fresh cuprizone chow was prepared weekly. Healthy mice received normal powdered chow in identical feeders. Powdered chow was introduced 2-3 days before cuprizone to ensure acclimation to the powdered food. Cages were changed weekly to avoid build-up of cuprizone chow in bedding, and to minimize reuptake of cuprizone chow following cessation of diet via coprophagia. This 3-week demyelination model that has been previously described^85^, results in ∼88% loss of *MOBP-EGFP* oligodendrocytes in the superficial primary motor cortex, and allows for *in vivo* tracking of the overlapping time courses of demyelination and oligodendrocyte regeneration. Cuprizone mice were weighed weekly during and after cuprizone feeding to confirm consumption of the drug.

### Image processing and analysis

4D longitudinal image stacks were analyzed using Fiji^86^. Cortical layers were defined using the Allen Reference Atlas^87^ and *in vivo* THG imaging data. Layer depth was calculated for individual mice using the depth of the white matter THG signal according to the equation below to eliminate variability in mouse size, field of view location, and imaging modality:

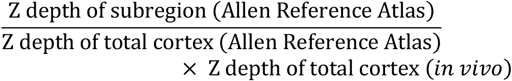

Unprocessed 1000 µm 4D stacks were cut into substacks of layers 1-3 (0 to 356 +/-21 µm), layers 4-6 (469 +/-22 µm) and the subcortical WM (96 +/-6 µm). Image registration was performed iteratively with the Poorman3Dreg plugin for lateral registration followed by the Correct3D drift plugin^88^ (EGFP channel, rigid body registration, 30×30×20 pixels X/Y/Z). Additional iterations of Correct3D drift were performed on high-signal ROIs as needed. The registered datasets were further processed with a median filter to remove salt and pepper noise and individual oligodendrocytes were manually tracked using custom FIJI scripts as previously reported^10^. Images generated for figures were brightness and contrast corrected for clarity and maximum z-projection widths are defined in the figure legends. Stable, new, and lost cells were segmented and pseudo-colored using the Lasso tool in Photoshop CS6 (Adobe).

### Analyses of imaging performance and white matter fiber orientation

For two-photon versus three-photon imaging comparisons, oligodendrocyte cell bodies were counted within x-y subregions greater than or equal to 200 x 200 µm^2^ and oligodendrocytes were only counted if SBR was greater than 2 (as measured by a line scan profile in ImageJ). For the SBR analyses, oligodendrocytes were randomly sampled throughout the imaging volume to include the full range of oligodendrocyte SBRs (i.e. dim, medium, and bright) with at least 10 oligodendrocytes / 100 µm z-depth. ROIs were drawn around the oligodendrocyte cell body and then immediately adjacent to the cell body in a background region. Single oligodendrocyte SBRs were calculated as the max intensity of the cell body / max intensity of the background. AO improvements in oligodendrocyte signal intensity were quantified in the same manner, while improvements in resolution were calculated on oligodendrocyte cell bodies or single myelin sheaths as the full width at half-maximum in the lateral or axial dimensions. Fiber orientation calculations were performed on single z-slices within the WM using the OrientationJ plugin^89,90^.

### EdU labeling and detection

To quantify regional differences in OPC proliferation and oligodendrocyte generation, EdU (A10044, Invitrogen) was diluted in sterile saline solution (0.9% NaCl) and injected intraperitoneally from P70-P76 (matched to starting age of *in vivo* imaging timeline) twice daily at 5 mg per kg for a total of 7 days. The time course and 5 mg per kg doses were chosen based on previously published work^91,92^, with the goal of labeling a substantial number of proliferating OPCs in the gray and white matter without saturating the population within either region^93^ and without generating false positives due to EdU binding at sites of DNA repair^94^ or inducing DNA damage pathways^95^. EdU-labeled cells were visualized using the AlexaFluor-647 Click-iT EdU Cell Proliferation Kit for Imaging (C10340, Invitrogen). The Click-iT reaction was performed per the manufacturer specifications immediately following the blocking step in the immunofluorescence protocol.

### Immunofluorescence analyses

Mice were anesthetized with an intraperitoneal injection of sodium pentobarbital (100 mg per kg body weight) and transcardially perfused with 4% paraformaldehyde in 0.1 M phosphate buffer (pH 7.0–7.4) immediately following the final imaging time point or 16-20 h post-exposure in the laser damage experiments. Brains were postfixed in 4% paraformaldehyde for 1-2 h at 4 °C, transferred to 30% sucrose solution in PBS (pH 7.4) and stored at 4 °C for at least 24 h. Brains were extracted, frozen in TissuePlus OCT, sectioned coronally at 30 µm, and immunostained as free-floating sections. Sections were incubated in blocking solution (5% normal donkey serum, 2% bovine γ-globulin, 0.3% Triton X-100 in PBS, pH 7.4) for 1–2 h at room temperature, then overnight at 4 °C in primary antibody. When staining for Aspartoacylase (ASPA) sections were incubated with Liberate Antibody Binding Solution (L.A.B., Polysciences) for 10 minutes prior to the blocking stop. Secondary antibody incubation was performed at room temperature for 1.5 h. Sections were mounted on slides with Vectashield antifade reagent (Vector Laboratories). Images were acquired using a laser-scanning confocal microscope (Leica SP8). For laser damage experiments, mild columnar damage regions (positive control tissue) were created using identical scanning settings as in the longitudinal group, but with the power map right-shifted to saturate the THG channel. 2-3 coronal sections of the PPC spaced ∼60 µm apart were chosen based on the stereotaxic coordinates of the 385 x 385 µm *in vivo* imaging field and aligned to the reference atlas using anatomical landmarks. Bulk intensity measurements were made as previously described^31,96^ on a 1000 x 1000 µm ROI drawn directly under the 385 x 385 µm imaging field, and in the identical region on the contralateral hemisphere. Irregularly-shaped lesions were cropped with a minimum area of 600 x 600 µm. For cell counts, images with a 0.56 µm pixel size were thresholded using the maximum entropy filter and then cell bodies in individual channels were segmented using FIJI’s particle analysis tool^86^ and then proofread manually using the ROI manager. The settings for the particle analyzer were as follows: Size filter: 40 - 2000, Circularity: 0-.7, Microglia; Size filter: 30 - 2000, Circularity: 0-1, Oligodendrocytes; Size filter: 200 - 3500, Circularity: 0-1 Neurons; Size filter: 30 - 2000, Circularity: 0-0.5; Astrocytes). For the OL-specific analyses of phototoxicity, the mean intensity of each segmented MOBP-EGFP ROI was measured together with a background ROI in the superficial cortex. Cells were considered HSP70/72- or γ –H2a.X-positive if the mean intensity within the cell body was greater than two times the background measurement. Automated segmentation of the vasculature was performed as previously described^37,38^. Briefly, one random section from each mouse was concatenated into a z-stack and ∼15 blood vessels of varying diameters and background regions were traced for each image using the pixel classifier. The classifier was saved and batch-applied to all the images in the dataset (available upon request), which were then binarized and analyzed for percentage area of vascular coverage. Pericytes were counted as above. All confocal images shown in Fig. 3 and Extended Data Fig. 6 were acquired with identical power settings, processed with a 100 pixel rolling ball background subtraction and 0.7 pixel gaussian blur, and brightness/contrast correction was applied identically across channels.

### RNA in situ hybridization with immunofluorescent detection of EGFP In situ

hybridization (ISH) was performed using the commercially available RNAscope V2 kit by adapting the manufacturer’s protocol (ACD Biotechne 323100). This procedure was specifically modified to include antibody labeling of EGFP+ oligodendrocytes in the same tissue sections labeled with three mRNA probes. Probes used in this study were designed for mouse Ptgds-C1 (ACD Biotechne 492781), Klk6-C2 (ACD Biotechne 493751), and Egr2-C3 (ACD Biotechne 407871). Briefly, 20 um, PFA-fixed, tissue sections from brain and spinal cord were washed and mounted on Superfrost microscope slides. Tissue dehydration, antigen retrieval, protease treatment, and probe hybridization were carried out according to the manufacturer’s protocol. Amplification and HRP reaction steps for C2, C3, and C1 (in this order) were carried out without deviation from the manufacturer’s protocol. Briefly, slides were first washed with 1x RNA Scope wash buffer 2x for 2 minutes at room temperature before incubation with Amp 1 for 30 minutes at 40C. This process was then repeated for Amp 2 and Amp 3. Following amplification steps slides were washed 2x for 2 minutes each in RNAscope wash buffer before incubation with the respective HRP conjugate for 15 min at 40C. Next, sections were washed 2x for 2 minutes each and incubated in the corresponding fluorophore for 30 minutes at 40C (C2, TSA Vivid 570 1:1500 in TSA buffer, C3 TSA Vivid 650 1:2000 in TSA buffer, or C1 Opal TSA-DIG (Akoya) 1:750 in TSA buffer). After HRP reaction slides were washed 2x for 2 minutes, incubated in HRP blocker for 15 min at 40C, and finally washed 2x for 2 minutes. This was then repeated for the remaining 2 probes. Following the final HRP blockade, slides were incubated in blocking buffer (5%NDS, 0.3% Triton X-100 in PBS) for 60 min at room temperature. Slides were then incubated with chicken anti-GFP 1:250 (AVES, GFP-1020) for 48 hours at 4C. Slides were next washed 3x for 15 minutes in PBS. Immunohistochemistry was completed with incubation in Donkey anti-chicken 488 1:500 (Jackson, 703-545-155) for 60 minutes. ISH was completed by washing slides 3x for 15 minutes in PBS before incubation in Polaris 780 (1:125 in diluent buffer, Akoya) for 30 minutes at 40C. Finally, slides were washed 2x for 2 minutes in RNAscope wash buffer and counterstained with RNAscope DAPI for 30 seconds before cover slipping with Prolong Gold (Invitrogen, P36934) for imaging.

### RNAscope image acquisition and processing

Fluorescent images used for RNAscope quantification were acquired using the Olympus DSU Spinning Disc microscope in the Advanced Light Microscopy Core at the University of Colorado Anschutz Medical Campus. To obtain a large field of view, adjacent images were acquired using a 20x air objective and a 10% tile overlap. A z-stack with 4 um steps was used to capture the entire thickness of each tissue section and reduce the detection of false positive labeled cells as described previously^7^. Images were stitched and max projected using a custom ImageJ macro based on the BaSiC plugin^97^.

### RNAscope quantification

Following imaging, images were analyzed using Qupath Software 0.4.398. RNA expression per cell was quantified using a custom analysis pipeline. Briefly, cortical layers were measured and drawn onto each tissue section as described above (*Image Processing and Analysis)* to allow for quantification of mRNA puncta within cells in each individual cortical layer. Nuclei were semi-automatically segmented based on the DAPI channel. Nuclear segmentations were dilated to encompass the predicted area of each cell. Finally, mRNA puncta were segmented and assigned to DAPI+/EGFP+ cell regions for oligodendrocyte-specific quantification. Threshold values for each channel were based on the background signal in each image. Cell and puncta detection was performed on each section in Qupath and the spreadsheets containing the number, size, and area of puncta were exported and processed to calculate the percentage of each population within each cortical layer using custom R scripts (https://github.com/sbudoff/OligoRNAscopeTools). We used thresholds of 14, 12, and 7 estimate puncta per GFP positive cell of *Egr2+, Klk6+, and Ptgds+*, respectively, to assign oligodendrocyte populations. For *Klk6*, we observed that positive cells in the cortical sections expressed high levels of entirely clustered puncta and therefore only included cells with 1 or 2 clusters that had an estimated puncta number greater than 12. To calculate the number of oligodendrocyte classes that are present within each layer we calculated a threshold which was the lowest proportion of cells present in healthy control animals. If an animal had a proportion of oligodendrocytes higher than this threshold it was considered to have this class present within that cortical layer. We used a threshold of 5.73%, 0.394%, and 5.82%, of MOL1, MOL2/3, and MOL5/6, respectively, to determine the presence or absence of each oligodendrocyte class.

### Statistics and Reproducibility

Sample sizes were comparable to relevant publications^8,10,15,80^. All micrographs are representative images showing phenomena seen in multiple mice. All data were initially screened for outliers using IQR methods. All mice in a litter underwent cranial window surgery and concurrent two-photon imaging and training timelines were designated to be a “batch”. When possible, experimental groups were replicated in multiple batches with multiple experimental groups per batch. To analyze *in vivo* changes in oligodendrogenesis, we used cumulative population growth and decline measures as previously described^10^. Briefly, oligodendrocyte gain / loss (%) was calculated as the change in oligodendrocyte number / initial cell number * 100, the rate of oligodendrocyte gain/loss (%) was calculated as the % change in oligodendrocyte number divided by a time interval (e.g. Remyelination phase), and oligodendrocyte replacement (%) as total oligodendrocyte gain / total oligodendrocyte loss * 100. The oligodendrocyte replacement (%) metric is used to control for differences in the magnitude of demyelination, as oligodendrocyte gain is strongly correlated with oligodendrocyte loss in the cuprizone model^1081,82^. To analyze regional differences in the rate and timing of oligodendrogenesis, we used statistical modeling of cell growth and decline. As healthy oligodendrogenesis rates decline with age^99^, we found that this process was best-fit using the exponential mechanistic growth model that is commonly used to describe population growth with competition for resources^100^. Extended Data Fig. 8 shows the mechanistic growth curves for example mice and group means along with R^2^ values, which were greater than or equal to 0.91 (cortex) and 0.96 (CC) for all analyzed mice. The exponential mechanistic growth equation was calculated according to the equation below:

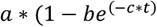

Where *a* = Asymptote, *b* = Scale, *c* = Growth rate, and *t* = time (days). Cuprizone potently inhibits the survival of new oligodendrocytes when it is actively being administered in the diet and cuprizone-induced cumulative OL loss and gain curves are sigmoidal and can be effectively modeled using three-parameter logistic regression^85^. In this study, we used a three-parameter Gompertz model commonly used to describe tumor growth^101,102^ that was robust to asymmetrical differences in the timing of the inflection point. Extended Data Fig. 8 shows the Gompertz growth curves for example mice and group means along with R^2^ values, which were greater than or equal to 0.99 (cortex) and 0.99 (CC) for all analyzed mice. The Gompertz function was calculated according to the equation below:

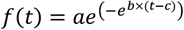

Where *a* = asymptote, *b* = growth rate, *c* = inflection point, and *t* = time (days).

Because the goal of this study was to test the hypothesis that different brain regions are independent with respect to oligodendrogenesis, we used unpaired rather than paired statistics for single comparisons between gray and white matter regions (unpaired two-tailed t-test, parametric; Mann Whitney U test, nonparametric). Statistical comparisons between multiple brain regions were made using one-way ANOVA followed by Tukey’s HSD post-hoc test (parametric) or Kruskal-Wallis followed by Dunn test (nonparametric). To compare oligodendrogenesis in the healthy brain to cuprizone remyelination, we scaled the datasets to the maximum value of healthy or cuprizone oligodendrogenesis. To compare healthy vs. cuprizone-induced oligodendrogenesis metrics we used full-factorial 2-way ANOVA for the interaction effect followed by multiple comparisons with Bonferroni correction. Bonferroni-adjusted significance levels are indicated in the figure legends. Normality was assessed using the Shapiro-Wilk Test and outliers were assessed by visualizing the interquartile ranges. All statistical analyses and modeling were conducted in JMP 16 (SAS).

## Data and materials availability

All data that support the findings, tools, and reagents will be shared on an unrestricted basis; requests should be directed to the corresponding author.

## Code availability

Code for analysis associated with the manuscript is available at https://github.com/EthanHughesLab/; RNA Scope analysis scripts are available at https://github.com/sbudoff/. The code to measure the brightest n% of pixels through a z-stack or time-series was modified from macro scripts available via the NYU Imaging Core website https://microscopynotes.com/imagej/.

## EXTENDED DATA

**Extended Data Fig. 1.**
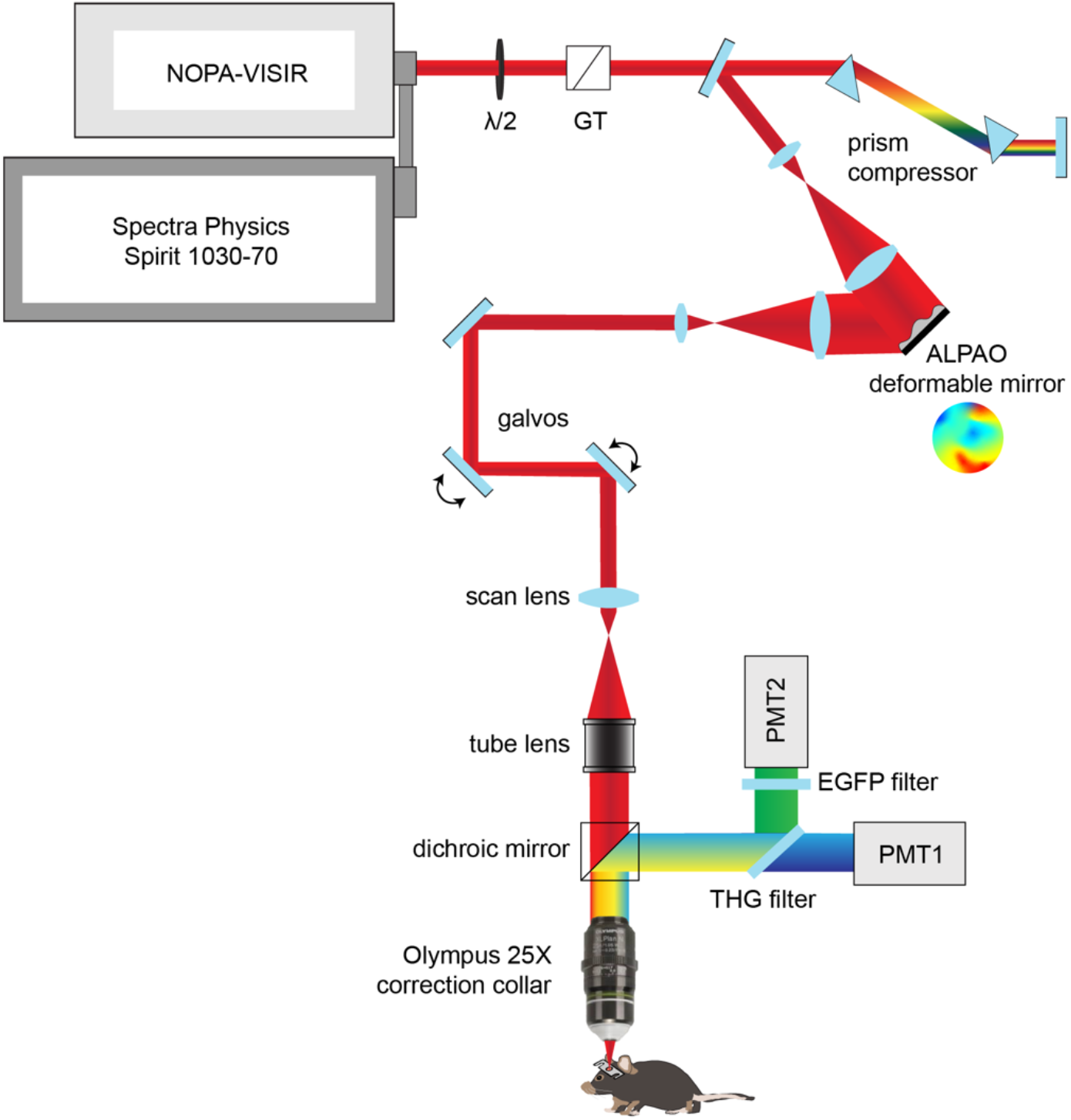
Custom three-photon light path and modifications for longitudinal *in vivo* imaging. 3P excitation source and light path, including motorized half-wave plate for power modulation (λ/2), Glan-Thompson polarizer (GT), dual prism compressor, beam expanding telescope, ALPAO deformable mirror, beam reduction lens relay, galvo-galvo scan mirrors, scan lens, tube lens, dichroic mirror (FF-488-di02, cut at 488 nm), and moveable objective microscope system with collection filters for third harmonic generation signal (430/25 nm, PMT1) and EGFP emission (520/70 nm, PMT2).

**Extended Data Fig. 2.**
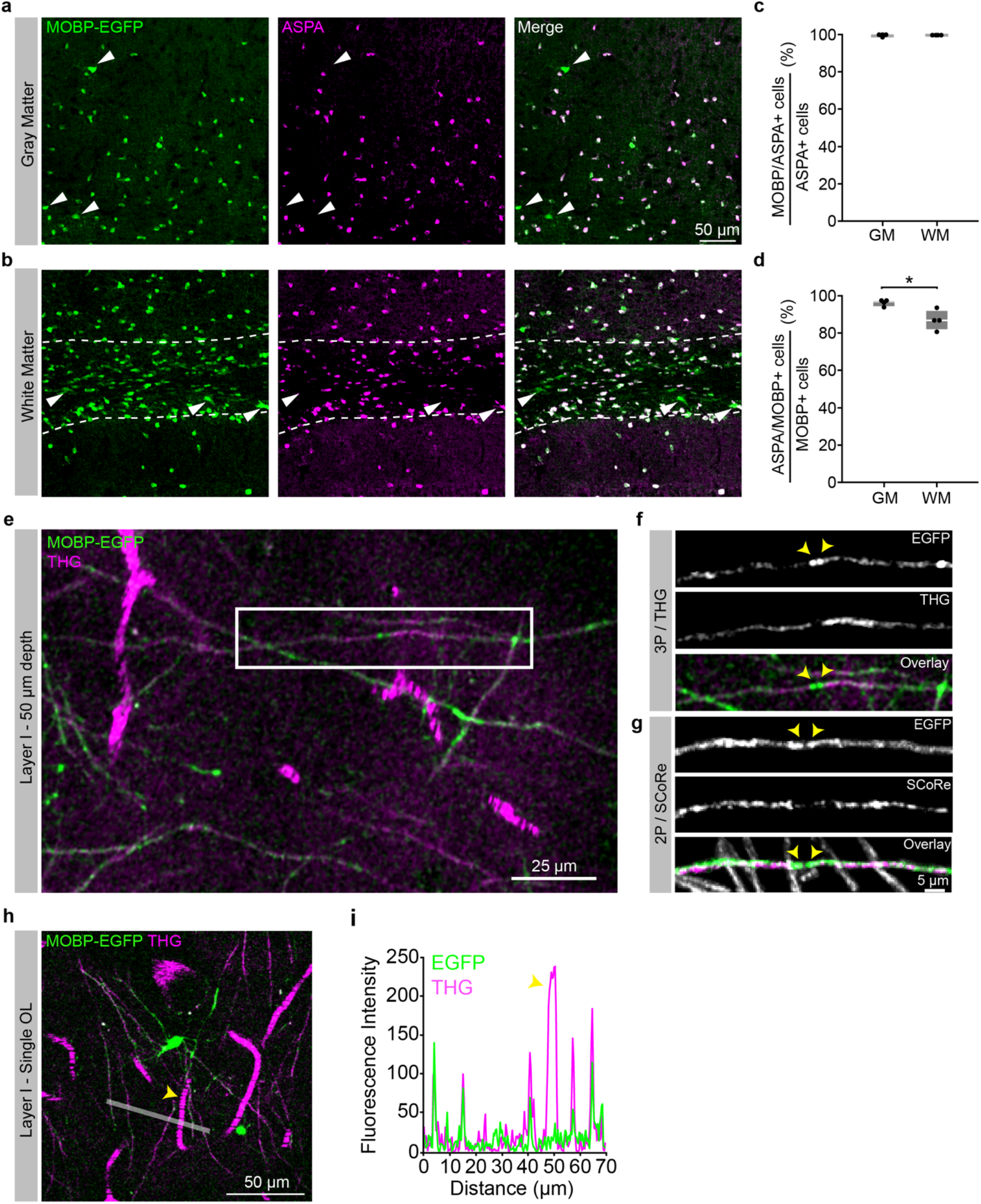
MOBP-EGFP and THG labeling of mature oligodendrocytes and myelin in the cortex and subcortical white matter. **a**) Confocal image of a tissue section from the deep posterior parietal cortex stained for transgenic MOBP-EGFP and ASPA. Arrowheads show large, putative newly-generated oligodendrocytes that are MOBP-EGFP-positive and ASPA-negative. **b**) Confocal image of ASPA/MOBP-EGFP immunofluorescence in the subcortical white matter beneath the posterior parietal cortex (dotted white lines). Arrowheads show MOBP-EGFP-positive and ASPA-negative white matter oligodendrocytes of varying sizes and brightness. **c**) 99.5% (cortex) and 99.6% (white matter) of ASPA-positive cells express MOBP-EGFP. **d**) 95.6% (cortex) vs. 86.5% (white matter) of MOBP-EGFP-positive oligodendrocytes were also positive for ASPA (Unpaired, two-tailed Student’s t-test for equal variance, p = 0.016). n = 4 mice, 2 sections per mouse. **e**) 9 μm z-projection of layer 1 in the posterior parietal cortex showing MOBP-EGFP labeling of myelin sheaths (green) and THG-labeling of myelin sheaths and blood vessels (magenta). **f**) Zoom image of an isolated EGFP/THG-labeled myelin sheath from the white box in (**e**) excited with 1300 nm three-photon excitation. **g**) Isolated MOBP-EGFP sheath from layer 1 of a separate mouse visualized with 920 nm two-photon excitation combined with Spectral Confocal Reflectance (SCoRe) microscopy. Note the decreased THG/SCoRe labeling at putative nodes of Ranvier in (**f**) and (**g**) (arrowheads). **h**) 9 μm z-projection of a single mature oligodendrocyte in layer 1 from a third mouse showing EGFP-labeled processes connected to multiple THG-labeled sheaths. **i**) 5-pixel line intensity plot from the white line drawn in (**d**) showing relative fluorescence intensity of EGFP- and THG-labeled myelin sheaths and a THG-labeled blood vessel (arrowhead). All images pixel size = 0.36 μm. *p <0.05, box plots represent the median, interquartile ranges and minimum/maximum values

**Extended Data Fig. 3.**
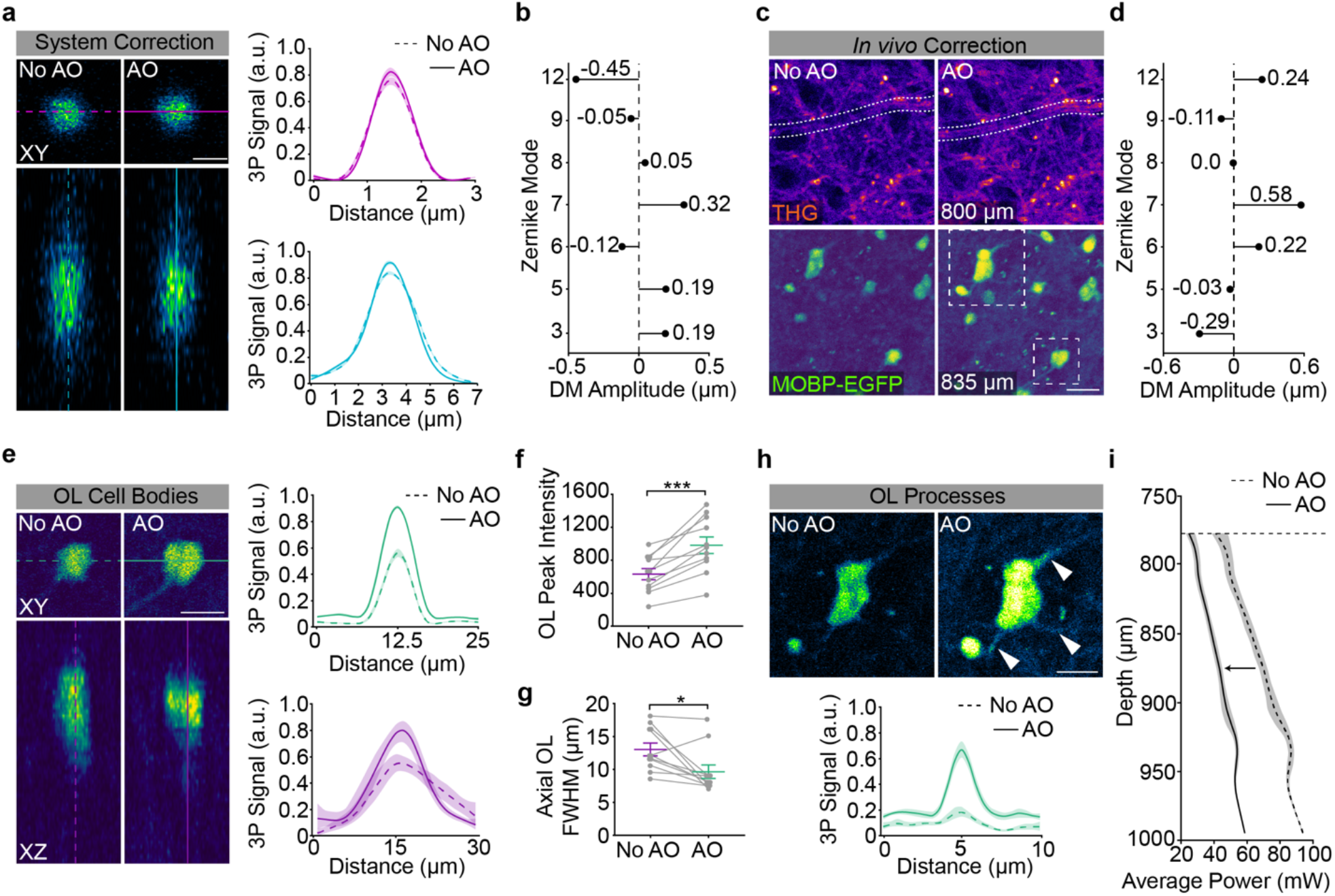
Adaptive optics improves SBR and axial resolution in the subcortical white matter. **a**) System adaptive optics (AO) correction with a two-dimensional 1.0 µm red polychromatic microsphere sample. Lateral (top) and axial (bottom) projections show a slight enhancement in peak signal after AO correction for system aberrations (8% increase, lateral; 9% increase, axial). The bead sample was diluted 1:10000, coverslipped with Prolong Gold, and the average power at the sample was <0.3 mW. System resolution = 0.88 µm (lateral) and 2.55 µm (axial) as calculated by the full width at half maximum of the gaussian fit after system AO correction. **b**) Deformable mirror (DM) amplitude plot showing the optimized stroke amplitudes (units = µm RMS) for each Zernike Mode in the system AO correction. **c**) *In vivo*, indirect, modal adaptive optics (AO) corrections were made at 775 - 825 µm depth (just above the white matter) by optimizing on the mean intensity of the third harmonic generation channel. The *in vivo* AO correction revealed cross sections through myelinated axons just dorsal to the corpus callosum (white dotted selection, top, 800 µm) and increased the mean intensity of EGFP-positive oligodendrocytes in the white matter ventral to the plane of the AO correction (bottom, 835 µm). Example images are single-slice images at the depth of the AO correction, 0.14 µm / pixel. **d**) Deformable mirror (DM) amplitude plot showing the optimized stroke amplitudes (units = µm RMS) for each Zernike Mode in the *in vivo* AO correction shown in (**c**). **e**) XY (top) and XZ (bottom) projections for a single EGFP-positive OL cell body from the example image shown in (**c**). The AO correction enhanced the normalized 3P signal by 68% (lateral) and 40% (axial). n = 2 mice, 11 OLs at > 800 µm depth. **f**) The peak intensity of the EGFP-oligodendrocyte signal was significantly enhanced by the AO correction (Paired, two-tailed Student’s t-test, p = 0.0002, n = 2 mice, 11 cells). **g**) The axial resolution of EGFP-oligodendrocyte cell bodies was significantly enhanced by the AO correction (Paired, two-tailed Student’s t-test, p = 0.014, n = 2 mice, 11 cells). **h**) AO correction enhances the peak intensity of OL processes by 202% (n = 2 mice, 4 processes). **i**) Average power vs. depth plot shows the average power at the sample through the depth of the white matter after AO correction (black, left), and the theoretical power curve (gray, right) necessary to maintain the same signal to background ratio without AO. Data are represented as individual points and mean ± SEM. Curves in **a, e**, and **h** represent the mean of the gaussian fits for each analyzed region (cell bodies in **a, e**, cell processes in **h**) with 95% confidence intervals. The intensity was integrated over a line plotted through the center of the cell structure (10 µm wide, cell bodies, 1 µm wide, processes). Zernike modes in (**b**), (**d**) = Spherical (12), Horizontal Trefoil (9), Horizontal Coma (8), Vertical Coma (7), Oblique Trefoil (6), Astigmatism (5), Oblique Astigmatism (3). For detailed statistics, see Supplementary Table 3.

**Extended Data Fig. 4.**
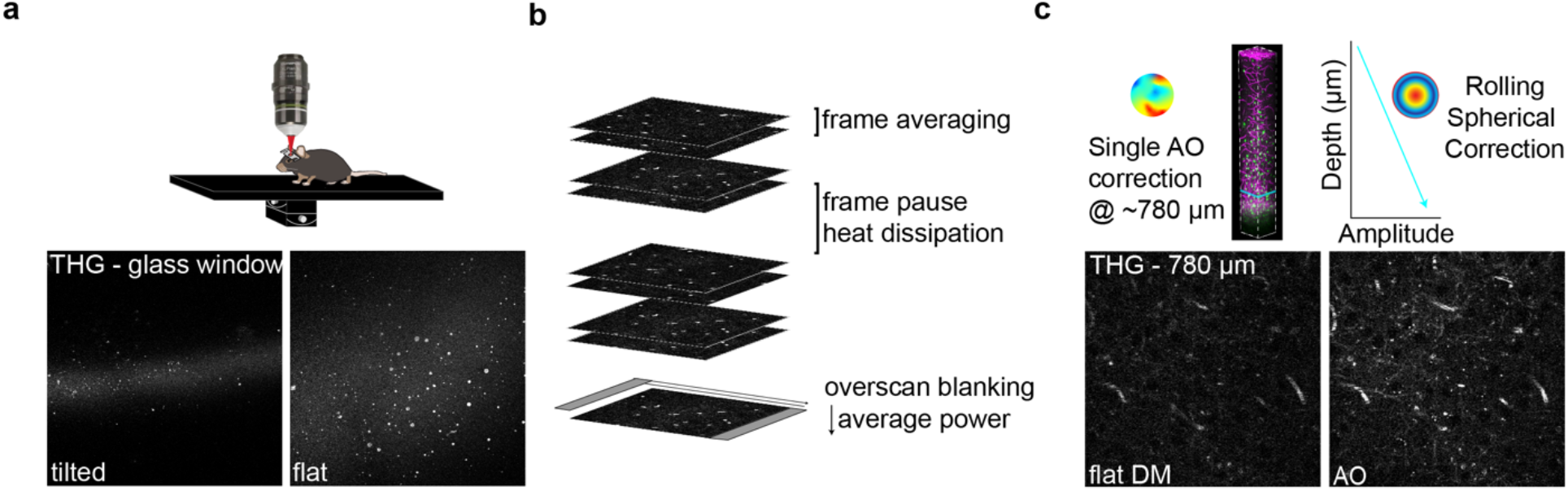
Mechanical, scanning, and optical modifications to increase 3P signal and decrease average power. **a**) The third harmonic generation (THG) signal at the surface of the cranial window was used to align the preparation orthogonally to the excitation light. **b**) Scanning modifications to increase SBR and decrease average power to mitigate risk of tissue damage. Frame averaging was advantageous compared to increasing the pixel dwell time to reduce risk of nonlinear damage. Z-stack acquisition was paused periodically to allow for heat dissipation (1 min. pause per 3 min. scanning). Laser pulses were blanked on the galvanometer overscan to reduce the average power applied to the preparation at each z-plane. **c**) Imaged-based AO correction increased SBR at depth (left) and modulating the spherical aberration correction linearly with depth improved SBR throughout the imaging volume (right). For longitudinal imaging, the AO correction was made before acquiring each time point at the same z-plane just above the scattering white matter (750-850 µm depth).

**Extended Data Fig. 5.**
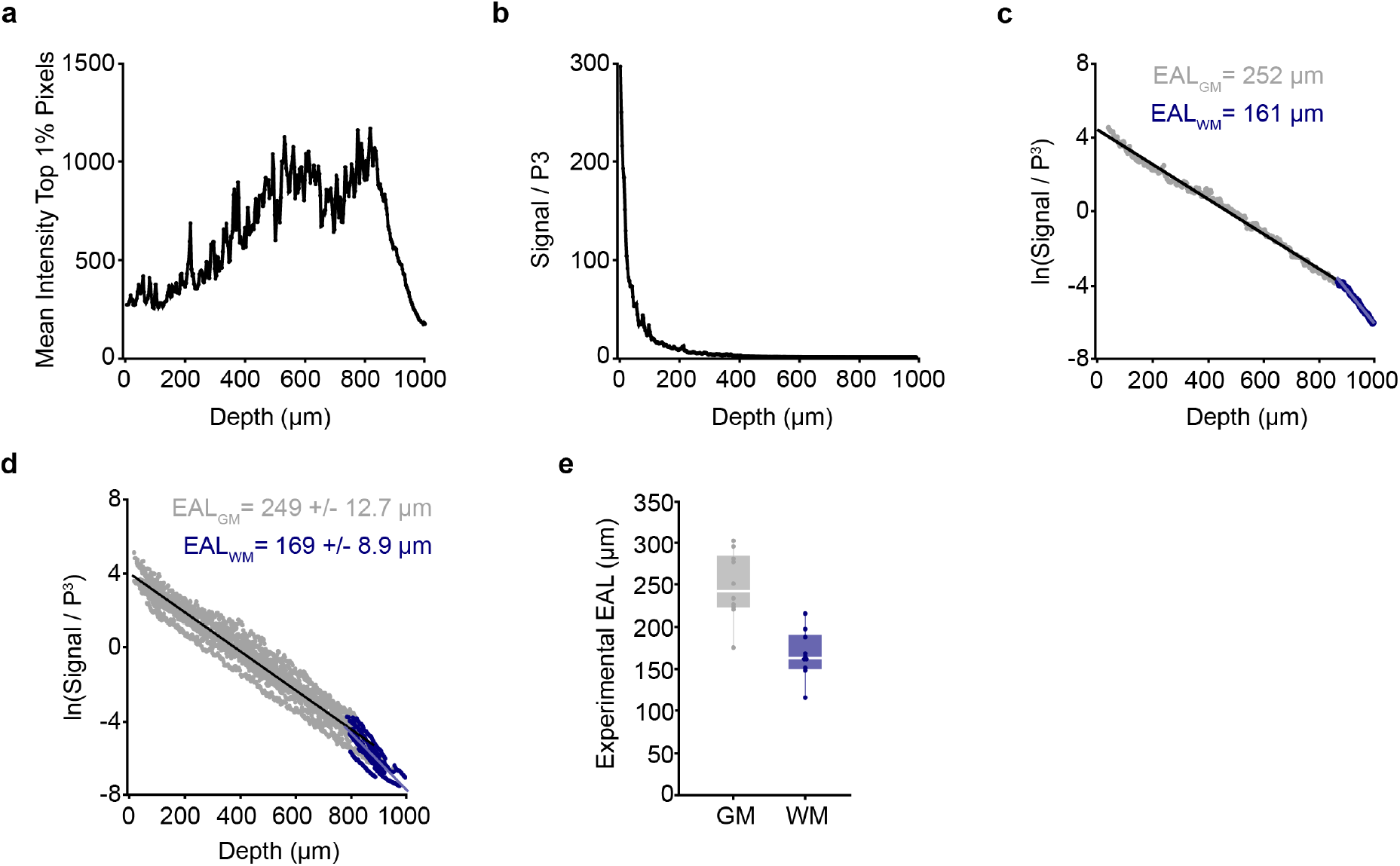
Calculation of mouse-specific effective attenuation lengths in the posterior parietal cortex. **a**) The mean intensity of the top 1% of the brightest pixels (experimental 3P signal) plotted with depth beneath the brain surface for an example mouse. **b**) Data from the mouse shown in (**a**) normalized to the cube of the pulse energy shows the decay of the 3P signal with depth in the mouse brain. **c**) Logarithm of the data in (**b**) shows a linear decrease with depth. The single mouse-specific effective attenuation length (EAL) can be calculated from the slope of the linear fit (gray = gray matter, blue = white matter). A steeper slope represents a shorter EAL due to increased scattering. **d**) Semilogarithmic plot showing gray matter and white matter EALs for n = 11 mice at the first time point of longitudinal imaging. Mean EAL (GM) = 249 +/- 12.7 µm, Mean EAL (WM) = 169 +/- 8.9 µm. **e**) Distributions of experimental mouse-specific EALs in the gray matter (gray) and white matter (blue). Linear fits (black line plots in **c,d**) were calculated separately by region. Box plots represent the median, interquartile ranges and minimum/maximum values.

**Extended Data Fig. 6.**
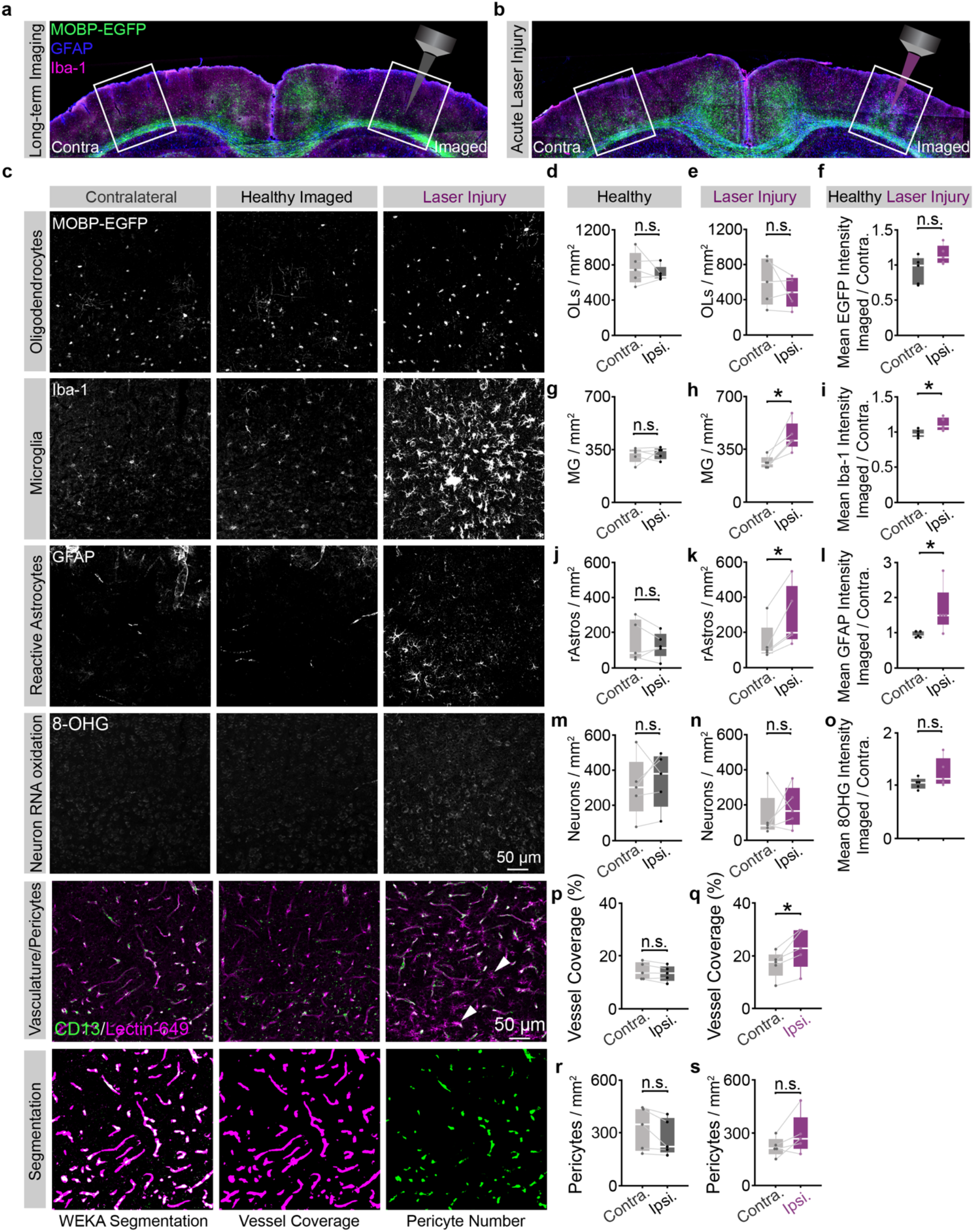
Optimized longitudinal three-photon imaging does not increase glial, neuronal, or vascular reactivity. **a**) Related to **Fig. 3**. Coronal brain section from the imaged (right) and contralateral (left) posterior parietal cortex (PPC) of a mouse that was perfused 24 hrs. following 10 weeks of chronic 3P imaging. **b**) Coronal brain section from the imaged (right) and contralateral (left) posterior parietal cortex (PPC) of a mouse that received supra-threshold excitation across the cortex and white matter to generate laser-induced positive control tissue (laser injury, right). **c**) High-resolution confocal images of the contralateral (left), longitudinal imaging field (middle) and laser-induced injury (right) regions in the deep cortical layers of the PPC, stained for transgenic MOBP-EGFP expression (mature oligodendrocytes), Iba-1 (microglia), GFAP (reactive astrocytes), 8-hydroxyguanosine (8-OHG, neuronal RNA oxidation), and CD13/Lectin-49 (pericytes, vasculature, respectively). All images were taken with identical power settings, processed with a 100 pixel rolling ball background subtraction and 0.7 pixel gaussian blur, and brightness/contrast correction was applied identically across channels. The bottom row shows an example of the trainable WEKA segmentation to measure pericytes (green) and vascular coverage (magenta, see Methods). Note the Lectin-positive microglia-like cells present in the laser injury vasculature image (white arrowheads). **d**) No difference in the density of MOBP-EGFP OLs between the contralateral and imaged regions in healthy mice. **e**) No difference in the density of MOBP-EGFP OLs between the contralateral and imaged regions in laser-induced injury tissue. **f**) The ratio of the EGFP mean intensity for the imaged:contralateral (contra.) hemispheres was ∼1 for healthy mice and did not change significantly in the positive control tissue. **g**) No difference in the density of Iba-1 microglia (MG) between the contralateral and imaged regions in healthy mice. **h**) Density of Iba-1 MG is significantly increased on the ipsilateral side of laser-induced injury tissue (Paired, two-tailed Student’s t-test, t(4) = 3.93, p = 0.016). **i**) The ratio of the imaged : contralateral Iba-1 mean intensity was ∼1 for healthy mice and significantly increased in positive controls (Unpaired, two-tailed Student’s t-test for equal variance, t(8) = 2.73, p = 0.026). **j**) No difference in the density of GFAP-positive reactive astrocytes (rAstros) between the contralateral and imaged regions in healthy mice. **k**) Density of GFAP+ rAstros is significantly increased on the damaged side in laser-induced injury tissue (Paired, two-tailed Student’s t-test, t(4) = 3.94, p = 0.017). **l**) The ratio of the imaged : contralateral (contra.) GFAP mean intensity was ∼1 for healthy mice and was significantly increased in positive controls (Wilcoxon rank sum test, Z = 2.09, p = 0.037). **m**) No difference in the density of 8-OHG-positive neurons after auto-thresholding (see Methods) between the contralateral and imaged regions in healthy mice. **n**) No difference in the density of 8-OHG-positive neurons between contralateral and imaged regions in laser-induced injury tissue. **o**) No difference in the imaged:contralateral ratio of 8-OHG mean intensities between healthy vs. laser-damaged mice. **p**) No difference in vessel coverage (% positive pixels) between the contralateral and imaged hemispheres of healthy multi-month longitudinal 3P imaging mice. **q**) Increase in vessel coverage (%) on the ipsilateral hemisphere of laser-induced injury positive control mice (Paired, two-tailed Student’s t-test, p = 0.022). **r**) No difference in the density of pericytes between the contra. and ipsi. hemispheres of healthy long-term 3P mice. **s**) No difference in the density of pericytes between the contra. and ipsi. hemispheres of laser-induced injury mice. *p < 0.05, **p < 0.01, n.s., not significant; n = 5 mice (healthy longitudinal), 5 mice (laser-induced injury), 2 sections, 4 hemispheres per condition. Box plots represent the median, interquartile ranges and minimum/maximum values. For detailed statistics, see Supplementary Table 3.

**Extended Data Fig. 7.**
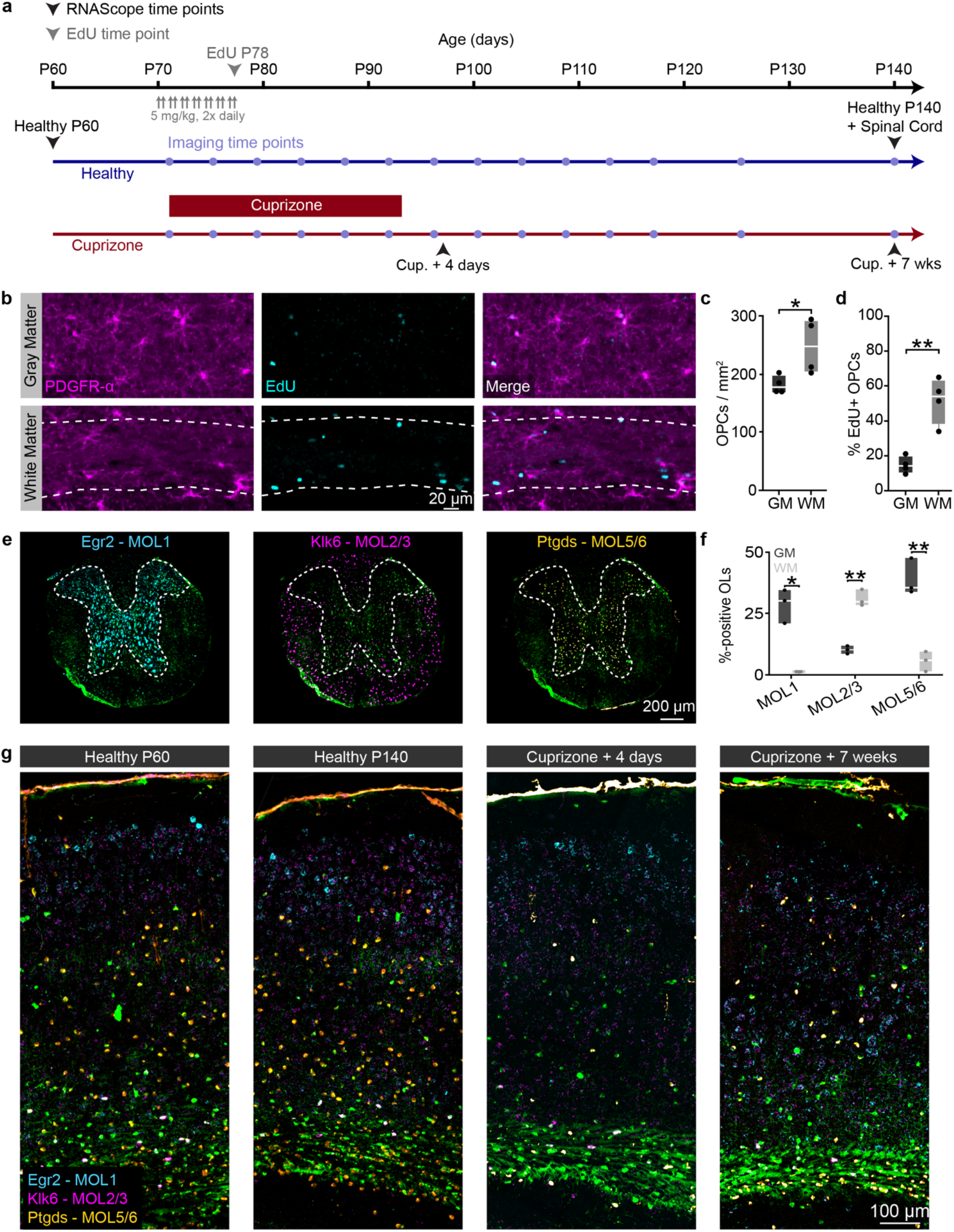
Immunofluorescent and *in situ* hybridization techniques allow for probing OPC proliferation and oligodendrocyte subpopulations in the cortex and subcortical white matter. **a**) Experimental timeline of tissue collection for EdU and RNAScope analyses in healthy and cuprizone mice. Healthy MOBP-EGFP mice were injected with 5 mg/kg of the thymidine analog EdU twice a day (10-12 hr. interval) for seven days starting at P70 and perfused the following day. Healthy MOBP-EGFP mice were perfused at P60 and P140 to assess aging-dependent changes in adult oligodendrocyte subpopulations. MOBP-EGFP mice that were fed 0.2% cuprizone for three weeks were perfused at 4 days post-cuprizone removal (peak demyelination, matched to *in vivo* imaging data, **Fig. 4**), and 7 weeks post-cuprizone removal (regeneration, matched to the final time point of *in vivo* imaging, see timeline, **Fig. 4**). **b**) EdU-positive proliferated OPCs in the posterior parietal cortex (top, GM) and subcortical white matter (bottom, WM, dashed border). **c**) The density of PDGFR-α-positive OPCs was significantly increased in the WM compared to the GM (248.6 ± 23.8 vs. 181.2 ± 7.7, Unpaired, two-tailed Student’s t-test for equal variance, t(6) = 2.69, p = 0.036). **d**) The percentage PDGFR-α+/EdU+ OPCs is increased in the WM compared to the GM (Unpaired, two-tailed Student’s t-test for equal variance, 51.6 ± 6.6 vs. 14.6 ± 2.5, t(6) = 5.29, p = 0.002). **e**) Coronal sections from the mid-thoracic spinal cord of healthy MOBP-EGFP mice were taken at P140 and run in parallel with brain sections to confirm the labeling specificity of our oligodendrocyte subpopulation probe-set (Egr2, MOL1, cyan; Klk6, MOL2/3, magenta; Ptgds, MOL5/6, orange). **f**) Egr2 and Ptgds preferentially label oligodendrocytes in the spinal gray matter, while Klk6 predominantly labels oligodendrocytes in the spinal white matter (Unpaired, two-tailed students t-test for unequal variance). **g**) Coronal sections from the posterior parietal cortex of MOBP-EGFP mice (−1.7 to -2.3 mm posterior and 1 to 3 mm lateral to bregma) showing the pattern of OL subpopulation labeling at the experimental time points described in (**a**). *p < 0.05, **p<0.01, n = 4 mice, 4 sections per mouse. Boxplots represent the median, interquartile range and the minimum and maximum values. For detailed statistics, see Supplementary Table 3.

**Extended Data Fig. 8.**
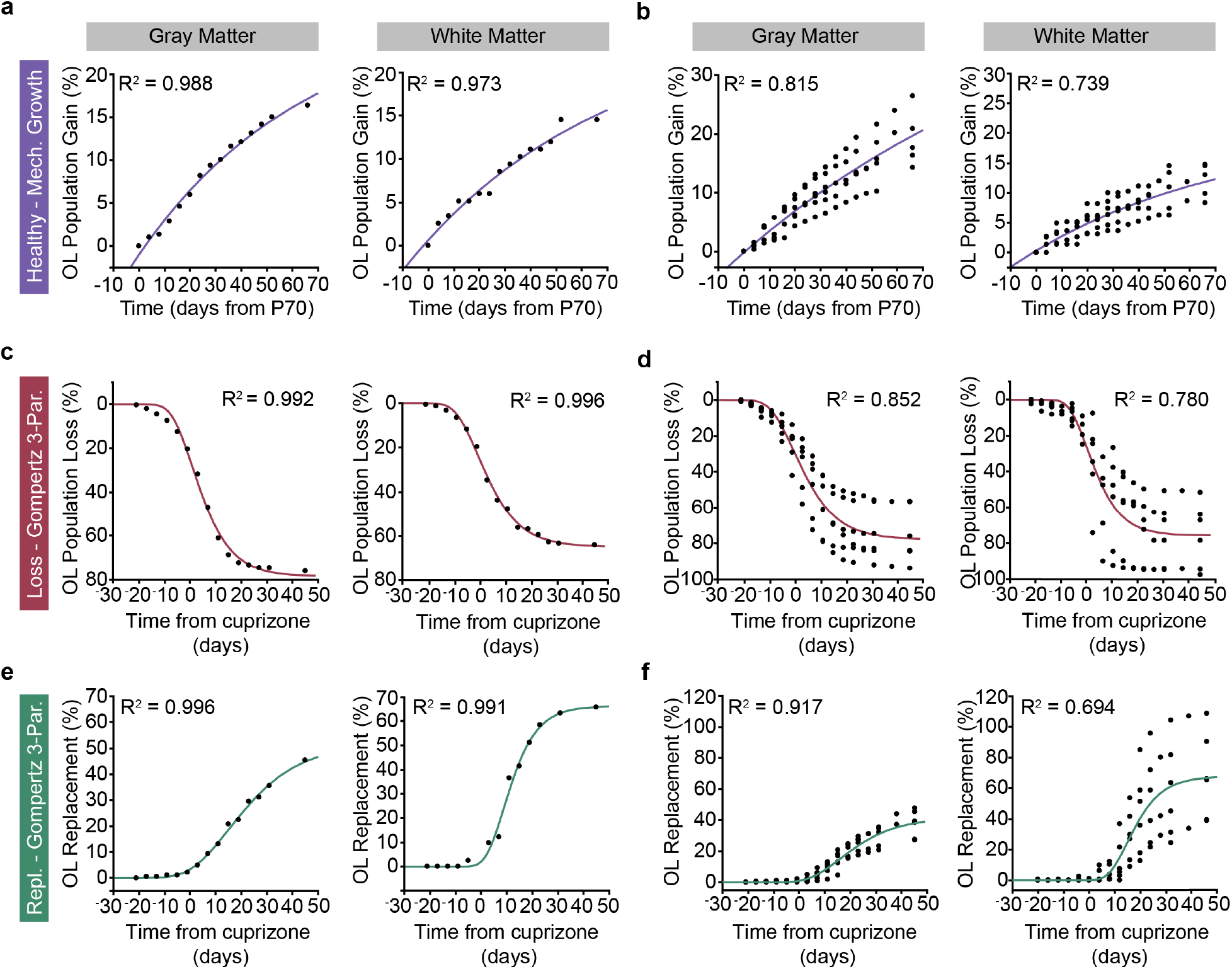
Modeling oligodendrocyte growth, loss, and regeneration in adult mouse cortex and white matter. **a-b**) Cumulative oligodendrocyte population growth (% gain over time) in the healthy brain was modeled using asymptote-restricted exponential mechanistic growth curve-fitting. **a**) Cumulative OL gain (%) and mechanistic growth fit in the gray matter (left) and the white matter (right) for an example mouse. **b**) Cumulative OL gain (%) and mechanistic growth fits in the gray matter (left) and the white matter (right) for the healthy group (n = 6 mice). **c-d**) Cumulative oligodendrocyte population loss (% loss over time) due to cuprizone administration was modeled using three-parameter Gompertz Sigmoid curve-fitting. **c**) Cumulative OL loss (%) and three-parameter Gompertz curves in the gray matter (left) and the white matter (right) for an example mouse. **d**) Cumulative OL loss (%) and three-parameter Gompertz curves in the gray matter (left) and the white matter (right) for the cuprizone de/remyelination group (n = 6 mice). **e-f**) Cumulative oligodendrocyte population regeneration (% cell replacement over time) following cuprizone cessation was modeled using three-parameter Gompertz Sigmoid curve-fitting. **e**) Cumulative OL replacement (%) and three-parameter Gompertz curves in the gray matter (left) and the white matter (right) for an example mouse. **f**) Cumulative OL replacement (%) and three-paramter Gompertz curves in the gray matter (left) and the white matter (right) for the cuprizone de/remyelination group (n = 6 mice). Modeled rate and timing metrics in **Main Figs. 4-7** were calculated by fitting curves to data from individual mice and then extracting summary data (e.g. timing of inflection point).

**Extended Data Fig. 9.**
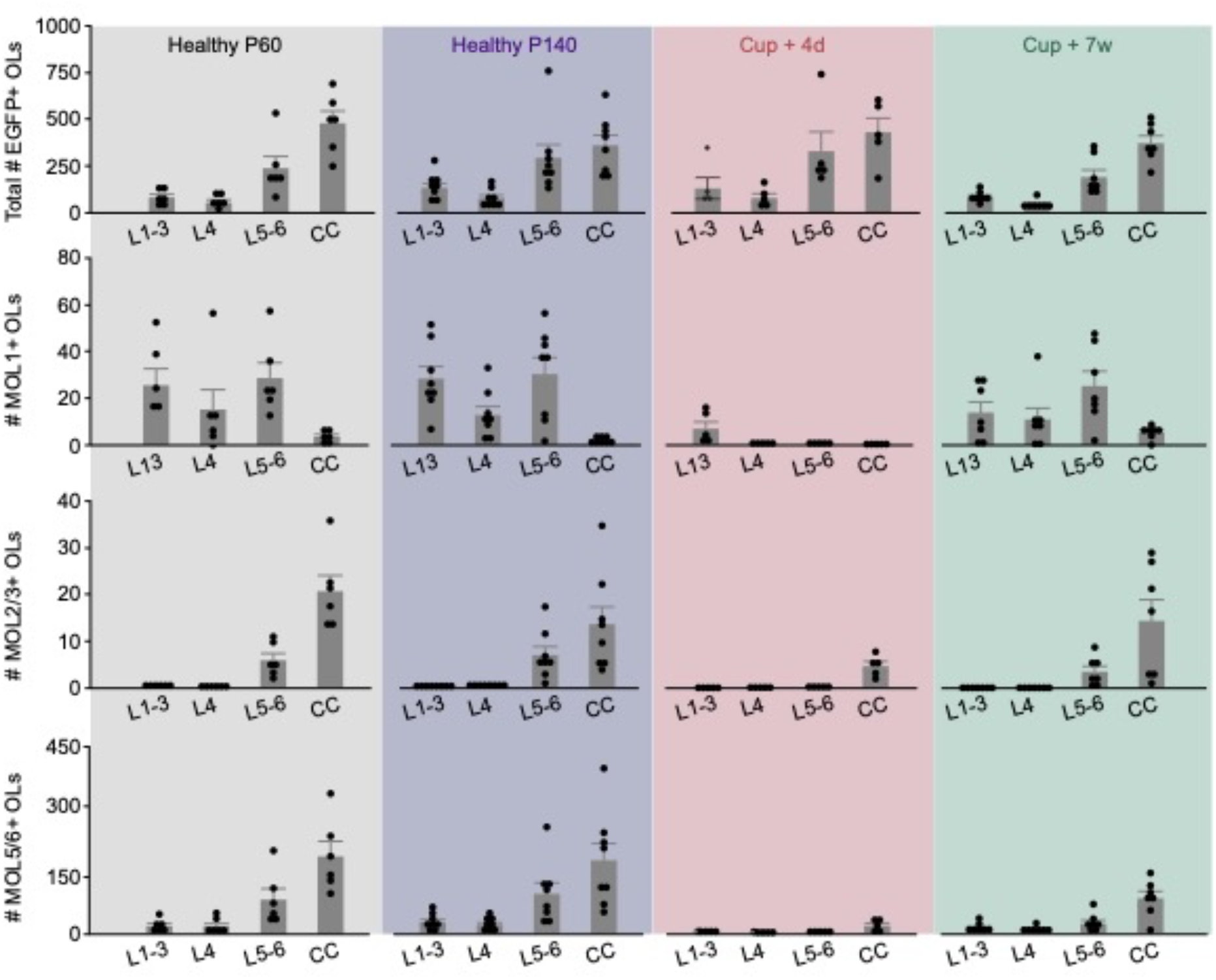
Layer and region-dependent proportions of MOL1, MOL2/3, and MOL5/6 across healthy aging and cuprizone treatment. Related to **Figs. 7,8**. Raw counts of the total # of segmented MOBP-EGFP OLs (top), MOL1+ OLs (row 2), MOL2/3+ OLs (row 3) and MOL5/6+ OLs (bottom) across cortical and subcortical layers (x-axes) and experimental time points (shaded background colors). Data presented in the main figures were expressed either as the percentage of total OLs for each probe (**Figs. 7-8**), or the change in proportion of each marker from the healthy P140 time point (**Figs. 4-6**).

**Extended Data Fig. 10.**
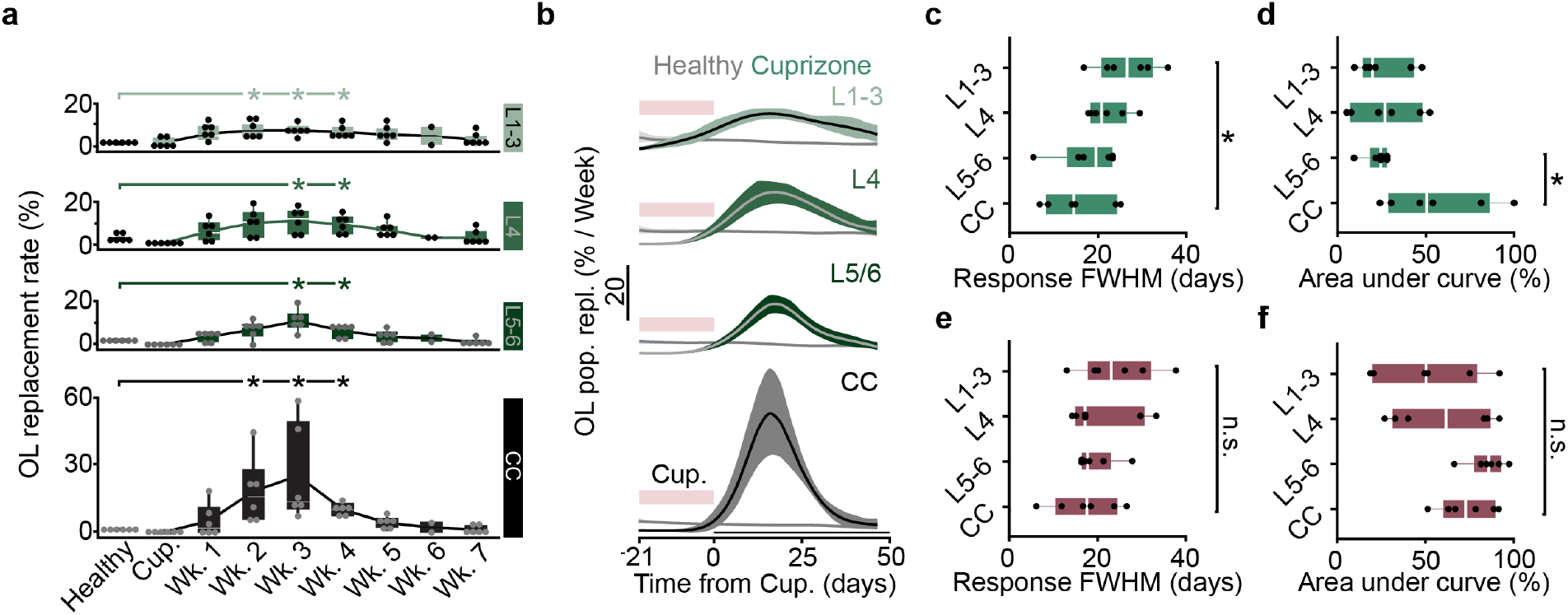
Layer-dependent differences in the temporal dynamics of cuprizone induced loss and regeneration. **a-e**) Related to Fig. 7. **a**) OL population replacement (%) plotted over weekly time bins following cuprizone cessation for each anatomically defined region. Replacement rate is increased from healthy baseline from 2-4 weeks post-cuprizone in L1-3 and CC, and 3-4 weeks post-cuprizone in L4 and L5/6 (Kruskal-Wallis test followed by Steel method for comparison with control, p = 0.034 for L1-3, Weeks 2-4 vs. Healthy; p = 0.034 for L4 Weeks 3-4 vs. Healthy; p = 0.034 for L5-6, Weeks 3-4 vs. Healthy; p = 0.034 for CC Weeks 2-4 vs. Healthy). **b**) First-derivative of the modeled growth curves for healthy baseline and OL replacement (Mechanistic Growth, Gompertz 3P, respectively) showing the response duration of regeneration across regions. **c**) The regeneration response duration, calculated as the full width at half maximum of the rate curves in (**b**), is significantly longer in the superficial layers 1-3 compared to the CC (One-way ANOVA followed by Tukey’s HSD, p = 0.048). **d**) The magnitude of the regeneration response, as calculated by the area under the curve above the healthy baseline rate, is significantly suppressed in L5-6 compared to the CC (One-way ANOVA followed by Tukey’s HSD, p = 0.037). **e-f)** No significant differences in the duration or magnitude of the cuprizone cell loss response across regions. Data in (**a)** were calculated based on the raw % change in cell population from baseline. Data in **b-f** were derived from the modeled and scaled growth curves. *p < 0.05, **p < 0.01, n.s. = not significant; cumulative growth curves represent cubic splines with 95% confidence intervals, box plots represent the median, interquartile ranges and minimum/maximum values. For detailed statistics, see Supplementary Table 3.

