## Supplemental Tables for "Long-term *in vivo* three-photon imaging reveals region-specific differences in healthy and regenerative oligodendrogenesis"

### Supplementary Table 1 | Histological to *in vivo* conversion of layer widths

Related to Figs. 7,8 and Extended Data Figs. 7,9,10

| Cortical Layer | Z-depth through PPC (Allen Brain Atlas) - Top (μm) | Z-depth through PPC (Allen Brain Atlas) - Bottom (μm) | Layer Widths (μm) | Multiplication Factor (Layer Width / 871 μm) |
|---|---|---|---|---|
| 1 | 0 | 130 | 130 | 0.149 |
| 2-3 | 130 | 378 | 248 | 0.285 |
| 4 | 378 | 492 | 114 | 0.131 |
| 5-6 | 492 | 871 | 379 | 0.435 |

Lein, E. S. *et al.* Genome-wide atlas of gene expression in the adult mouse brain. *Nature* **445**, 168–176 (2007).

Supplementary Table 2 | Antibody specifications

| | Antigen | Host | Time of L.A.B. Treatment | Source | Antibody # | Dilution |
|---|---|---|---|---|---|---|
| Primary Antibodies | Anti-Enhanced Green Fluorescent Protein (EGFP) | Chicken | NA | Aves | GFP1020 | 1:1000 |
| | Anti-Enhanced Green Fluorescent Protein (EGFP) | Goat | NA | SciGen | AB0066 | 1:2000 |
| | Anti-Glial Fibrillary Acidic Protein (GFAP) | Mouse IgG1 | NA | Sigma | G3893-GA5 | 1:1000 |
| | Anti-Heat Shock Proteins 70/72 (HSP70/72) | Mouse IgG1 | 7 min. | Enzo | ADI-SPA-810-D | 1:400 |
| | Anti-Ionized Calcium-Binding Adapter Molecule 1 (Iba-1) | Rabbit | NA | Wako | 019-19741 | 1:1000 |
| | Anti-H2A Histone Family Member X (γ-H2AX) | Rabbit | NA | Cell Signaling | S139-20E3 | 1:500 |
| | Anti-8-hydroxyguanosine (8-OHG) | Mouse IgG2b | NA | Fisher | MA110602-15A3 | 1:1500 |
| | Anti-Platelet-Derived Growth Factor Receptor - Alpha (Pdgfr-α) | Goat | NA | R&D Biosystems | AF1062 | 1:2000 |
| | Anti-aspartoacyclase (ASPA) | Rabbit | 5 min. | Genetex | ABN1698 | 1:1000 |
| | Anti-neuronal nuclear protein (NeuN) | Chicken | NA | Millipore | ABN91 | 1:500 |
| | Anti pan-Neurofilament (SMI-312) | Mouse IgG1 | NA | BioLegend | NC1239357 | 1:250 |
| | Anti-vesicular glutamate transporter 2 (Vglut2) | Guinea Pig | NA | EMD Millipore | AB2251-I | 1:1000 |
| | **Antigen** | **Host** | **Time of L.A.B. Treatment** | **Source** | **Antibody #** | **Dilution** |
| Secondary Antibodies | anti-Rabbit 405 | Donkey | see above for antibody pairings | Invitrogen | A48258 | 1:500 |
| | anti-Chicken 405 | Donkey | | Jackson | 703-475-155 | 1:300 |
| | anti-Chicken 488 | Donkey | | Jackson | 703-545-155 | 1:500 |
| | anti-Goat 488 | Donkey | | Jackson | 705-545-003 | 1:500 |
| | anti-Goat 546 | Donkey | | Jackson | A11056 | 1:500 |
| | anti-Rabbit Cy3 | Donkey | | Jackson | 711-165-152 | 1:500 |
| | anti-Mouse IgG2b 570 | Goat | | Jackson | 115-297-187 | 1:500 |
| | anti-Guinea Pig 568 | Goat | | Invitrogen | A11075 | 1:500 |
| | anti-Mouse IgG1 647 | Goat | | Jackson | 115-607-185 | 1:500 |

### Supplementary Table 3 | Statistics results table

*Statistics results are sorted by figure, and both main figures and supplementary figures are listed in the order they appear in the text.*

**Statistically significant comparisons are bolded**

#### Table 3.1 | Statistics for Figure 1 and associated extended data (Extended Data Figs. 2,3)

| Fig. 1 | Measure | Values | N | Statistical test | Significance |
|---|---|---|---|---|---|
| **Fig. 1c** | Signal to background ratio with two-photon vs. three-photon imaging at different brain depths | 0-100 µm: 55.400 ± 12.439 (two-photon) vs. 20.889 ± 5.124 (three-photon) | n=3-4 two-photon mice, 50 - 74 oligodendrocytes analyzed per mouse, n=4 three-photon mice, 101 - 123 analyzed oligodendrocytes per mouse. | Shapiro-Wilk W Test on distributions of imaging modality and depth not significant. Depth pairs from different mice compared piecewise with F-test followed by unpaired two-tailed student's t-Test for equal or unequal variance. | **0-100 µm: t(6) = -2.565, p=0.043** |
| | | 101-400 µm: no significant differences. | | | 101-400 µm: no significant differences. |
| | | 401-500 µm: 7.259 ± 3.085 (two-photon) vs. 21.840 ± 3.935 (three-photon) | | | **401-500 µm: t(6) = 2.916, p=0.027** |
| | | 501-600 µm: 4.602 ± 1.580 (two-photon) vs. 19.662 ± 2.775 (three-photon) | | | **501-600 µm: t(6) = 4.716, p=.003** |
| | | 601-700 µm: 3.667± 0.353 (two-photon) vs. 17.876 ± 3.206 (three-photon) | | | **601-700 µm: t(3.072) = 4.406, p = 0.021** |
| | | 701-1100 µm: insufficient two-photon data for comparison | | | |
| **Fig. 1d** | Number of oligodendrocyte cell bodies > 2 SBR per imaging volume with two-photon vs. three-photon, single comparison at layer 5 | Layer 5: 41.362 ± 10.650 vs. 78.753 ± 10.153. | n=4 two-photon mice, n=8 three-photon mice. | Shapiro-Wilk W Test for normality by imaging modality and depth not significant. Single comparison at layer 5 using F-Test followed by unpaired two-tailed student's t-Test for equal variance. | **t(10) = 2.286, p=0.045** |
| Fig. 1f | Orientation of THG-positive fibers in the corpus callosum vs. alveus. | 15.27 ± 4.942 (corpus callosum) vs. -0.403 ± 1.041 (alveus) degrees | n=41 cells from 3 mice pre-learning, and n=29 cells from 3 mice post-learning | Shapiro-Wilk W Test not significant. F-Test followed by unpaired two-tailed student's t-Test for unequal variance. | **t(5.573) = -3.244, p=0.026** |
| **Ext. Data Fig. 2** | **Measure** | **Values** | **N** | **Statistical test** | **Significance** |
| Fig. 2d | Percentage of MOBP+ cells that also express ASPA | 95.6 ± 0.75% (cortex) vs. 86.5 ± 2.63% (white matter) | n=4 mice, 2 sections per mouse | Shapiro-Wilk W Test not significant. F-Test followed by unpaired two-tailed student's t-Test for equal variance. | **t(6) = -3.32, p = 0.016** |
| **Ext. Data Fig. 3** | **Measure** | **Values** | **N** | **Statistical test** | **Significance** |
| Fig. 3f | Lateral intensity of oligodendrocyte cell bodies with and without Adaptive Optics | 631.77 ± 68.15 vs. 983.34 ± 100.95 a.u. | n = 2 mice, 11 oligodenrocytes at >800 µm depth | Shapiro-Wilk W Test not significant. Paired one-sample t-Test, two-tailed. | **p = 0.0002** |
| Fig. 3g | Axial intensity of oligodendrocyte cell bodies with and without Adaptive Optics | 12.96 ± 0.97 vs. 9.59 ± 1.03 µm | n = 2 mice, 11 oligodenrocytes at >800 µm depth | Shapiro-Wilk W Test not significant. Paired one-sample t-Test, two-tailed. | **p = 0.014** |

Supplementary Table 3 | Statistics results table

*Statistics results are sorted by figure, and both main figures and supplementary figures are listed in the order they appear in the text.*

**Statistically significant comparisons are bolded**

Table 3.2 | Statistics for Figure 3 and associated extended data (Extended Data Fig. 6)

| Fig. 3 | Measure | Values | N | Statistical test | Significance |
|---|---|---|---|---|---|
| **Fig. 3e** | # of HSP70/72-positive oligodendrocytes per mm$^2$ (healthy long-term imaging) | 20.952±12.67 (contralateral) vs. 27.73±11.72 (ipsilateral) | n = 5 mice, 2 sections / 4 hemispheres per mouse | Shapiro-Wilk W Test not significant. Paired one-sample t-Test, two-tailed. | p = 0.551 |
| **Fig. 3f** | # of HSP70/72-positive oligodendrocytes per mm$^2$ (laser-induced injury) | 39.2±18.6 (contralateral) vs. 71.7±22.6 (ipsilateral) | n = 5 mice, 2 sections / 4 hemispheres per mouse | Shapiro-Wilk W Test not significant. Paired one-sample t-Test, two-tailed. | **p = 0.041** |
| **Fig. 3g** | Ratio of the ipsilateral : contralateral normalized HSP70/72 fluorescence intensity | Mean intensity (imaged) / Mean intensity (contralateral). 0.91±0.08 (Healthy) vs. 5.26±2.46 (laser-injury) | n = 5 mice (healthy), n= 5 mice (laser-injury) 2 sections / 4 hemispheres per mouse | Shapiro-Wilk W Test significant. Wilcoxon rank sum test. | **z = 2.089, p = 0.037** |
| **Fig. 3h** | # of γ-H2A.X-positive oligodendrocytes per mm$^2$ (healthy long-term imaging) | 21.94±7.59 (contralateral) vs. 36.45±12.39 (ipsilateral) γ-H2A.X-positive oligodendrocytes per mm2 | n = 5 mice, 2 sections / 4 hemispheres per mouse | Shapiro-Wilk W Test not significant. Paired one-sample t-Test, two-tailed. | t(4) = 1.36, p = 0.244 |
| **Fig. 3i** | # of γ-H2A.X-positive oligodendrocytes per mm$^2$ (laser-induced injury) | 19.69±9.98 (contralateral) vs. 35.306±13.28 (ipsilateral) γ-H2A.X-positive oligodendrocytes per mm2 | n = 5 mice, 2 sections / 4 hemispheres per mouse | Shapiro-Wilk W Test not significant. Paired one-sample t-Test, two-tailed. | t(4) = 1.41, p = 0.231 |
| **Fig. 3j** | Ratio of the ipsilateral : contralateral normalized γ-H2A.X- fluorescence intensity | Mean intensity (imaged) / Mean intensity (contralateral). 0.94±0.04 (Healthy) vs. 2.39±0.68 (laser-injury) | n = 5 mice (healthy), n= 5 mice (laser-injury) 2 sections / 4 hemispheres per mouse | Shapiro-Wilk W Test significant. Wilcoxon rank sum test. | z = 1.462, p = 0.144 |
| **Fig. 3l** | Mean rate of healthy oligodendrocyte gain per week (Two-photon vs. Three-photon) | Two-photon = 1.7±0.2 vs. Three-photon = 1.4±0.3% gained per week | n=5 mice, 477 oligodendrocytes (Two-photon); n=4 mice, 340 oligodendrocytes (Three-photon) | Shapiro-Wilk W Test not significant. F-Test followed by one-way, two-tailed unpaired Student's t-Test. | t(6) = 0.819, p = 0.440 |

| Ext. Data Fig. 6 | Measure | Values | N | Statistical test | Significance |
|---|---|---|---|---|---|
| **Fig. 6d** | # of MOBP-EGFP-positive oligodendrocytes per mm$^2$ (healthy long-term imaging) | 765.02±83.00 (contralateral) vs. 703.67±38.55 (ipsilateral) | n = 5 mice, 2 sections / 4 hemispheres per mouse | Shapiro-Wilk W Test significant. Wilcoxon signed rank test. | Prob > \|S\| = 1.00 |
| **Fig. 6e** | # of MOBP-EGFP-positive oligodendrocytes per mm$^2$ (laser-induced injury) | 610.60±119.3 (contralateral) vs. 488.03±75.64 (ipsilateral) | n = 5 mice, 2 sections / 4 hemispheres per mouse | Shapiro-Wilk W Test not significant. Paired one-sample t-Test, two-tailed. | t(4) = -1.16, p = 0.311 |
| **Fig. 6f** | Ratio of the ipsilateral : contralateral normalized MOBP-EGFP fluorescence intensity | Mean intensity (imaged) / Mean intensity (contralateral). 0.92±0.09 (Healthy) vs. 1.14±0.06 (laser-injury) | n = 5 mice (healthy), n= 5 mice (laser-injury) 2 sections / 4 hemispheres per mouse | Shapiro-Wilk W Test not significant. F-Test followed by unpaired two-tailed student's t-Test for equal variance. | t(4) = 2.04, p = 0.076 |
| **Fig. 6g** | # of Iba-1-positive microglia per mm$^2$ (healthy long-term imaging) | 312.19±22.16 (contralateral) vs. 325.39±17.45 (ipsilateral) | n = 5 mice, 2 sections / 4 hemispheres per mouse | Shapiro-Wilk W Test not significant. Paired one-sample t-Test, two-tailed. | t(4) = 0.48, p = 0.658 |
| **Fig. 6h** | # of Iba-1-positive microglia per mm$^2$ (laser-induced injury) | 262.2±18.3 (contralateral) vs. 440.7±45.6 (ipsilateral) | n = 5 mice, 2 sections / 4 hemispheres per mouse | Shapiro-Wilk W Test not significant. Paired one-sample t-Test, two-tailed. | **t(4) = 3.99, p = 0.016** |
| **Fig. 6i** | Ratio of the ipsilateral : contralateral normalized Iba-1 fluorescence intensity | Mean intensity (imaged) / Mean intensity (contralateral). 0.98±0.03 (Healthy) vs. 1.11±0.04 (laser-injury) | n = 5 mice (healthy), n= 5 mice (laser-injury) 2 sections / 4 hemispheres per mouse | Shapiro-Wilk W Test not significant. F-Test followed by unpaired two-tailed student's t-Test for equal variance. | **t(8) = 2.73, p = 0.026** |
| **Fig. 6j** | # of GFAP-positive astrocytes per mm$^2$ (healthy long-term imaging) | 148.08±52.70 (contralateral) vs. 126.49±32.99 (ipsilateral) | n = 5 mice, 2 sections / 4 hemispheres per mouse | Shapiro-Wilk W Test not significant. Paired one-sample t-Test, two-tailed. | t(4) = -0.73, p = 0.504 |
| **Fig. 6k** | # of GFAP-positive astrocytes per mm$^2$ (laser-induced injury) | 142.4±49.7 (contralateral) vs. 290.8±76.4 (ipsilateral) | n = 5 mice, 2 sections / 4 hemispheres per mouse | Shapiro-Wilk W Test significant. Wilcoxon signed rank test. | **t(4) = 3.94, p = 0.017** |
| **Fig. 6l** | Ratio of the ipsilateral : contralateral normalized GFAP fluorescence intensity | Mean intensity (imaged) / Mean intensity (contralateral). 1.04±0.05 (contralateral) vs. 1.24±0.27 (ipsilateral) | n = 5 mice (healthy), n= 5 mice (laser-injury) 2 sections / 4 hemispheres per mouse | Shapiro-Wilk W Test significant. Wilcoxon rank sum test. | **z = 2.09, p = 0.037** |
| **Fig. 6m** | # of 8-OHG-positive neurons greater than 90% pixel threshold per mm$^2$ (healthy long-term imaging) | 303.72±77.31 (contralateral) vs. 342.39±69.71 (ipsilateral) | n = 5 mice, 2 sections / 4 hemispheres per mouse | Shapiro-Wilk W Test not significant. Paired one-sample t-Test, two-tailed. | t(4) = 0.62, p = 0.572 |
| **Fig. 6n** | # of 8-OHG-positive neurons greater than 90% pixel threshold per mm$^2$ (laser-induced injury) | 137.14±61.38 (contralateral) vs. 187.65±51.03 (ipsilateral) | n = 5 mice, 2 sections / 4 hemispheres per mouse | Shapiro-Wilk W Test significant. Wilcoxon signed rank test. | t(4) = 0.59, p = 0.589 |
| **Fig. 6o** | Ratio of the ipsilateral : contralateral normalized 8-OHG fluorescence intensity | Mean intensity (imaged) / Mean intensity (contralateral). 0.93±0.04 (contralateral) vs. 1.63±0.30 (ipsilateral) | n = 5 mice (healthy), n= 5 mice (laser-injury) 2 sections / 4 hemispheres per mouse | Shapiro-Wilk W Test not significant. F-Test followed by unpaired two-tailed student's t-Test for equal variance. | t(8) = 1.57, p = 0.154 |
| **Fig. 6p** | Percentage of vascular coverage (segmented Lectin-649 positive area, healthy long-term imaging) | 14.22±1.43 (contralateral) vs. 13.30±1.30% (ipsilateral) of image area | n = 5 mice, 2 sections / 4 hemispheres per mouse | Shapiro-Wilk W Test not significant. Paired one-sample t-Test, two-tailed. | t(4) = -1.91, p = 0.13 |
| **Fig. 6q** | Percentage of vascular coverage (segmented Lectin-649 positive area, laser-induced injury) | 16.8±2.3 (contralateral) vs. 22.8±3.4% (ipsilateral) of image area | n = 5 mice, 2 sections / 4 hemispheres per mouse | Shapiro-Wilk W Test not significant. Paired one-sample t-Test, two-tailed. | **t(4) = 3.65, p = 0.022** |
| **Fig. 6r** | # of CD13-positive pericytes per mm$^2$ (healthy long-term imaging) | 320.94±53.36 (contralateral) vs. 272.21±46.27 (ipsilateral) | n = 5 mice, 2 sections / 4 hemispheres per mouse | Shapiro-Wilk W Test not significant. Paired one-sample t-Test, two-tailed. | t(4) = -2.02, p = 0.114 |
| **Fig. 6s** | # of CD13-positive pericytes per mm$^2$ (laser-induced injury) | 218.60±23.67 (contralateral) vs. 300.77±51.11 (ipsilateral) | n = 5 mice, 2 sections / 4 hemispheres per mouse | Shapiro-Wilk W Test not significant. Paired one-sample t-Test, two-tailed. | t(4) = 2.24, p = 0.088 |

Supplementary Table 3 | Statistics results table

*Statistics results are sorted by figure, and both main figures and supplementary figures are listed in the order they appear in the text.*

**Statistically significant comparisons are bolded**

Table 3.3 | Statistics for Figure 4 and associated extended data (Extended Data Fig. 7)

| Fig. 4 | Measure | Values | N | Statistical test | Significance |
|---|---|---|---|---|---|
| **Fig. 4e** | Total # new oligodendrocytes / 350 x 350 x 60 µm volume at 66 day time point (Gray Matter vs. White Matter) | 33.8±5.5 vs. 10.9±1.3 OLs / imaging volume | n = 5 mice | Shapiro-Wilk W Test not significant. F-Test followed by one-way, two-tailed unpaired Student's t-Test for unequal variance | **t(4.45) = 4.03, p = 0.013** |
| | Rate of new oligodendrocytes gained / Day / 350 x 350 x 60 µm volume (Gray Matter vs. White Matter) | 3.9±0.5 vs. 1.2±0.2 OLs / volume / week | n = 6 mice | Shapiro-Wilk W Test not significant. F-Test followed by one-way, two-tailed unpaired Student's t-Test for unequal variance | **t(6.86) = 5.14, p = 0.002** |
| **Fig. 4g** | Total % Oligodendrocyte Gain at 66 day time point (Gray Matter vs. White Matter) | 19.1±2.1% vs. 12.1±1.3% | n = 5 mice | Shapiro-Wilk W Test not significant. F-Test followed by one-way, two-tailed unpaired Student's t-Test for equal variance | **t(8) = -2.83, p = 0.022** |
| **Fig. 4h** | % Rate of Oligodendrocyte Gain (Gray Matter vs. White Matter) at 5 week time point calculated from Mechanistic Growth Curves | 2.3±0.3 vs. 1.3±0.2% per week | n = 6 mice | Shapiro-Wilk W Test not significant. F-Test followed by one-way, two-tailed unpaired Student's t-Test for equal variance | **t(10) = -2.756, p = 0.020** |
| **Fig. 4i** | Time at 50% OL gain (days) calculated from Mechanistic Growth Curves (Gray Matter vs. White Matter) | 26.8±2.2 vs. 24.3±4.4 days post-P70 | n = 6 mice | Shapiro-Wilk W Test not significant. F-Test followed by one-way, two-tailed unpaired Student's t-Test for equal variance | t(10) = -0.52, p = 0.615 |
| **Fig. 4j** | Two-week binned rates of oligodendrocyte gain (not modeled, % per week, GM vs. WM) | 1.0±0.4 vs. 2.5±0.5, WM, Weeks 5-6 vs. Weeks 1-2; 0.8±0.2 vs. 2.5±0.5, WM, Weeks 9-10 vs. Weeks 1-2 | n = 6 mice | Shapiro-Wilk W Test not significant. One-way ANOVA followed by Dunnett's test with control. | **p = 0.037 for Weeks 5-6 vs. Weeks 1-2; p = 0.022 for Weeks 9-10 vs. Weeks 1-2** |
| **Fig. 4n** | Change in percentage of MOL1-positive oligodendrocytes with aging in GM (%P140 - %P60) | -4.94±3.45% vs. normalized % at P60 | n = 6 mice (P60), n = 8 mice (P140), 2 sections per mouse | Shapiro-Wilk W Test not significant. F-Test followed by one-way, two-tailed unpaired Student's t-Test for equal variance | t(12) = -0.81, p = 0.434 |
| | Change in percentage of MOL2/3-positive oligodendrocytes with aging in GM (%P140 - %P60) | 0.07±0.51% vs. normalized % at P60 | n = 6 mice (P60), n = 8 mice (P140), 2 sections per mouse | Shapiro-Wilk W Test not significant. F-Test followed by one-way, two-tailed unpaired Student's t-Test for equal variance | t(12) = 0.10, p = 0.919 |
| | Change in percentage of MOL5/6-positive oligodendrocytes with aging in GM (%P140 - %P60) | -3.99±3.42% vs. normalized % at P60 | n = 6 mice (P60), n = 8 mice (P140), 2 sections per mouse | Shapiro-Wilk W Test not significant. F-Test followed by one-way, two-tailed unpaired Student's t-Test for equal variance | t(12) = 0.80, p = 0.440 |
| | Change in percentage of MOL1-positive oligodendrocytes with aging in WM (%P140 - %P60) | 0.19±0.17% vs. normalized % at P60 | n = 6 mice (P60), n = 8 mice (P140), 2 sections per mouse | Shapiro-Wilk W Test not significant. F-Test followed by one-way, two-tailed unpaired Student's t-Test for equal variance | t(12) = -0.67, p = 0.514 |
| | Change in percentage of MOL2/3-positive oligodendrocytes with aging in WM (%P140 - %P60) | 0.80±0.82% vs. normalized % at P60 | n = 6 mice (P60), n = 8 mice (P140), 2 sections per mouse | Shapiro-Wilk W Test significant. Wilcoxon rank sum test. | z = 1.36, p = 0.175 |
| | Change in percentage of MOL5/6-positive oligodendrocytes with aging in WM (%P140 - %P60) | 2.30±4.69% vs. normalized % at P60 | n = 6 mice (P60), n = 8 mice (P140), 2 sections per mouse | Shapiro-Wilk W Test not significant. F-Test followed by one-way, two-tailed unpaired Student's t-Test for equal variance | t(12) = 0.25, p = 0.805 |
| **Fig. 4o** | Percentage of MOBP-EGFP oligodendrocytes that are MOL1-positive at P140 | 17.2 ± 3.4 (GM) vs. 0.6 ± 0.2% (WM) | n = 6 mice (P60), n = 8 mice (P140), 2 sections per mouse | Shapiro-Wilk W Test not significant. F-Test followed by one-way, two-tailed unpaired Student's t-Test for unequal variance | **t(7.033) = -4.804, p = 0.002** |
| | Percentage of MOBP-EGFP oligodendrocytes that are MOL2/3-positive at P140 | 1.8 ± 0.5 (GM) vs. 3.8 ± 0.8% (WM) | n = 6 mice (P60), n = 8 mice (P140), 2 sections per mouse | Shapiro-Wilk W Test significant. Wilcoxon rank sum test. | **z = 2.363, p = 0.018** |
| | Percentage of MOBP-EGFP oligodendrocytes that are MOL5/6-positive at P140 | 27.4 ± 3.4 (GM) vs. 46.4 ± 4.7% (WM) | n = 6 mice (P60), n = 8 mice (P140), 2 sections per mouse | Shapiro-Wilk W Test not significant. F-Test followed by one-way, two-tailed unpaired Student's t-Test for equal variance | **t(14) = 3.283, p = 0.005** |

| Ext. Data Fig. 7 | Measure | Values | N | Statistical test | Significance |
|---|---|---|---|---|---|
| **Fig. 7c** | Density of OPCs in posterior parietal cortex (GM) vs. white matter (WM) | 181.2 ± 7.7 (GM) vs. 248.6 ± 23.8 (WM) $mm^2$ | n = 4 mice, 3 sections per mouse | Shapiro-Wilk W Test not significant. F-Test followed by one-way, two-tailed unpaired Student's t-Test for equal variance | **t(6) = 2.69, p = 0.036** |
| **Fig. 7d** | Density of EdU-positive OPCs in posterior parietal cortex (GM) vs. white matter (WM) | 14.6 ± 2.5 (GM) vs. 51.6 ± 6.6% (WM) | n = 4 mice, 3 sections per mouse | Shapiro-Wilk W Test not significant. F-Test followed by one-way, two-tailed unpaired Student's t-Test for equal variance | **t(6) = 5.29, p = 0.002** |
| **Fig. 7f** | % MOBP-EGFP oligodendrocytes positive for Egr2 (MOL1) in the spinal cord GM vs. WM | 18.61 ± 3.9 (GM) vs. 0.05 ± 0.02% (WM) | n = 3 mice, 2 spinal cord sections per mouse | Shapiro-Wilk W Test not significant. F-Test followed by one-way, two-tailed unpaired Student's t-Test for unequal variance | **t(2.00) = -4.68, p = 0.043** |
| | % MOBP-EGFP oligodendrocytes positive for Klk6 (MOL2/3) in the spinal cord GM vs. WM | 1.28 ± 0.26 (GM) vs. 37.23 ± 4.1% (WM) | n = 3 mice, 2 spinal cord sections per mouse | Shapiro-Wilk W Test not significant. F-Test followed by one-way, two-tailed unpaired Student's t-Test for unequal variance | **t(2.02) = 8.86, p = 0.012** |
| | % MOBP-EGFP oligodendrocytes positive for Ptgds (MOL5/6) in the spinal cord GM vs. WM | 32.27 ± 5.31 (GM) vs. 0.90 ± 0.50% (WM) | n = 3 mice, 2 spinal cord sections per mouse | Shapiro-Wilk W Test not significant. F-Test followed by one-way, two-tailed unpaired Student's t-Test for unequal variance | **t(2.03) = -5.88, p = 0.027** |

Supplementary Table 3 | Statistics results table

*Statistics results are sorted by figure, and both main figures and supplementary figures are listed in the order they appear in the text.*

**Statistically significant comparisons are bolded**

Table 3.4 | Statistics for Figure 5

| Fig. 5 | Measure | Values | N | Statistical test | Significance |
|---|---|---|---|---|---|
| **Fig. 5c** | % depth of corpus callosum analyzed longitudinally (Healthy vs. Cuprizone) | 84±8.0% (healthy) vs. 86.8±3.5% (cuprizone) | n=6 mice (Healthy), n = 6 mice (Cuprizone) | Shapiro-Wilk W Test significant. Wilcoxon rank sum test. | Z = 0.48, p = 0.629 |
| **Fig. 5h** | Total # lost oligodendrocytes / 350 x 350 x 60 µm volume at 66 day time point (Gray Matter vs. White Matter) | 36.8±6.0 vs. 157.4±37.8 OLs / imaging volume | n = 6 mice (Cuprizone) | Shapiro-Wilk W Test not significant. F-Test followed by one-way, two-tailed unpaired Student's t-Test for unequal variance | **t(5.25) = 3.15, p = 0.024** |
| **Fig. 5j** | Total % lost oligodendrocytes (Gray Matter vs. White Matter) at 66 day time point. | 75.3±6.3% (GM) vs. 75.6±7.4% (WM) | n = 6 mice (Cuprizone) | Shapiro-Wilk W Test not significant. F-Test followed by one-way, two-tailed unpaired Student's t-Test for equal variance | t(10) = 0.04, p = 0.970 |
| **Fig. 5k** | % Rate of Oligodendrocyte Loss during demyelination (Gray Matter vs. White Matter) calculated from Gompertz 3-parameter growth curves | 9.0±0.7% vs. 9.2±1.0% lost per week calculated from Gompertz 3-parameter modeling (Gray Matter vs. White Matter) | n = 6 mice (Cuprizone) | Shapiro-Wilk W Test not significant. F-Test followed by one-way, two-tailed unpaired Student's t-Test for equal variance | t(10) = 0.16, p = 0.876 |
| **Fig. 5l** | Inflection point of loss (days from end of cuprizone) calculated from Gompertz 3-parameter modeling (Gray Matter vs. White Matter) | 0.4±1.4 (GM) vs. 2.5±2.0 (WM) days post cuprizone | n = 6 mice (Cuprizone) | Shapiro-Wilk W Test significant. Wilcoxon rank sum test. | Z = 1.04, p = 0.298 |
| **Fig. 5m** | Rates of oligodendrocyte loss binned by 1-3 weeks relative to cuprizone administration (not modeled, % per week, GM vs. WM) | 9.6±1.8 vs. 3.7±0.8 % per week, GM, Weeks -2 to 0 vs. Week -3; 18.9±1.9 vs. 3.7±0.8 % per week, GM, Weeks 0 to 2 vs. Week -3; 22.6±3.1 vs. 4.1±2.5 % per week, WM, Weeks 1 to 2 vs. Week -3 | n = 6 mice (Cuprizone) | Shapiro-Wilk W Test significant. Kruskal-Wallis followed by Steel Method for nonparametric multiple comparisons with control. | **p = 0.046 for Weeks -2-0 vs. Weeks -3; p = 0.046 for Weeks 1 to 2 vs. Week -3** |
| **Fig. 5q** | Change in percentage of MOL1-positive oligodendrocytes with cuprizone demyelination in GM (%Cup. + 4d. - %P140 ) | -15.44±0.65% vs. normalized % at P140 | n = 8 mice (P140), n = 5 mice (Cuprizone + 4 days) | Shapiro-Wilk W Test significant. Wilcoxon rank sum test. | **Z = -2.71, p = 0.007** |
| | Change in percentage of MOL2/3-positive oligodendrocytes with cuprizone demyelination in GM (%Cup. + 4d. - %P140 ) | -1.76±0.05% vs. normalized % at P140 | n = 8 mice (P140), n = 5 mice (Cuprizone + 4 days) | Shapiro-Wilk W Test significant. Wilcoxon rank sum test. | **Z = -2.90, p = 0.004** |
| | Change in percentage of MOL5/6-positive oligodendrocytes with cuprizone demyelination in GM (%Cup. + 4d. - %P140 ) | -24.24±1.11% vs. normalized % at P140 | n = 8 mice (P140), n = 5 mice (Cuprizone + 4 days) | Shapiro-Wilk W Test not significant. F-Test followed by one-way, two-tailed unpaired Student's t-Test for unequal variance | **t(8.39) = -6.75, p =0.0001** |
| | Change in percentage of MOL1-positive oligodendrocytes with cuprizone demyelination in WM (%Cup. + 4d. - %P140 ) | -0.57±0.06% vs. normalized % at P140 | n = 8 mice (P140), n = 5 mice (Cuprizone + 4 days) | Shapiro-Wilk W Test significant. Wilcoxon rank sum test. | **Z = -2.07, p = 0.038** |
| | Change in percentage of MOL2/3-positive oligodendrocytes with cuprizone demyelination in WM (%Cup. + 4d. - %P140 ) | -2.58±0.52% vs. normalized % at P140 | n = 8 mice (P140), n = 5 mice (Cuprizone + 4 days) | Shapiro-Wilk W Test significant. Wilcoxon rank sum test. | **Z = -2.86, p = 0.004** |
| | Change in percentage of MOL5/6-positive oligodendrocytes with cuprizone demyelination in WM (%Cup. + 4d. - %P140 ) | -41.56±1.58% vs. normalized % at P140 | n = 8 mice (P140), n = 5 mice (Cuprizone + 4 days) | Shapiro-Wilk W Test not significant. F-Test followed by one-way, two-tailed unpaired Student's t-Test for unequal variance | **t(8.50) = -8.40, p <0.0001** |
| **Fig. 5r** | Percentage of MOBP-EGFP oligodendrocytes that are MOL1-positive 4 days post-cuprizone removal | 1.81 ± 0.65 (GM) vs. 0.09% ± 0.06% (WM) | n = 5 mice (Cuprizone + 4 days) | Shapiro-Wilk W Test significant. Wilcoxon rank sum test. | **Z = -2.54, p = 0.011** |
| | Percentage of MOBP-EGFP oligodendrocytes that are MOL2/3-positive 4 days post-cuprizone removal | 0.05 ± 0.04 (GM) vs. 1.2 ± 0.23% (WM) | n = 5 mice (Cuprizone + 4 days) | Shapiro-Wilk W Test significant. Wilcoxon rank sum test. | **Z = 2.59, p = 0.0097** |
| | Percentage of MOBP-EGFP oligodendrocytes that are MOL5/6-positive 4 days post-cuprizone removal | 3.13 ± 1.11 (GM) vs. 4.9 ± 1.6% (WM) | n = 5 mice (Cuprizone + 4 days) | Shapiro-Wilk W Test not significant. F-Test followed by one-way, two-tailed unpaired Student's t-Test for equal variance | t(8) = 0.89, p = 0.398 |

### Supplementary Table 3 | Statistics results table

*Statistics results are sorted by figure, and both main figures and supplementary figures are listed in the order they appear in the text.*

**Statistically significant comparisons are bolded**

#### Table 3.5 | Statistics for Figure 6

| Fig. 6 | Measure | Values | N | Statistical test | Significance |
|---|---|---|---|---|---|
| **Fig. 6e** | Total # new oligodendrocytes / 350 x 350 x 60 µm volume after Cuprizone at 66 day time point (Gray Matter vs. White Matter) | 13.3±2.0 (GM) vs. 123±47.3 (WM) OLs/imaging volume | n = 6 mice (cuprizone) | Shapiro-Wilk W Test significant. Wilcoxon rank sum test. | **Z = 2.80, p = 0.005** |
| **Fig. 6g** | Total % Replacement after Cuprizone (% Gain normalized to % Lost) at 66 day time point (Gray Matter vs. White Matter) | 37.6±3.6% (GM) vs. 68.0±11.3% (WM) | n = 6 mice (cuprizone) | Shapiro-Wilk W Test not significant. F-Test followed by one-way, two-tailed unpaired Student's t-Test for unequal variance | **t(5.97) = 2.57, p = 0.043** |
| **Fig. 6h** | % Replacement Rate during remyelination (Gray Matter vs. White Matter) calculated from Gompertz 3-parameter growth curves | 5.6±0.6% (GM) vs. 11.1±1.9% (WM) | n = 6 mice (cuprizone) | Shapiro-Wilk W Test not significant. F-Test followed by one-way, two-tailed unpaired Student's t-Test for unequal variance | **t(5.72) = 2.59 p = 0.043** |
| **Fig. 6i** | Inflection point of Cumulative % Replacement (days from end of cuprizone) calculated from Gompertz 3-parameter modeling (Gray Matter vs. White Matter) | 15.352±0.77 (GM) vs. 15.358±1.56 Days post-cuprizone (WM) | n = 6 mice (cuprizone) | Shapiro-Wilk W Test not significant. F-Test followed by one-way, two-tailed unpaired Student's t-Test for equal variance | t(10) = 0.0037, p = 0.997 |
| **Fig. 6j** | Rates of oligodendrocyte gain binned by 1-3 weeks relative to cuprizone administration (not modeled, % per week, GM vs. WM) | 8.2±1.1 vs. 3.0±0.5% per week, GM, Weeks 3-4 vs. Week 7; 18.1±5.1 vs. 4.5±1.3% per week, WM, Weeks 3-4 vs. Week 7 | n = 6 mice (cuprizone) | Shapiro-Wilk W Test not significant. One-way ANOVA followed by Dunnett's method for comparison with control | **p = 0.002 for Weeks 3-4 vs. Week 7 in the GM; p = 0.0052 for Weeks 3-4 vs. Week 7 in the WM** |
| | Peak oligodendrocyte gain rate (3-4 weeks post-cuprizone) | 8.2±1.1% (GM) vs. 18.1±5.1 (WM) | | Two-way ANOVA followed by piecewise Student's t comparison with Bonferroni correction for multiple comparisons | **p = 0.0009; Bonferroni-corrected alpha = 0.0125** |
| **Fig. 6n** | Change in percentage of MOL1-positive oligodendrocytes with cuprizone demyelination in GM ( %Cup. + 7w. - %P140 ) | -1.50±3.36% vs. normalized % at P140 | n = 8 mice (P140), n = 7 mice (Cup. + 7 weeks) | Shapiro-Wilk W Test not significant. F-Test followed by one-way, two-tailed unpaired Student's t-Test for equal variance | t(13) = -0.31, p = 0.763 |
| | Change in percentage of MOL2/3-positive oligodendrocytes with cuprizone demyelination in GM ( %Cup. + 7w. - %P140 ) | -0.58±0.44% vs. normalized % at P140 | n = 8 mice (P140), n = 7 mice (Cup. + 7 weeks) | Shapiro-Wilk W Test not significant. F-Test followed by one-way, two-tailed unpaired Student's t-Test for equal variance | t(13) = -0.86, p = 0.407 |
| | Change in percentage of MOL5/6-positive oligodendrocytes with cuprizone demyelination in GM ( %Cup. + 7w. - %P140 ) | -14.9±2.0% vs. normalized % at P140 | n = 8 mice (P140), n = 7 mice (Cup. + 7 weeks) | Shapiro-Wilk W Test not significant. F-Test followed by one-way, two-tailed unpaired Student's t-Test for unequal variance | **t(13) = -3.712, p = 0.003** |
| | Change in percentage of MOL1-positive oligodendrocytes with cuprizone demyelination in WM ( %Cup. + 7w. - %P140 ) | 0.80±0.33% vs. normalized % at P140 | n = 8 mice (P140), n = 7 mice (Cup. + 7 weeks) | Shapiro-Wilk W Test significant. Wilcoxon rank sum test. | z = 1.85, p = 0.064 |
| | Change in percentage of MOL2/3-positive oligodendrocytes with cuprizone demyelination in WM ( %Cup. + 7w. - %P140 ) | 0.31±1.22% vs. normalized % at P140 | n = 8 mice (P140), n = 7 mice (Cup. + 7 weeks) | Shapiro-Wilk W Test not significant. F-Test followed by one-way, two-tailed unpaired Student's t-Test for equal variance | t(13) = 0.22, p = 0.833 |
| | Change in percentage of MOL5/6-positive oligodendrocytes with cuprizone demyelination in WM ( %Cup. + 7w. - %P140 ) | -22.6±4.2% vs. normalized % at P140 | n = 8 mice (P140), n = 7 mice (Cup. + 7 weeks) | Shapiro-Wilk W Test significant. Wilcoxon rank sum test. | **z = -2.720, p = 0.007** |
| **Fig. 6o** | Percentage of MOBP-EGFP oligodendrocytes that are MOL1-positive 7 weeks post-cuprizone removal | 15.8±3.3 (GM) vs. 1.5±0.3% (WM) | n = 8 mice (P140), n = 7 mice (Cup. + 7 weeks) | Shapiro-Wilk W Test not significant. F-Test followed by one-way, two-tailed unpaired Student's t-Test for unequal variance | **t(6.118) = -4.234, p = 0.005** |
| | Percentage of MOBP-EGFP oligodendrocytes that are MOL2/3-positive 7 weeks post-cuprizone removal | 1.2±0.4 (GM) vs. 4.1±1.2% (WM) | n = 8 mice (P140), n = 7 mice (Cup. + 7 weeks) | Shapiro-Wilk W Test not significant. F-Test followed by one-way, two-tailed unpaired Student's t-Test for unequal variance | **t(7.560) = 2.187, p = 0.063** |
| | Percentage of MOBP-EGFP oligodendrocytes that are MOL5/6-positive 7 weeks post-cuprizone removal | 12.6±2.0 (GM) vs. 24.0±4.2% (WM) | n = 8 mice (P140), n = 7 mice (Cup. + 7 weeks) | Shapiro-Wilk W Test not significant. F-Test followed by one-way, two-tailed unpaired Student's t-Test for equal variance | **t(12) = 2.432, p = 0.032** |

Supplementary Table 3 | Statistics results table

*Statistics results are sorted by figure, and both main figures and supplementary figures are listed in the order they appear in the text.*

**Statistically significant comparisons are bolded**

Table 3.6: Statistics for Fig. 7

| Fig. 7 | Measure | Values | N | Statistical test | Significance |
|---|---|---|---|---|---|
| **Fig. 7d** | % new oligodendrocyte gain in the healthy brain per week by layer | 2.6±0.2 (L4) vs. 1.4±0.2% (CC) | n = 6 healthy mice | Shapiro-Wilk W Test not significant. One-way ANOVA followed by Tukey's HSD | **p = 0.012** |
| **Fig. 7e** | Percentage of MOBP-EGFP oligodendrocytes that are MOL1-positive at P140 by layer | 23.44±3.54% (L1-3) vs. 18.27±4.50% (L4) vs. 13.84±3.88% (L5-6) vs. 0.66±0.16% (CC) | n = 8 mice (P140) | Shapiro-Wilk W Test not significant. One-way ANOVA followed by Tukey's HSD | **p = 0.0004, L1-3 vs. CC; p = 0.006, L4 vs. CC** |
| **Fig. 7f** | Percentage of MOBP-EGFP oligodendrocytes that are MOL2/3-positive at P140 by layer | 0% (L1-3) vs. 0% (L4) vs. 2.98±0.78% (L5-6) vs. 3.75±0.0.82% (CC) | n = 8 mice (P140) | Shapiro-Wilk W Test significant. Kruskal-Wallis test followed by Dunn's test for multiple comparisons | **p = 0.0034 (L4 vs. L5-6); p = 0.0034 (L1-3 vs. L5-6); p = 0.0009 (L4 vs. CC); p = 0.0009 (L5-6 vs. CC)** |
| **Fig. 7g** | Percentage of MOBP-EGFP oligodendrocytes that are MOL5/6-positive at P140 by layer | 21.674±4.12% (L1-3) vs. 26.38±3.1% (L4) vs. 31.26±3.09% (L5-6) vs. 46.42±4.69% (CC) | n = 8 mice (P140) | Shapiro-Wilk W Test not significant. One-way ANOVA followed by Tukey's HSD | **p = 0.042 (L5-6 vs. CC); p = 0.005 (L4 vs. CC); p = 0.0005 (L1-3 vs. CC)** |
| **Fig. 7i** | % cuprizone-induced oligodendrocyte loss per week by layer | 1.12±0.26 (L1-3) vs. 1.52±0.32(L4) vs. 2.106±0.12% (L5-6) vs. 1.88±0.23 (CC) | n = 6 cuprizone mice | Shapiro-Wilk W Test not significant. One-way ANOVA. | Prob > F = 0.0504 |
| **Fig. 7j** | Percentage of MOBP-EGFP oligodendrocytes that are MOL1-positive at 4 days post-cuprizone by layer | 7.56±2.88% (L1-3) vs. 1.23±0.67% (L4) vs. 0.31±0.17% (L5-6) vs. 0.09±0.06% (CC) | n = 5 mice (Cuprizone + 4 days) | Shapiro-Wilk W Test significant. Kruskal-Wallis test followed by Dunn's test for multiple comparisons | **p = 0.016, L1-3 vs. CC** |
| **Fig. 7k** | Percentage of MOBP-EGFP oligodendrocytes that are MOL2/3-positive at 4 days post-cuprizone by layer | 0% (L1-3) vs. 0% (L4) vs. 0.09±0.09% (L5-6) vs. 01.18±0.23% (CC) | n = 5 mice (Cuprizone + 4 days) | Shapiro-Wilk W Test significant. Kruskal-Wallis test followed by Dunn's test for multiple comparisons | **p = 0.022 L5-6 vs. CC; p = 0.004 L4 vs. CC; p = 0.004 L1-3 vs. CC** |
| **Fig. 7l** | Percentage of MOBP-EGFP oligodendrocytes that are MOL5/6-positive at 4 days post-cuprizone by layer | 6.65±2.28% (L1-3) vs. 4.41±01.33% (L4) vs. 1.90±0.82% (L5-6) vs. 4.85±1.59% (CC) | n = 5 mice (Cuprizone + 4 days) | Shapiro-Wilk W Test not significant. One-way ANOVA. | Prob > F = 0.250 |
| **Fig. 7n** | Oligodendrocyte replacement rate during remyelination (% of lost cells replaced per week) by layer | 6.04±0.93% (L1-3) vs. 7.32±1.46% (L4) vs. 5.58±0.56% (L5-6) vs. 11.15±1.93% (CC) | n = 6 cuprizone mice | Shapiro-Wilk W Test not significant. One-way ANOVA followed by Tukey's HSD | **p = 0.035 (L5-6 vs. CC)** |
| **Fig. 7o** | Percentage of MOBP-EGFP oligodendrocytes that are MOL1-positive at 7 weeks post-cuprizone by layer | 16.34±4.63% (L1-3) vs. 20.60±5.47% (L4) vs. 13.66±2.91% (L5-6) vs. 1.46±0.33% (CC) | n = 7 mice (Cuprizone + 7 weeks) | Shapiro-Wilk W Test not significant. One-way ANOVA followed by Tukey's HSD | **p = 0.012 (L4 vs. CC)** |
| **Fig. 7p** | Percentage of MOBP-EGFP oligodendrocytes that are MOL2/3-positive at 7 weeks post-cuprizone by layer | 0% (L1-3) vs. 0% (L4) vs. 2.33±0.97% (L5-6) vs. 4.06±1.22% (CC) | n = 7 mice (Cuprizone + 7 weeks) | Shapiro-Wilk W Test significant. Kruskal-Wallis test followed by Dunn's test for multiple comparisons | **p = 0.025 (L1-3 vs. L5-6); p = 0.025 (L4 vs. L5-6); p = 0.003 (L1-3 vs. CC); p = 0.003 (L4 vs. CC)** |
| **Fig. 7q** | Percentage of MOBP-EGFP oligodendrocytes that are MOL5/6-positive at 7 weeks post-cuprizone by layer | 13.87±1.86% (L1-3) vs. 19.67±2.57% (L4) vs. 10.58±2.17% (L5-6) vs. 23.98±4.24% (CC) | n = 7 mice (Cuprizone + 7 weeks) | Shapiro-Wilk W Test not significant. One-way ANOVA followed by Tukey's HSD | **p = 0.015 (L5-6 vs. CC)** |

Supplementary Table 3 | Statistics results table

*Statistics results are sorted by figure, and both main figures and supplementary figures are listed in the order they appear in the text.*

**Statistically significant comparisons are bolded**

Table 3.7: Statistics for Fig. 8 and associated extended data (Extended Data Fig. 10)

| Fig. 8 | Measure | Values | N | Statistical test | Significance |
|---|---|---|---|---|---|
| **Fig. 8b** | Scaled Total % Population Gain (Cuprizone vs. Healthy) | 0.33±0.08 (cuprizone) vs. 0.57±0.14 (healthy) | n = 6 mice (healthy), n = 6 mice (cuprizone) | Shapiro-Wilk W Test not significant. Two-way ANOVA followed by piecewise Student's t comparison with Bonferroni correction for multiple comparisons | **p = 0.006, Bonferroni-corrected alpha = 0.0125** |
| **Fig. 8c** | Scaled % Gain Rate (Cuprizone vs. Healthy) | 0.30±0.03 (cuprizone) vs. 0.57±0.06 (healthy) | n = 6 mice (healthy), n = 6 mice (cuprizone) | Shapiro-Wilk W Test not significant. Two-way ANOVA followed by piecewise Student's t comparison with Bonferroni correction for multiple comparisons | **p = 0.0025, Bonferroni-corrected alpha = 0.0125** |
| **Fig. 8d** | Scaled Inflection point of % Cumulative Gain or Replacement (days from end of cuprizone) calculated from Gompertz 3-parameter modeling (Cuprizone vs. Healthy) | Scaled inflection points across treatment and layers = 0.67±0.02 | n = 6 mice (healthy), n = 6 mice (cuprizone) | Shapiro-Wilk W Test not significant. Two-way ANOVA no significant interaction. | Prob > F = 0.284 |
| **Fig. 8e** | # of MOL subtypes present at each layer across healthy, Cup. + 4d., and Cup. + 7 weeks groups | L1-3: 2.0±0.0 classes (Healthy P140); 0.8±0.49 (Cup. + 4d.); 1.71±0.18 (Cup. + 7wk.)<br><br>L4: 1.88±0.13 classes (Healthy P140); 0.4-±0.24 (Cup. + 4d.); 1.71±0.18 (Cup. + 7wk.)<br><br>L5-6: 2.0±0.0 classes (Healthy P140); 1.4±0.25 (Cup. + 4d.); 1.71±0.29 (Cup. + 7wk.)<br><br>CC: 2.0±0.0 classes (Healthy P140); 1.4±0.25 (Cup. + 4d.); 1.71±0.29 (Cup. + 7wk.) | n = 8 mice (P140), n = 5 mice (Cup. + 4d.), n = 7 mice (Cup. + 7 weeks) | Shapiro-Wilk W Test significant. Kruskal-Wallis test followed by Dunn's test for multiple comparisons | **L1-3: Prob > ChiSq = 0.036; p = 0.021 (Healthy vs. Cup + 4d.)**<br><br>**L4: Prob > ChiSq = 0.002; p = 0.003 (Healthy vs. Cup + 4d.); p = 0.019 (Cup. + 7wks. vs. Cup. + 4d.)**<br><br>**L5-6: Prob > ChiSq = 0.002; p = 0.005 (Healthy vs. Cup + 4d.); p = 0.009 (Cup. + 7wks. vs. Cup. + 4d.)**<br><br>CC: Prob > ChiSq = 0.051 |
| **Fig. 8f** | Percentage of MOBP-EGFP oligodendrocytes that are MOL1-positive across healthy, Cup. + 4d., and Cup. + 7 weeks groups | Percentage MOL1-positive oligodendrocytes across time point, treatment, and layers = | n = 8 mice (P140), n = 5 mice (Cup. + 4d.), n = 7 mice (Cup. + 7 weeks) | Two-way ANOVA no significant interaction. | Prob > F = 0.1397 |
| **Fig. 8g** | Percentage of MOBP-EGFP oligodendrocytes that are MOL2/3-positive across healthy, Cup. + 4d., and Cup. + 7 weeks groups | Percentage MOL1-positive oligodendrocytes across time point, treatment, and layers = | n = 8 mice (P140), n = 5 mice (Cup. + 4d.), n = 7 mice (Cup. + 7 weeks) | Two-way ANOVA no significant interaction. | Prob > F = 0.105 |
| **Fig. 8h** | Percentage of MOBP-EGFP oligodendrocytes that are MOL5/6-positive across healthy, Cup. + 4d., and Cup. + 7 weeks groups | L4: 26.4±3.11 (Healthy P140) vs. 4.4±1.33 (Cup. + 4d.)<br><br>L5-6: 31.3±3.09 (Healthy P140) vs. 1.9±0.82 (Cup. + 4d.); vs. 10.6±2.17 (Cup. + 7w.)<br><br>CC: 46.4±4.69 (Healthy P140) vs. 4.9±1.58 (Cup. + 4d.) vs. 24.0±4.24 (Cup. + 7w.) | n = 8 mice (P140), n = 5 mice (Cup. + 4d.), n = 7 mice (Cup. + 7 weeks) | Shapiro-Wilk W Test not significant. Two-way ANOVA followed by Tukey's HSD. | **Prob > F = 0.004 (Condition x Layer)**<br><br>**L4: p = 0.001 (Healthy P140 vs. Cup. + 4d.)**<br><br>**L5/6: p <0.0001 (Healthy P140 vs. Cup. + 4d.); p <0.0007 (Healthy P140 vs. Cup. + 7wk.);** p = 0.834 (Cup. + 4d. Vs. Cup. + 7wk.)<br><br>**CC: p < 0.0001 (Healthy P140 vs. Cup. + 4d.), p = 0.0001 (Healthy P140 vs. Cup. + 7wk.), p = 0.012 (Cup. + 4d. Vs. Cup. + 7wk.)** |

| Ext. Data Fig. 10 | Measure | Values | N | Statistical test | Significance |
|---|---|---|---|---|---|
| **Fig. 10a** | Oligodendrocyte replacement rate (% per week) across one week time bins (unmodeled) | L1-3: 2.04±0.28 (healthy); 7.59±1.27 (Week 2); 7.69±1.06 (Week 3); 6.91±1.19 (Week 4)<br><br>L4: 2.62±0.29 (healthy); 10.96±2.24 (Week 3); 9.35±1.70 (Week 4)<br><br>L5-6: 2.11±0.24 (healthy); 11.13±2.02 (Week 3); 6.56±1.17 (Week 4)<br><br>CC: 21.42±0.21 (healthy); 18.98±6.17 (Week 2); 25.99±9.04 (Week 3); 10.68±1.13 (Week 4) | n = 6 mice (healthy), n = 6 mice (cuprizone) | Shapiro-Wilk W Test significant. Kruskal-Wallis followed by Steel Method for nonparametric multiple comparisons with control. | **p = 0.034 for L1-3, Weeks 2-4 vs. Healthy; p = 0.034 for L4 Weeks 3-4 vs. Healthy; p = 0.034 for L5-6, Weeks 3-4 vs. Healthy; p = 0.034 for CC Weeks 2-4 vs. Healthy** |
| **Fig. 10c** | Full-width at half-maximum of the oligodendrocyte replacement response curve in (10b) by layer (days) | 19.89±2.14 (L1-3) vs. 11.70±2.37 (CC) | n = 6 mice (healthy), n = 6 mice (cuprizone) | Shapiro-Wilk W Test not significant. One-way ANOVA followed by Tukey's HSD | **p = 0.048** |
| **Fig. 10d** | Area under the curve (after subtraction of the healthy gain curve) of the oligodendrocyte replacement response in (10b) by layer | 22.8±2.9 (L5-6) vs. 56.2±12.1% (CC) | n = 6 mice (healthy), n = 6 mice (cuprizone) | Shapiro-Wilk W Test not significant. One-way ANOVA followed by Tukey's HSD | **p = 0.037** |
| **Fig. 10e** | Full-width at half-maximum of the oligodendrocyte loss response by layer | 15.10±2.23 (L1-3) vs. 13.04±2.08 (L4) vs. 12.06±1.13 (L5-6) vs. 10.64±1.91 (CC) | n = 6 mice (healthy), n = 6 mice (cuprizone) | Shapiro-Wilk W Test significant. Kruskal-Wallis test. | Prob > ChiSq = 0.515 |
| **Fig. 10f** | Area under the curve (after subtraction of the healthy gain curve) of the oligodendrocyte loss response by layer | 51.32±12.07 (L1-3) vs. 60.31±12.31 (L4) vs. 85.49±4.39 (L5-6) vs. 73.86±6.51 (CC) | n = 6 mice (healthy), n = 6 mice (cuprizone) | Shapiro-Wilk W Test not significant. One-way ANOVA | Prob > F = 0.087 |
